# Independent colonisations of serpentine habitats highlight species-specific evolutionary histories of lineage diversification

**DOI:** 10.1101/2025.11.04.686350

**Authors:** Pablo Arrufat, Noelia Hidalgo-Triana, Nazaret Keen, Jaime Francisco Pereña-Ortiz, Andrés V. Pérez-Latorre, Amaia Leunda-Esnaola, Alexandra García-Flórez, Vladimir R. Kaberdin, Filip Kolář, David López-Idiáquez, Peter B. Pearman

## Abstract

Serpentine soils are characterized by high levels of heavy metals, low nutrient availability, and water scarcity, presenting significant ecological challenges for plant species. Nonetheless, some species have adapted successfully to these conditions. We investigate the population genomic structure, evolutionary history and phenotypic differentiation of three diploid generalist plant species, *Lavandula stoechas* L., *Halimium atriplicifolium* (Lam) Spach. subsp. *atriplicifolium*, and *Phlomis purpurea* L., all of which inhabit adjacent serpentine and non-serpentine soils in the Málaga region in the southern Iberian Peninsula (Spain). We explore whether populations from serpentine and non-serpentine soils represent distinct evolutionary lineages and whether there is genomic and phenotypic differentiation associated with serpentine conditions. We measured plant height and specific leaf area (SLA) to detect potential ecotypic variation associated with soil type. A ddRADseq SNP dataset was generated for each species, representing 10 populations from serpentine and 10 from non-serpentine soils. We ordinated genotype data to assess genomic variation, conducted an ADMIXTURE analysis to infer ancestral groups, and used Treemix analyses to investigate phylogenetic relationships and gene flow events between populations. Isolation by distance (IBD) analyses evaluated the role of geographic separation in observed genomic differentiation. Our results reveal species-specific patterns of genomic, and to some extent phenotypic, differentiation between serpentine and non-serpentine populations, with evidence of multiple colonisations of serpentine sites in all three species. The study highlights the role of historical differentiation and subsequent gene flow in shaping the genomic structure of plant populations, alongside observed variation in phenotypic traits across environments.

## Introduction

A central goal of evolutionary biology is to understand how the geographical and ecological distributions of species change over time and what historical processes underlie those shifts (Avise, 2000). Such dynamics often involve colonization of new environments, and persistence under challenging ecological conditions (Szűcs et al., 2017). For example, climatic oscillations, landscape fragmentation, and heterogeneous selective regimes can leave lasting imprints on evolutionary history and population structure of species (Fenderson et al., 2020). The corresponding genomic and demographic patterns provide insights into how species have responded to past environmental challenges and interactions, including range expansions, contractions, and occupation of extreme habitats (Lee & Mitchell-Olds, 2011). Inferring the processes that likely shape biodiversity across spatial and temporal scales, as well as across ecological boundaries, can be supported by integrating genomic data with the ecological and demographic context of populations (Ellegren & Galtier, 2016; Savolainen et al., 2013). Such understanding is essential not only for advancing evolutionary theory but also for the development and implementation of conservation strategies in the face of ongoing environmental change (Cook & Sgrò, 2019).

The colonization of novel and heterogeneous environments often drives the emergence of ecotypes, which constitute genetically and ecologically distinct lineages within a species (Millien et al, 2006). Genomic analyses allow disentangling whether such divergence arises from a single ancestral component or through repeated, independent evolutionary events (James et al., 2023). In some cases, ecotypes form monophyletic clades, reflecting a single-origin divergence followed by ecological differentiation. This pattern is observed in marine *Prochlorococcus* (Cyanobacteria) populations, where high-light and low-light lineages represent distinct clades that emerged during colonization of different ocean depths (Rocap et al., 2003). A similar case occurs in Atlantic cod (*Gadus morhua*), where resident uord and migratory North Sea ecotypes form genomically coherent clades that are maintained across spatial scales, indicating divergence from a single evolutionary origin (Knutsen et al., 2018). Conversely, repeated, independent colonization of similar habitats can also produce ecotypic divergence. For instance, pelagic and coastal ecotypes of bottlenose dolphins (*Tursiops truncatus*) have evolved in parallel across multiple ocean basins, with genome-wide analyses revealing independent origins from standing genetic variation in multiple regions (Louis et al., 2020). Similarly, genomic analyses reveal the repeated arising of coastal ecotypes in the wildflower *Senecio lautus*, in which coastal forms have evolved independently across multiple lineages, and parallel divergence has occurred despite gene flow (Roda et al., 2013). It remains unclear whether parallel diversification to notably harsh abiotic environments with steep environmental gradients is a common phenomenon.

Just as ecotypic divergence can result from colonization of novel habitats across broad geographic and ecological gradients, exposure to distinct edaphic environments can lead to the emergence of genomically distinct plant populations (Wang et al., 2017). This process can give rise to populations specialized for particular soil conditions, thereby shaping patterns of genomic differentiation and demographic history (Ke et al., 2022). Since most terrestrial vascular plants are rooted in place, they are continuously exposed to the soil they inhabit, potentially making edaphic components a strong selective agent structuring plant lineages (Hulshof & Spasojevic, 2020). Among notable edaphic qualities, serpentine soils, which are characterized by low nutrient availability, high heavy metal content, and water scarcity, present a physiologically stressful environment for plants (Kruckeberg, 1985; Anacker, 2014), and impose strong ecological constraints that can drive phenotypic variation and local adaptation (Konečná et al., 2020). Consequently, serpentine ecotypes typically show reduced stature and leaf size combined with more extensive root systems compared to conspecifics on other soils (Brady et al., 2005). Despite their proximity to non-serpentine habitats, serpentine-tolerant populations often maintain distinct genomic identities (Sambatti & Rice, 2006) with various barriers to gene flow. These can include shifts in flowering time (Hidalgo-Triana & Pérez-Latorre, 2019) and reproductive incompatibilities that reinforce divergence even when populations on differing substrates are in close proximity (Moyle et al., 2012). The fragmented, isolated and replicated distribution of serpentine outcrops provides a suitable scenario for determining whether populations on serpentine soil typically arise from a single evolutionary event or from repeated, independent colonisations by multiple lineages (Konečná et al., 2020).

So far, studies analysing the evolutionary histories of plant species inhabiting serpentine soils have found that divergences have frequently arisen through parallel evolution, where distinct lineages colonize serpentine habitats independently, and populations converge on similar morphological and physiological traits. For example, independent origins of serpentine tolerance have occurred in *Mimulus guttatus* (Selby et al., 2014), *Cerastium alpinum* (Berglund et al., 2004), and *Arabidopsis arenosa* (Konečná et al., 2021), demonstrating that serpentine environments can promote recurrent and predictable evolutionary trajectories. However, most studies have focused on single species in specific regions, particularly in North America and Central Europe (Patterson & Givnish, 2004; Konečná et al., 2020), limiting our ability to generalize about the prevalence of parallel evolutionary processes across diverse non-model species in other regions.

Here, we aim to address the pervasiveness of parallel evolution to serpentine habitat by analysing three plant species from southern Spain: *Halimium atriplicifolium*, *Lavandula stoechas*, and *Phlomis purpurea*. Specifically, we ask whether serpentine tolerant populations of these species evolved independently, share patterns of genomic differentiation, and exhibit convergent phenotypic functional traits. To test these questions, we employed ddRAD sequencing combined with multiple tools for genomic analysis tools to decipher population structure, reconstruct evolutionary history, and detect potential gene flow among populations (Esposito et al., 2020). We also tested for isolation by distance (IBD) to evaluate the influence of geographical separation on population genomic differentiation (Bockelmann et al., 2003), and measured the phenotypic functional traits plant height and specific leaf area (SLA), to detect potential differences between serpentine and non-serpentine populations (Samojedny et al., 2022; Brady et al., 2005). We predict multiple independent colonisations of serpentine soils in all three species, and species-specific population structures and evolutionary histories, based on the diverse life histories, dispersal capacities, and ecological strategies of the species. This multispecies, multipopulation framework enables the disentangling of shared versus lineage-specific responses, and the assessment of the extent to which similar environmental stressors produce parallel genomic and phenotypic outcomes, both within and across species facing a strong edaphic gradient.

## Materials and Methods

### Study system

We studied three diploid plant species: *Halimium atriplicifolium*, *Lavandula stoechas* and *Phlomis purpurea*, all of which occur in the southern Iberian Peninsula. This region is a hotspot for biodiversity on serpentine soils and is characterized by numerous spatially discrete serpentine outcrops, the largest of which is found on the Sierra Bermeja massif (Pérez-Latorre et al, 2024). The presence of populations of the same species on both serpentine and non-serpentine soils, separated by a few hundred meters, provides an opportunity to investigate the genomic basis of the origins of edaphic habitation and tolerance. *H. atriplicifolium* is an evergreen shrub common in dry, nutrient-poor Mediterranean scrublands (Castro-Díez et al., 2005). *L. stoechas* is an aromatic subshrub that grows on thin, rocky soil and sun-exposed habitats (Bella et al, 2015), while *P. purpurea* is a robust perennial shrub, typically found in dry, rocky habitats in scrublands (Neves et al., 2023).

### Sampling design and collections

Plant leaf tissue was collected through targeted sampling in spring over three years (2022-2024). We sampled 20 populations that were chosen to form 10 pairs, each pair having one population on serpentine soil, and one on other soil (Fig. 1). Sampling was carried out in the province of Málaga, where the largest serpentine area on the Iberian Peninsula is found. We identified candidate population locations of the three species using georeferenced records from the Global Biodiversity Information Facility (GBIF.org, 2022), and integrating the field knowledge of NH-T, JFP-O, and AVP-L. We sampled 15 individual plants per population along an informal transect, oriented through the longest dimension of the population and starting with a plant at an arbitrary population edge. We sampled subsequent plants that were at least 2 m apart, except in two cases where the populations were so small as to make this infeasible. In these cases, neighbours were not collected whenever possible to reduce the collection of siblings and better span the extent of the population. For each individual plant, we collected approximately 1 g of fresh green leaves into manila coin envelopes, sealed these, and placed them immediately into tripled plastic zip-lock bags with silica bead desiccant to maintain DNA integrity. At each site, we collected six soil subsamples along the transect and combined them into a single sample. The chemical analysis of these soil samples was conducted by Eurofins Spain and used to confirm whether or not initial assignation to serpentine or non-serpentine soil type was correct (see supplementary information (SI)-1, for further information).

**Fig 1.**
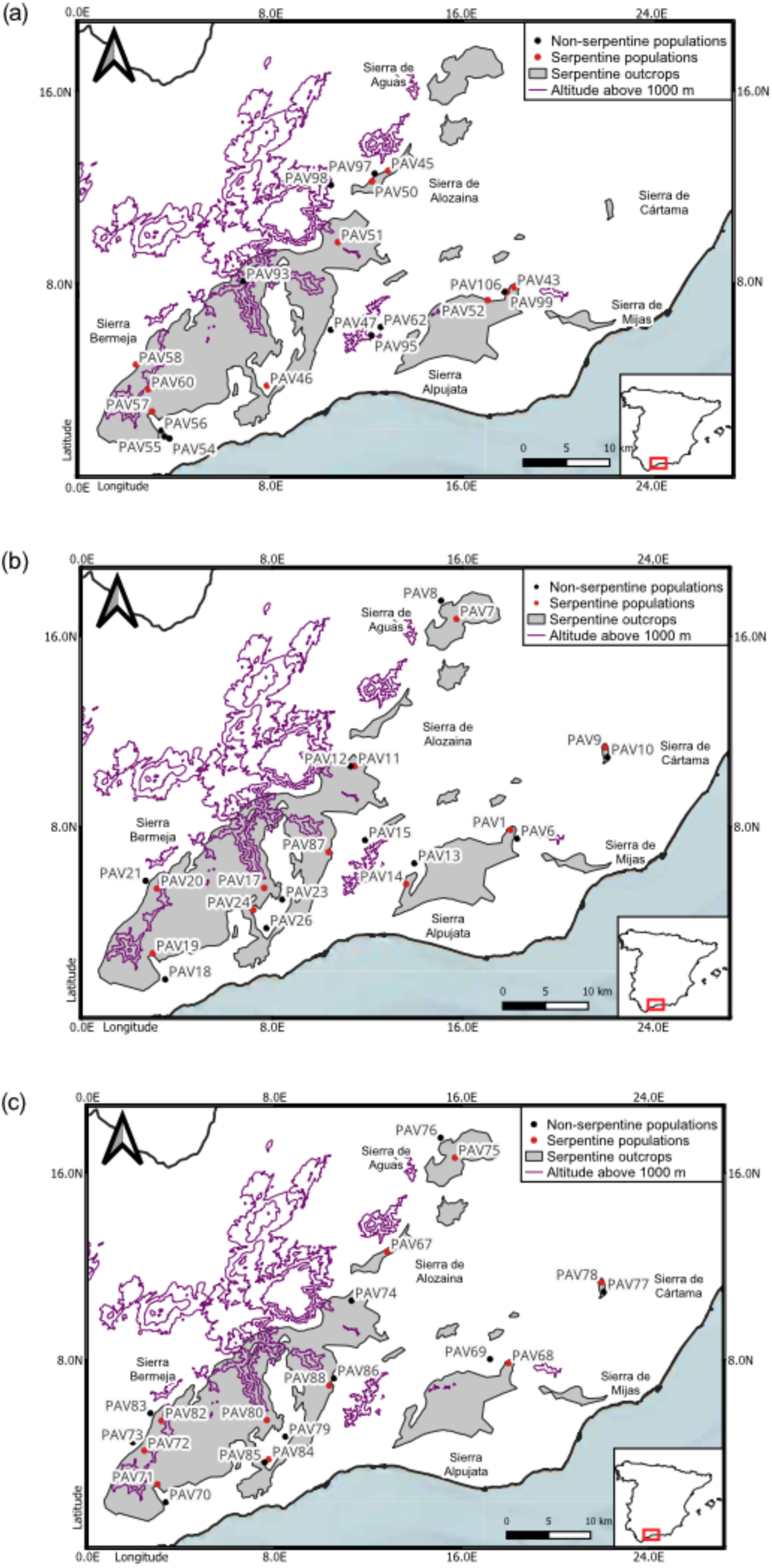
Locations of the sampled populations. Each dot represents a population from which 15 individual plants were collected for analysis. (a) *Halimium atriplicifolium*, (b) *Lavandula stoechas* and, (c) *Phlomis purpurea*.

### Functional traits

We tested for ecotypic variation between serpentine and non-serpentine soils using two functional traits: plant height and specific leaf area (SLA), which were measured following standardized procedures (Cornelissen et al., 2003; Perez-Harguindeguy 2016). We measured plant height using a measuring tape as the distance from the base of the stem to the highest living part of the plant (excluding inflorescences). We measured SLA (defined as one-sided area of a fresh leaf divided by leaf dry mass) during subsequent years in different field campaigns for logistical reasons. Although all sampled individuals were marked with metal tags during the first campaign, not all tagged individuals could be relocated in later years (557 out of 885 were found). Consequently, sample sizes for SLA analyses differed among populations. We collected fully developed, undamaged adult leaves from the upper part of each relocated plant.

We placed each leaf in a sealed plastic bag, and then all leaves from the same population were stored together in a second bag. The samples were kept cool until they were processed in the laboratory. Fresh leaves were first scanned using an Epson Perfection V600 Photo® device, with the program ScanWizard Pro (Microtek International, Inc, Hsinchu City, Taiwan). We then measured leaf area from the digital images with ImageJ (NIH, Maryland, USA). We determined leaf dry mass by wrapping leaves separately in aluminium foil and heating them at 60 °C in a drying oven for 72 hours. We immediately measured dry mass with a microbalance. Leaf SLA was then calculated for each individual as the ratio of leaf area to dry mass. We used linear mixed-effects models to assess differences in SLA between soil types, with soil type (serpentine vs. non-serpentine) as a fixed effect and population as a random effect, in order to account for the non-independence of samples collected from individuals in the same population. We modelled plant height and SLA separately for each species using customized scripts in R (R Core Team, 2023).

### DNA isolation and Sequencing

Approximately 30 milligrams of the silica-dried, young fresh leaves were ground with a hand pestle to a fine powder in a mortar refrigerated with underlying liquid nitrogen. After trial extractions with several commercial kits, whole genomic DNA was isolated from 14 individuals per population with the DNeasy® Plant Mini Kit (Qiagen, Hilden, Germany) for *H. atriplicifolium* and *P. purpurea*. For *L. stoechas* leaves, we used the Nucleospin Plant II Kit (Macherey-Nagel, Duren, Germany) because of improved purity of extracted DNA. Each extracted DNA sample was evaluated for DNA concentration, integrity, and purity. DNA concentration was measured by fluorimetry using a Qubit^TM^ 4 fluorometer (Thermo Fisher Scientific, Waltham, MA, USA), while integrity was assessed by electrophoresis in 1% agarose gels. Purity of DNA was determined using a BioDrop® spectrophotometer (BioDrop Ltd, Cambridge, UK), based on absorbance ratios at 260/280 nm and 260/230 nm, indicative of protein and organic compound contamination, respectively. We considered DNA samples acceptable if in electrophoresis they exhibited a distinct band larger than 20 kb with no visible degradation pattern (e.g., smearing), a concentration of at least 20 ng/uL, a 260/280 ratio between 1.8 and 2.0, and a 260/230 ratio between 1.6 and 2.2.

The twelve highest-quality DNA extractions of population samples, as determined by concentration, integrity, and purity criteria, were selected for subsequent library preparation. Double-digest restriction-site associated DNA (ddRAD) libraries were constructed following a standardized protocol, using the restriction enzymes *EcoRI* and *MseI* (Parchman et al., 2012). Barcode adapters were designed to allow for two sequencing errors without confusion of samples. Barcode adapters and sequencing primers were obtained from Integrated DNA Technologies (Coralville, Iowa, USA). We constructed libraries separately by species in four 96-well plates per species, and we randomized sample locations across the four plates using a random number generator in R. A total of 240 samples per species (12 individuals per population, 20 populations total) were multiplexed (pooled by species) and size selection was conducted using High-Prep PCR paramagnetic beads (MagBio Genomics, Gaithersburg, USA) to restrict the size range of each multiplexed library to between 300 to 500 base pairs. Single-end sequencing of the multiplexed libraries was conducted at the SGIker Genomic Services unit of the University of the Basque Country, Leioa, Spain, using an Illumina NovaSeq 6000 platform, to generate reads of 111 base pairs.

### Demultiplexing, SNP calling and Filtering

Demultiplexing of raw Illumina reads was carried out for each species separately using the process_radtags utility (Stacks v2.66; Catchen et al., 2013). In this step, we cleaned the data by discarding reads with any uncalled bases and reads with phred quality scores lower than 10. We rescued barcodes and RAD-Tag cut sites, and then trimmed the barcode adapter and deleted restriction site sequences. Cleaned reads were then processed through the Stacks pipeline for de novo loci assembly and SNP calling. First, ustacks was used to align reads from each individual and build putative loci. These loci were passed to cstacks to create a comprehensive catalog of consensus loci, selecting from each population the three individuals with the highest number of constructed loci. Then, we used sstacks to match each individual’s loci to the catalog. Tsv2bam and gstacks were used to align reads to the catalog and call SNPs across all individuals. Finally, the Stacks populations module generated population-level summary statistics and exported variant data to a Variant Calling Format (VCF) file, one for each species (Catchen et al., 2013) (see SI-2 for further information). We discarded one *H. atriplicifolium* population (PAV62) due to a misidentification of the species.

To ensure high-quality genotype data for downstream population genomic analyses, we filtered the VCF datasets using VCFtools v0.1.16 (Danecek et al. 2011). We retained only biallelic SNPs with a minor allele frequency higher or equal to 0.01 and filtered variants by read depth, keeping those with coverage between 6x and 120x to exclude low confidence calls and potential artefacts such as PCR duplicates or paralogs. After these filtering steps, we assessed data completeness and quantified the proportion of missing data per individual. We retained data from the ten individuals with the lowest levels of missing data in each population. We optimized the data for downstream analyses by testing different thresholds of allowed missing data and examining patterns of missingness across populations to identify any population-specific biases. Additionally, we retained one random SNP per RAD locus to eliminate closely linked loci.

### Population Structure and Genomic Clustering

We constructed a matrix of ones and zeros to represent missing data at SNP loci and produced a matrix of Jaccard distances among the samples. We used this matrix in a Non-metric Multidimensional Scaling ordination (cmdscale function; R-stats package, R Core Team, 2023) to examine the SNP data for substantial pattern in SNP missing values that might bias downstream analyses (Yi & Latch, 2022). Datasets were filtered to allow a maximum of 30% missingness per site, as this threshold balanced SNP retention with minimizing biased missingness across populations (Cerca et al., 2021). We proceeded directly with analysis of genotypic data due to the absence of strong patterns of missingness. Subsequently, population genomic statistics, including Nei genetic distances and pairwise population F_st_ values, were calculated using the R package StAMPP v1.6.3 (Pembleton et al., 2013). Nei’s distances were visualized as heatmaps to interpret genomic similarities and differentiation between populations. Pairwise F_st_ values were converted into a distance matrix, geographic distances were calculated with the Haversine formula, and their correlation was tested with Mantel tests in the vegan R package (Oksanen et al., 2025). Then, after verifying the absence of potentially influential patterns in missing SNP data, we used Principal Component Analysis (PCA) of genotypic values of individuals to explore the structure and genotype clustering of populations.

We conducted admixture analyses with ADMIXTURE v1.3.0 (Alexander et al., 2009) to further investigate population structure and patterns of population ancestry. We standardized the analyses by including 10 individuals per population, in order to minimize bias that can arise from unequal sampling intensity among populations, thus reducing the possibility of spurious admixture signals driven by among-population differences in sample size (Toyama et al., 2020). We examined the optimal number of potential ancestral populations by comparing the cross-validation (CV) errors for values of K ranging from 1 to 10. The K value with the lowest CV error rate was selected as the best-fit. Minimizing of CV error when selecting K ensured the robustness of ancestry inference to overfitting and best reflected genomic structure (Alexander et al., 2009). For the chosen K, we calculated the average proportion of each inferred ancestral group in each population. These proportions were then visualized geographically using pie charts overlaid on population sampling locations to facilitate interpretation of spatial genomic structure.

### Phylogenomic Inference of Historical Population Dynamics

We used TreeMix v1.13 (Pickrell & Pritchard, 2012) to model population splits using genome-wide allele frequency data, and to infer historical relationships and gene flow among populations. Original filtered VCF files were converted to TreeMix input format using the populations utility from Stacks v2.66. Since TreeMix is highly sensitive to missing data, we initially tested datasets with no allowed missingness; however, this retained fewer than 600 SNPs per species. To balance data completeness with marker density, we retained SNPs with up to 10% missingness in *P. purpurea* and *H. atriplicifolium* datasets, and up to 20% missingness for *Lavandula stoechas*, for which a stricter threshold resulted in an insufficient number of SNPs. Missing genotypes were imputed using allele frequency estimates within populations using customized scripts in the BASH utility awk, enabling TreeMix to calculate covariance matrices from consistent data.

We followed a standardized protocol for TreeMix analyses (Dahms, 2021). First, we ran multiple TreeMix replicates across a range of migration events (from 1 to 10) to build a consensus tree and identify consistent topologies. We defined outgroups for rooting the trees by selecting the most differentiated populations in the PCA and ADMIXTURE analyses (Zhou et al., 2024) for *P. purpurea* and *L. stoechas* (PAV76 and PAV8, respectively). However, since there was not a clearly differentiated population of *H. atriplicifolium* among the 10 serpentine/non-serpentine population pairs, we used a genetically and geographically distant population from another project (PAV109, not one of the 20 populations, but in the original round of SNP calling and in the VCF). Second, we applied the OptM R package (Fitak, 2021) to determine the optimal number of migration edges based on changes in likelihood values. Third, we performed 30 additional TreeMix runs with the selected optimal number of migration events. Finally, we visualized the final rooted tree and annotated it with inferred migration weights, providing both topological structure and quantitative estimates of historical gene flow between populations. A schematic overview of the study design and workflow to facilitate interpretation of the results appears in Fig. 2.

**Figure 2.**
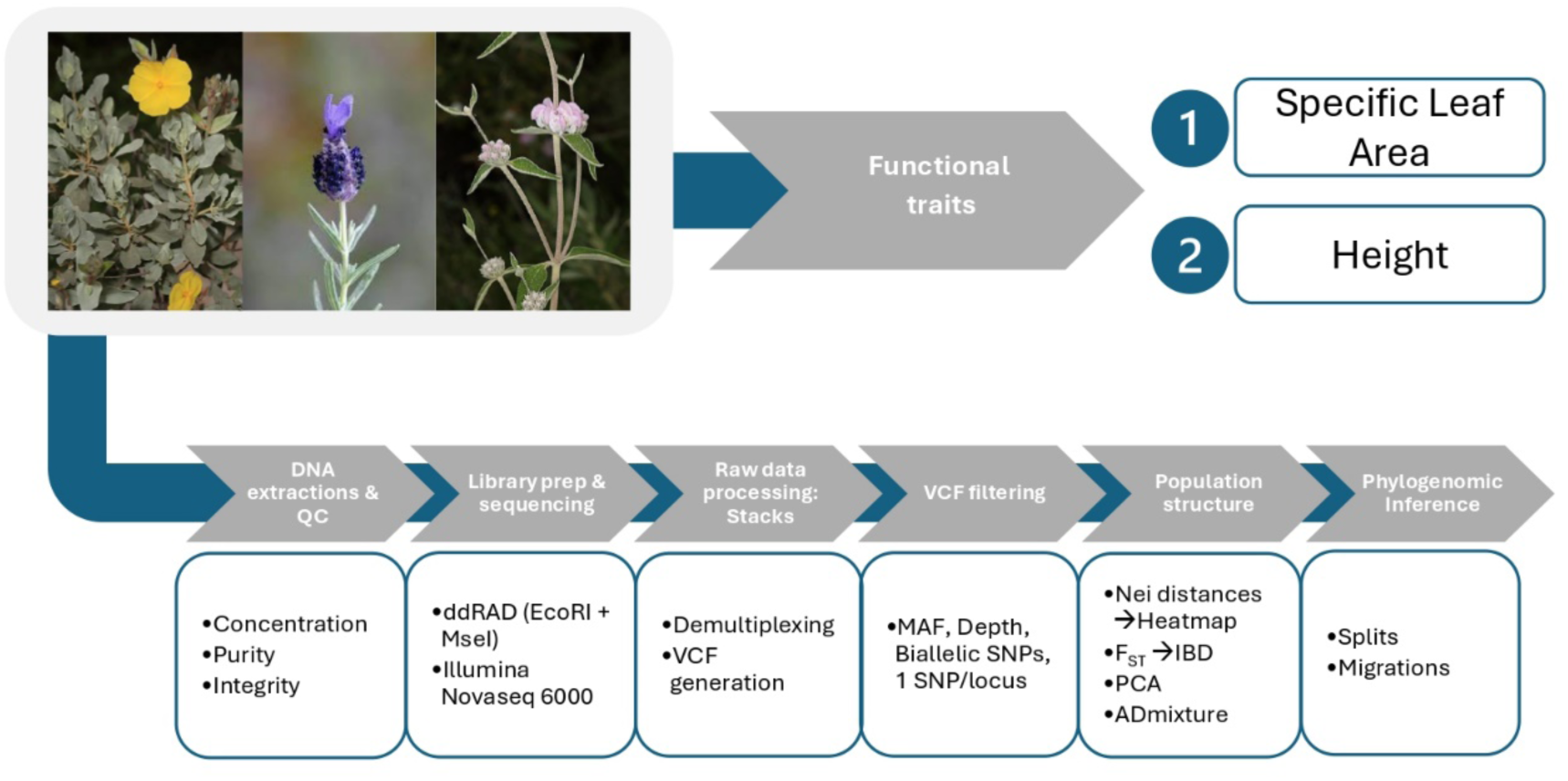
Schematic overview of the workflow. The diagram summarizes the main experimental and analytical stages, which are divided into functional trait and genomic analyses.

## Results

### Functional traits

Linear mixed-effect models, with populations as random factors, detect significant differences in plant height for *L. stoechas* and *P. purpurea*, with serpentine populations exhibiting values on average 13.56 cm ± 5.65 s.e. and 16.35 cm ± 4.84 s.e. shorter, respectively. *H. atriplicifolium* also exhibits reduced height, as serpentine populations are on average 6.86 cm ± 8.99 s.e. smaller than non-serpentine populations, although this difference was not statistically significant (SI3-Figure S1 and Table S2). In contrast, SLA values do not differ significantly between serpentine and non-serpentine populations in any of the three species (SI3-Figure S2 and Table S3).

### Genomic Structure and Clustering

We characterize genomic structure across populations of the three focal species using PCA, pairwise genetic distances, and ancestry-based clustering. The filtered VCF datasets with 30% allowed missingness per site (Cerca et al., 2021) retain differing numbers of high-quality SNPs: 135,693 for *H. atriplicifolium,* 19,766 for *L. stoechas*, and 433,823 for *P. purpurea*. Patterns of missing data across individuals and populations show no evident structure (SI-4 Figure S1). Heatmaps of Nei’s genetic distances among populations provide visualization of populations similarities and dissimilarities for each species (SI-5 Figure S1). In these heatmaps, we observe distinct patterns of population similarity across the three species. For *P. purpurea*, most populations show high similarity, with two clear outliers: PAV76 and PAV83. A similar trend appears in *L. stoechas* plot, where PAV8 and PAV87 stand out as notably different from the other populations. In contrast, *H. atriplicifolium* displays a more structured pattern, with clear groupings of similar populations and greater differentiation between groups. Mantel tests to detect IBD indicate that *H. atriplicifolium* exhibits the strongest isolation-by-distance signal, with pairwise F_st_ values highly correlated (r = 0.763, p = 0.001) with geographic distances. *L. stoechas* shows a moderate but significant correlation (r = 0.404, p = 0.001), while *P. purpurea* displays the weakest, yet still significant, association (r = 0.347, p = 0.005) between geography and genomic differentiation (SI-6 Figure S1).

Principal Component Analyses reveal distinct patterns of genomic clustering among the species (Fig. 3). Populations of *H. atriplicifolium* form well-separated clusters in PCA space, with individuals tightly grouped by population and clear separation between different populations (Fig. 3a1). This structure is mirrored in the ADMIXTURE results (Fig. 4a), which show minimal admixture between populations and distinct ancestry profiles consistent with the geographical proximity of the populations. In contrast, *P. purpurea* and *L. stoechas* display weaker population structure. PCA biplots for both species reveal a central cluster containing the majority of populations, with a few clearly differentiated populations positioned outside this main cloud of points (Fig. 3b1 and 3c1). These patterns indicate partial population subdivision, with more diffuse genomic structure than in *H. atriplicifolium*. ADMIXTURE analyses are consistent with these results (Fig. 4b and 4c), with individuals in the central cluster generally exhibiting shared ancestry components, while peripheral populations in PCA space show distinct ancestry profiles and individuals with little admixture. For *H. atriplicifolium,* optimal number of admixture groups is found with K equal to three, while for both *P. purpurea* and *L. stoechas,* four admixture groups minimized the CV errors (SI-7 Figure S1). Examination of the clustering of sample values in the PCA space reveal that *L. stoechas* and *P. purpurea* exhibit clearer separation between individuals of serpentine and non-serpentine origin than does *H. atriplicifolium* (Fig. 3b2, c2). In contrast, *H. atriplicifolium* in PCA analysis shows greater overlap between samples from different soil types than do values of the other two species (Fig. 3a2).

**Fig 3.**
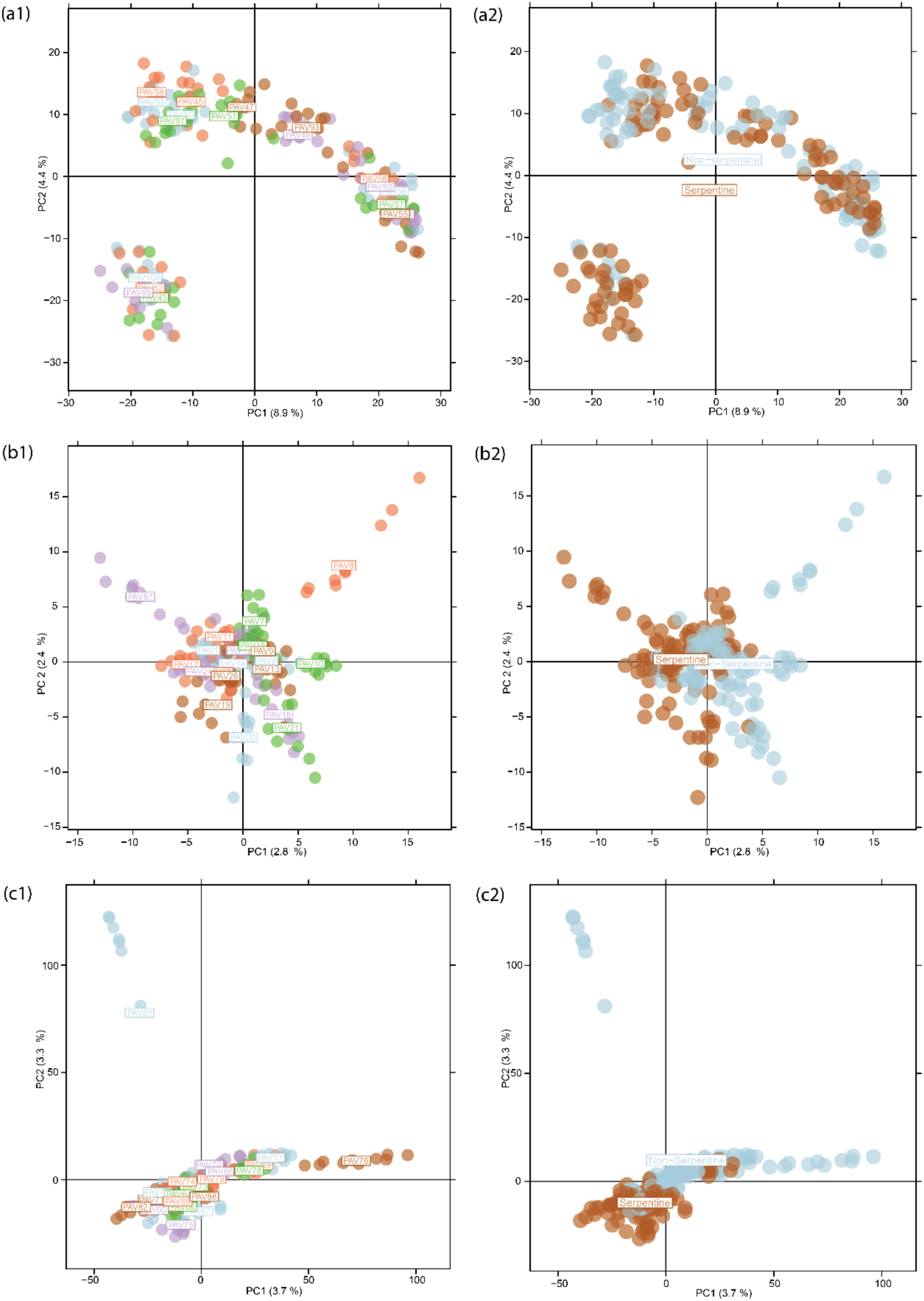
Principal Component Analysis (PCA) of SNP data for the three focal species. Each panel displays individual genotypes projected onto the first two principal components: (a) *Halimium atriplicifolium*, (b) *Lavandula stoechas*, and (c) *Phlomis purpurea*. Each dot represents an individual, coloured by population in left panels, and by soil type in right panels. Population labels correspond to the centroid of the individuals from each population in PCA space.

**Fig 4.**
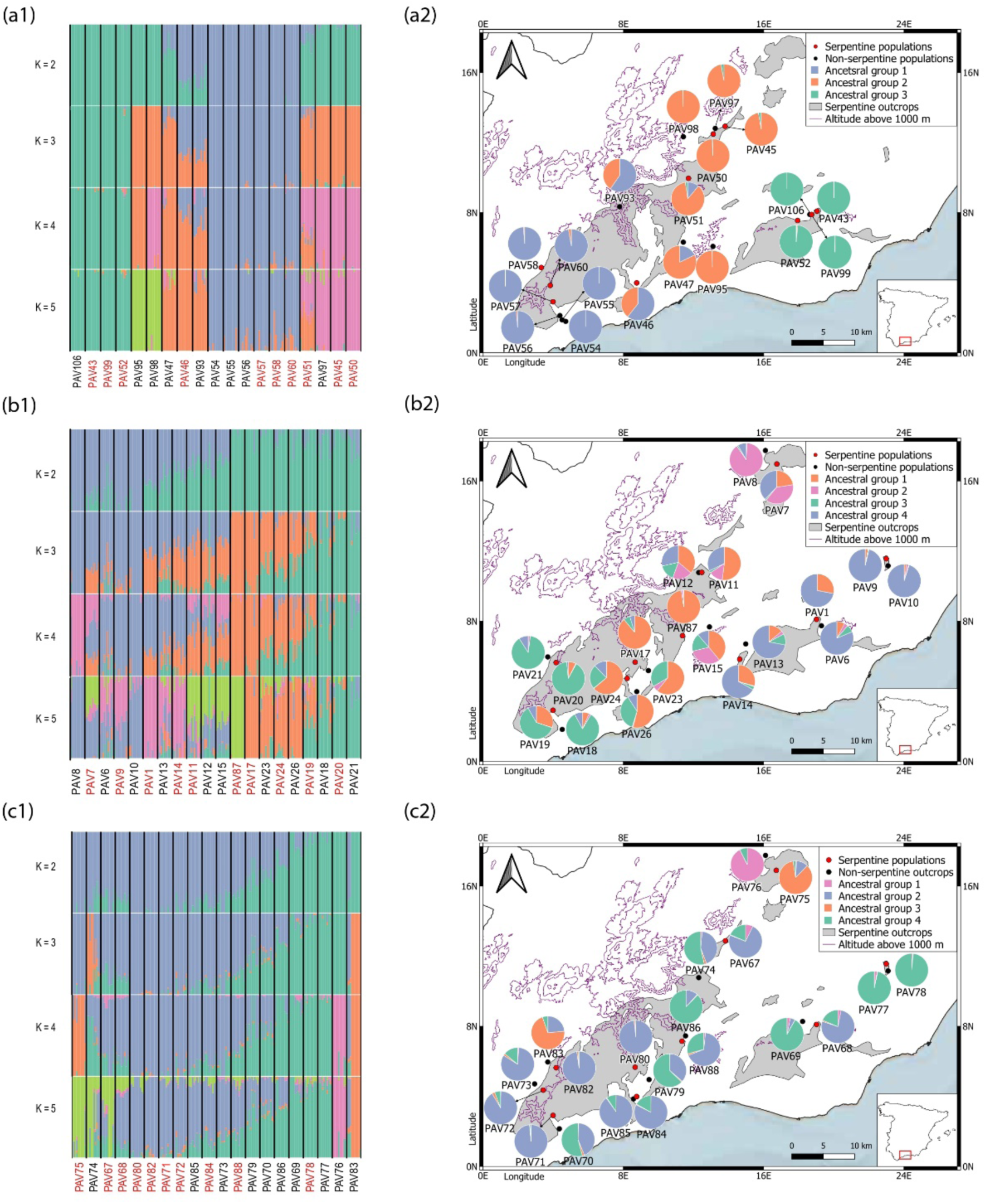
Genomic structure and spatial distribution of inferred ancestry by ADMIXTURE for the three focal species. Panels (a1-c1) show individual ancestry proportions for values of K=2 to K=5, for (a) *Halimium atriplicifolium*, (b) *Lavandula stoechas,* and (c) *Phlomis purpurea.* Each vertical bar represents an individual, grouped by population and coloured by inferred ancestral components. Red text indicates populations on serpentine while black text indicates non-serpentine soil Panels (a2-c2) display maps of the study region with population locations overlaid with pie charts representing the average ancestry proportions for optimal K: K=3 for *H. atriplicifolium* and K=4 for *L. stoechas* and *P. purpurea*.

We calculated the average ancestry proportions for each inferred cluster across populations. Distinct ancestry profiles across species emerge when pie charts of these proportions are ordered in geographic space (Fig. 4). *H. atriplicifolium* shows strong geographical structuring, with populations forming groups based on geographic location, while soil type appears to have little influence, and admixture between populations is minimal (Fig. 4a1, a2). In *L. stoechas*, although geography plays a role, an association with edaphic classification is more apparent than in data for *H. atriplicifolium*. Serpentine populations generally show more of Ancestral group 1 (orange) than do their paired non-serpentine populations, indicating a genomic signal of shared ancestry in the serpentine populations (Fig. 4b1, b2). Geographical influence is even less evident in *P. pupurea*, and ancestral patterns align more closely with edaphic classification (Fig. 4c1). This is most visible in Fig. 4c2, in which serpentine populations tend to have a larger component from Ancestral group 2 (blue) than do their non-serpentine populations in every case except one (PAV77, PAV78).

### Phylogenomic Relationships and Historical Gene Flow

We infer population relationships and potential historical gene flow events for each species using TreeMix analyses based on allele frequency data. After filtering for allowable levels of missing data (10% for *H. atriplicifolium* and *P. purpurea*; 20% for *L. stoechas*), and imputing missing genotypes using population-specific allele frequencies, each dataset contained a sufficient number of SNPs for model estimation. The final SNP counts used in TreeMix were 54,498 for *H. atriplicifolium*, 8,854 for *L. stoechas*, and 150,748 for *P. purpurea*.

Across all three species, TreeMix analyses produce well-resolved phylogenetic trees that broadly mirrored the population clustering patterns identified through PCA and ADMIXTURE analyses. In *H. atriplicifolium* (Fig. 5a), population relationships are clearly delineated, with strong tree structure and minimal residual covariance, especially among geographically distant populations, indicating limited historical gene flow. Using optM R package, we inferred two historical gene flow events for this species (SI-8 Figure S1), both involving long-distance connections between non-serpentine populations. These results suggest that *H. atriplicifolium* populations are relatively isolated, with rare but notable instances of gene flow across broader geographic distances.

**Fig. 5.**
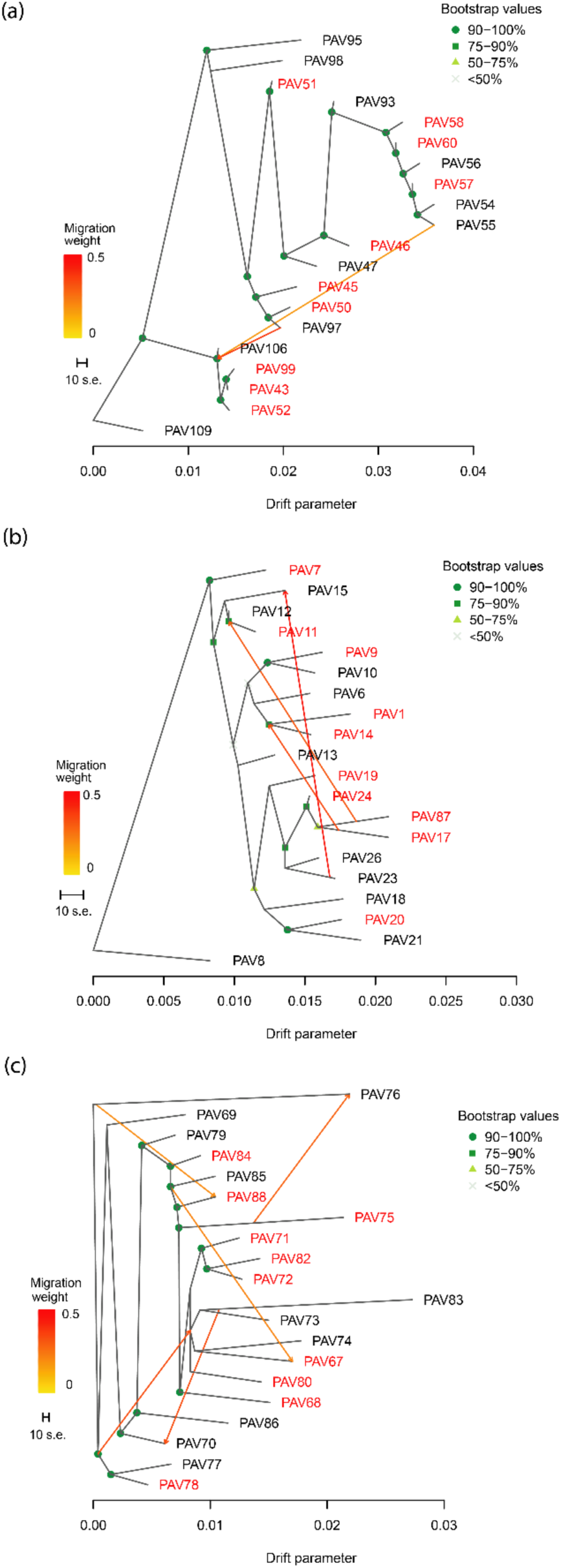
Historical population relationships and inferred gene flow from TreeMix analyses. (a) *Halimium atriplicifolium*, (b) *Lavandula stoechas,* and (c) *Phlomis purpurea*. Each panel shows the consensus tree with the optimal number of gene flow events indicated by arrows, with direction and intensity corresponding to migration weight. Red text represents serpentine and black text non-serpentine populations.

In contrast, *L. stoechas* (Fig. 5b) and *P. purpurea* (Fig. 5c) exhibit more complex, reticulate patterns accompanied by higher population diversity, with higher inferred levels of gene flow. Both of these species are abundant in individuals and populations (P.A. pers. obs.). Three gene flow events are identified in *L. stoechas*, and its tree topology reflects more recent divergence and connectivity between populations. While serpentine populations are scattered throughout the tree, some level of grouping is still observable, suggesting that certain serpentine populations may share common ancestry or parallel evolutionary histories. Nonetheless, the overall structure indicates admixture and little separation along edaphic lines.

*P. purpurea* displayed the most intricate pattern, with five inferred gene flow events. Several of these connect populations across both ecological and geographic boundaries. One notable case of gene flow is observed between geographically proximate serpentine and non-serpentine populations, PAV75 and PAV76, presenting the only case observed among 60 possible pair combinations across the three species. Still, we observe that several serpentine populations cluster together in the tree (PAV71, PAV72, PAV82), indicating closely related populations on the Sierra Bermeja massif. The multiple gene flow events and relatively low drift levels in *P. purpurea* compared to the other two species reflect a dynamic demographic history involving both historical divergence and intermittent connectivity.

## Discussion

We show that all three focal species underwent multiple, independent colonisations of serpentine sites. These parallel colonisations resemble well-documented serpentine habitation in other taxa, such as for the tetraploid *Arabidopsis arenosa* (Arnold et al., 2016) and the diploid *Arabidopsis lyrata* (Turner et al., 2010). Across replicated serpentine regions in southern Spain, we detect consistent reductions in plant height among serpentine populations compared to ones off serpentine, in agreement with general trends towards smaller stature under the nutrient-poor, metal-rich conditions of serpentine soils, and generally regarded part of the broader serpentine syndrome (Brady et al., 2005; Hidalgo-Triana et al, 2023). Such phenotypic differences may also reflect environmentally induced plasticity rather than fixed genomic divergence, and distinguishing between plastic and heritable components of these traits will require controlled transplant experiments (O’Dell & Rajakaruna, 2011). In contrast, we find no significant differences in specific leaf area (SLA) between serpentine and non-serpentine populations. This finding contrasts with studies that report lower SLA on serpentine plants (Adamidis et al., 2014; Hidalgo-Triana et al., 2023), but parallels report of weak or absent SLA differentiation in some serpentine systems (Delhaye et al., 2024). Finally, the patterns of genomic differentiation we observe here are species-specific, underscoring the importance of examining multiple species when investigating how landscape features and ecological contexts shape population structure and evolutionary history (Todesco et al., 2020).

### Species-specific genomic responses

Species-specific genomic responses to environmental heterogeneity are increasingly recognized as a common outcome when different lineages interact with selective pressures, dispersal limitations, and historical demographic processes (Dauphin et al., 2023; Bonte et al., 2024; Hou et al., 2022). In the context of serpentine soils, numerous studies have shown that plant populations often exhibit strong local adaptation and limited gene flow across edaphic boundaries, even when serpentine and non-serpentine populations occur in close proximity (Selby & Willis, 2018; Anacker, 2014). However, the strength and nature of genomic differentiation in response to serpentine habitats vary widely among species, often depending on life history traits, dispersal mechanisms, and population structure (Turner et al., 2010; Preite et al., 2019). Some studies illustrate differences in evolutionary potential across species, suggesting that species-specific habitat requirements and responses to environmental stressors may be better predictors of evolutionary patterns than simply the presence of strong, physiologically relevant environmental gradients (Nielsen et al., 2020).

*H. atriplicifolium* displays strong genomic structure shaped primarily by geographic separation rather than substrate type. Serpentine and non-serpentine populations in this species are generally scarce and distantly separated in our study area (Blanca et al., 2011), reducing opportunities for gene flow across soil boundaries. Additionally, this species likely experiences pollen limitation, meaning that its reproductive success may be constrained by low pollen delivery (Ashman et al., 2004; Teixido 2014). This is consistent with the strong geographic pattern in the genomic structure of *H. atriplicifolium*, indicating that physical distance and limited dispersal, rather than ecological or edaphic differentiation, are the primary drivers of genomic divergence. In this context, limited pollinator-mediated gene flow may act as a key mechanism maintaining spatial genomic structure (Feigs et al., 2022).

In contrast, *L. stoechas* and *P. purpurea* display weaker genomic structure in the PCA and ADMIXTURE plots, and higher inferred levels of gene flow in TreeMix analyses. Both species are pollinated by large, mobile insects, such as bumblebees, honeybees and long-tongued bees, which are known to facilitate pollen movement over hundreds of meters or more (Petanidou & Vokou, 1993; de Vega et al, 2022). In addition, populations of these two species are abundant and widely distributed (Blanca et al., 2011). Indeed, serpentine populations in the Sierra Bermeja region consistently show elevated proportions of a distinct ancestral component, suggesting a localized substrate-associated ancestry, with some serpentine populations forming monophyletic clades within Sierra Bermeja, the largest continuous serpentine outcrop in the Iberian Peninsula. This highlights the species-specific genomic differentiation patterns in our dataset, as this differs with the general pattern in *H. atriplicifolium*.

This species-specificity has also been observed in plants across other environmental gradients, such as elevation (Chapman et al., 2013) and aridity (Hu et al., 2021), where co-occurring species often display contrasting patterns of neutral genomic differentiation. The variability observed among species in our study underscores the importance of multispecies approaches, as the results point to complex ecological, evolutionary, and biogeographical dynamics that shape population differentiation across heterogeneous landscapes.

### Multiple, independent colonisations of serpentine sites

Our results support the hypothesis of multiple, independent colonization events into serpentine habitats across the three species, rather than the expansion of a single serpentine-associated lineage across the landscape, which is consistent with our original prediction. This aligns with findings in other serpentine systems, where recurrent, independent colonisations are increasingly recognized as a common evolutionary outcome, for example, in *Silene paradoxa* (Lazzaro et al., 2021), and *Solidago virgaurea* (Sakaguchi et al., 2019), reviewed in Konečná et al., 2020. Single origins of serpentine populations are rarely reported among facultative serpentinophytes species (Sakaguchi et al., 2018), but are observed in serpentine endemics (Cecchi & Selvi, 2009). The presence of divergent genomic backgrounds among serpentine populations supports the idea that serpentine habitation in these species has arisen repeatedly and independently, shaped by local demographic and ecological contexts. These repeated colonisations of serpentine habitats suggest the potential for multiple evolutionary pathways to overcoming the pressures imposed by serpentine soils. This would further suggest that serpentine habitation is not constrained to a single genomic solution but could instead arise though diverse genetic and ecological routes (Arnold et al., 2016, Konečná et al., 2020). However, in some cases, such as *Arabidopsis arenosa*, similar adaptive pathways have repeatedly evolved from shared standing genetic variation (Konečná et al., 2021), underscoring the flexibility of evolutionary responses to extreme edaphic stress.

In both *P. purpurea* and *L. stoechas*, some serpentine populations from different outcrops exhibit distinct ancestry profiles, suggesting multiple colonisations, but with the notable exception of Sierra Bermeja where multiple populations with a shared ancestry are found. This example indicates that, once radiated to serpentine, populations have the potential to disperse widely and colonize additional adjacent habitat (Anacker et al., 2011). In *H. atriplicifolium*, there is no evidence that populations across distant geographical regions share ancestry of serpentine habitat. This observation further supports the model of multiple, geographically isolated serpentine colonisations. Together, our PCA, ADMIXTURE, and TreeMix analyses converge to support a scenario where serpentine habitation has evolved multiple times independently within each species, reflecting the complex interplay of geography, limited gene flow, and local edaphic constraints. These recurrent colonisations could have been facilitated by a broad pool of potentially adaptive variation already present in non-serpentine populations, a hypothesis that warrants further testing in future studies. By comparing these patterns across the three species, our results provide strong evidence that serpentine colonization has not only been shaped by lineage-specific attributes but also by the ecological and geographic context in which serpentine habitation occur (Harrison & Rajakaruna, 2011), highlighting the importance of a multispecies, landscape-scale approach to understand the evolutionary dynamics of edaphic specialization.

### Historical gene flow and phylogenomic inference

Overall, our results indicate that *H. atriplicifolium* retains strong geographic structure with minimal admixture, while *L. stoechas* and *P. purpurea* reveal greater gene flow and more complex evolutionary relationships. Importantly, across all three species, we detect signals of grouping among some serpentine populations, suggesting that despite gene flow and geographic mixing, edaphic specialization still plays a role in shaping phylogenetic structure. These results highlight the role of serpentine environments as natural systems for studying ecological speciation, where strong selective pressures interact with partial reproductive isolation to maintain divergence despite potential gene flow. Such dynamics underscore how localized ecological barriers can contribute to long-term biodiversity patterns and endemism in serpentine floras (Rajakaruna, 2018; Sianta & Kay 2021).

Despite geographic proximity, signals of gene flow between proximate serpentine and non-serpentine populations appear limited in our analyses, with only a single inferred gene flow event bridging these habitats, observed in *P. purpurea*. However, the methods used here, such as TreeMix, have limited power to detect gene flow between recently diverged or closely related serpentine and non-serpentine population pairs, and thus the apparent absence of admixture should not be interpreted as definitive evidence for reproductive isolation. Limited gene flow could be influenced by intrinsic reproductive barriers, such as differences in flowering times, mating system dynamics or genomic incompatibilities, which can evolve alongside habitation of contrasting soil types (Nosil et al., 2009; Dittmar & Schemske, 2018; Toll & Willis, 2024). The exception found in *P. purpurea* may reflect a localized breakdown of these barriers, potentially due to recent secondary contact or microhabitat transition zones that temporarily reduce reproductive isolation, or because intrinsic incompatibilities between serpentine and non-serpentine populations have not yet fully developed (Sianta & Kay, 2021). Nevertheless, its singularity underscores the general trend: serpentine and non-serpentine populations, exhibit limited gene flow even when closely situated, suggesting that edaphic specialization potentially limits gene flow and contributes to the maintenance of ecologically divergent lineages (Anacker 2014).

### Evolutionary and conservation implications

The contrasting patterns of genomic connectivity between serpentine and non-serpentine populations raise important questions about their evolutionary trajectories under future environmental change. While current phenological or reproductive barriers may limit gene flow, occasional or facilitated gene flow, especially from serpentine to non-serpentine populations, could prove sources of adaptive variation under increasing climatic stress (Hoffmann & Sgrò, 2011). Serpentine populations, with their evolved tolerance to drought, metal toxicity, and nutrient-poor soils, may harbour alleles conferring resilience to extreme climatic conditions (Brady et al., 2005; Harrison et al., 2015). This directionality from serpentine populations suggests that they could act as a reservoir of adaptive variation for non-serpentine populations. Conversely, persistent genomic isolation may increase vulnerability to environmental change by reducing the pool of shared genetic diversity available for adaptation (Jump et al., 2009). This underlines the conservation importance of maintaining both serpentine and non-serpentine populations, not only to preserve independent evolutionary trajectories, but also to safeguard the full complement of genomic diversity across environmental gradients. Collectively, these results support the view that serpentine habitats function not just as ecological islands but as evolutionary crucibles, generating and maintaining unique adaptive variation with potential importance far beyond their limited spatial extent.

Finally, interspecific differences in SNP density and genome complexity may influence the resolution of inferred population structure and gene flow, potentially affecting cross-species comparisons (Andrews et al, 2016). Additionally, unmeasured environmental factors, such as climatic variation, biotic interactions, or edaphic heterogeneity beyond serpentine and non-serpentine dichotomies, could also influence genomic patterns. To extend understanding of serpentine habitation and ecological divergence, future studies should integrate genome-wide scans to identify candidate loci under differential edaphic selection (Lotterhos & Whitlock, 2015), as well as reciprocal transplant or common garden experiments, in order to assess fitness trade-offs across soil types (Brady et al, 2005). Ecological niche modelling, particularly when combined with genomic data and climate projections, could also help predict the spatial dynamics of serpentine habitation under changing environmental conditions (Warren et al., 2010). Such integrative approaches will be key to disentangling the roles of environment, demography, and selection in shaping evolutionary responses across heterogeneous landscapes.

## Acknowledgements

We thank Eurofins AgroSciences for soil analysis. Thanks to SGiker genomic services, University of the Basque Country, for technical assistance and sequencing of ddRAD libraries. Thanks to A. Torre for help with L. stoechas DNA extractions. The research was supported by grant PID2020-118028GB-I00 to P.B.P from the Spanish Ministerio de Ciencia, Innovación y Universidades. P.A. was supported by fellowship PRE_2024_2_0288 from the Basque Government and by travel award FM/d/2023-2-003 from Charles University, Prague, Czech Republic.

## Data Accessibility

The raw ddRAD sequencing data and required metadata have been deposited in the NCBI Sequence Read Archive (SLA) Bioproject PRJNA1338063. Additional metadata describing samples, sequencing libraries, and experimental design, as well as other relevant files required to reproduce the analyses, are available at https://doi.org/10.5281/zenodo.17144186.

## Author Contributions

P.A. designed field sampling of botanical material with assistance from P.B.P., and was assisted in collecting by N.H.-T., N.K., and J.F.P.-O., who also conducted analysis of SLA. A.V.P.-L. provided additional guidance on sampling design, and filtered potential locations of plant populations. P.A. collected soil samples for metal analysis and analysed them with assistance from A.G.-F. N.K. contributed to map figures. A.L.-E. supported sample preservation and DNA extractions. P.A. wrote the first draft of the manuscript, and all co-authors contributed to its further conceptual development and writing, and approved the final version. P.B.P. conceived the research, secured funding, and supervised the project.

