## Supplementary Information SI-1 to SI-8 for "Independent colonisations of serpentine habitats highlight species-specific evolutionary histories of lineage diversification"

**SI-1:** Chemical analysis of the soil samples

**SI-2:** Methods on demultiplexing and SNP calling

**SI-3:** Phenotypic variation across soil types

**SI-4:** Missingness patterns of individuals at SNP loci.

**SI-5:** Nei's distances among populations.

**SI-6:** Isolation by distance (IBD) tests.

**SI-7:** Cross-validation errors from ADMIXTURE runs.

**SI-8**: Selection of migration edges (m) in TreeMix.

#### SI-1: Chemical analysis of the soil samples

Soil samples were analysed by Eurofins Analytico (Parets del Valles, Spain, https://www.eurofins-environment.es) for a range of soil parameters (SI-1 Table 1). We analysed several physicochemical properties of each soil sample, including pH, the content of zinc (Zn), lead (Pb), arsenic (As), mercury (Hg), copper (Cu), iron (Fe), chromium (Cr), and nickel (Ni), and the calcium-to-magnesium ratio (Ca:Mg). To explore potential patterns in soil composition, particularly in relation to serpentine and non-serpentine origins, we conducted principal component analysis (PCA) using the prcomp() function from the stats package in R v4.3.1. This analysis assessed whether the two nominal soil types formed distinct clusters based on the measured variables. Subsequently, we identified the variables that most strongly contributed to the separation observed in the PCA and performed one-way ANOVA tests on these variables to determine whether their concentrations differed significantly between serpentine and non-serpentine soils. These analyses, combined with visual assessment of soil appearance and the dominant vegetation observed at each sampling site, were used to confirm the nominal soil classification for each location.

Principal component analysis (PCA) of measured soil parameters reveals a consistent and clear separation between putative serpentine and non-serpentine sites across all three species. In each case, PCA ordinations show two distinct groups with no overlap between nominal soil types (SI-1 Figure 1). This separation is consistent across species, reflecting shared underlying environmental gradients across the sampling region. The primary variables contributing to this division include notably elevated concentrations of nickel, iron, and chromium in serpentine soils, alongside substantially lower calcium-to-magnesium ratios. One-way ANOVA tests confirm that differences in Ni, Fe, and Cr concentrations, as well as Ca:Mg ratios, are statistically significant between soil types in all three species (SI-1 Figure 1), with the exception of Ca:Mg ratios in *H. atriplicifolium*, where the trend is in the expected direction but not statistically significant. Evidence from multivariate analysis, individual soil variables, and field validation provides robust support for the initial classification of soil type, validating the use of these designations.

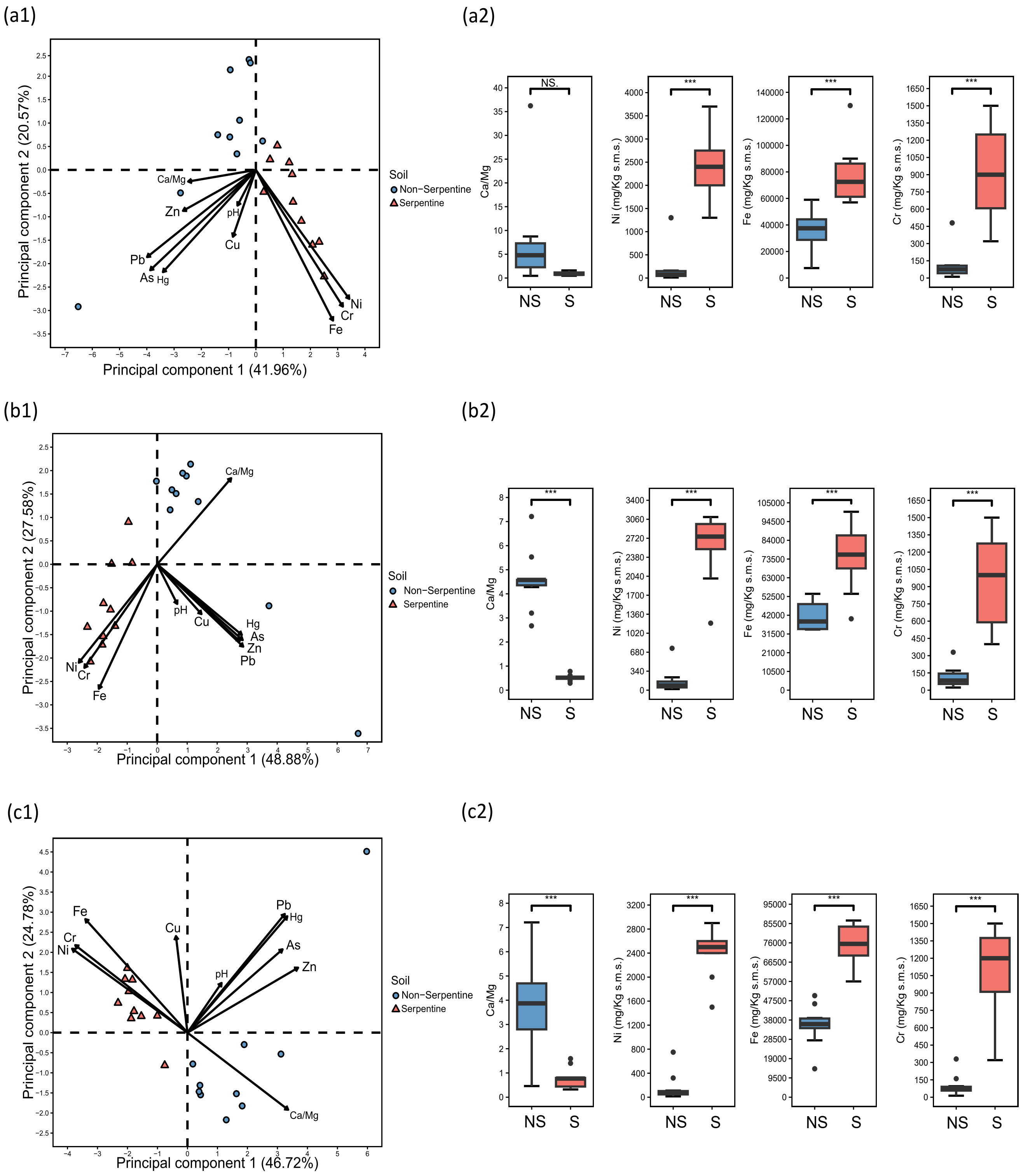


**SI-1 Figure S1**. Chemical properties of collection sites across the three focal species. (a) *Halimium atriplicifolium*. (b) *Lavandula stoechas*. (c) *Phlomis purpurea.* (a1-c1) Principal Component Analysis (PCA) of soil samples based on measured chemical parameters. Arrows represent the loadings of the variables, showing their contribution to the PCA axes. (a2-c2) Boxplots showing the distribution of four key variables: calcium-to-magnesium ratio, and concentrations of nickel, iron, and chromium. Asterisks indicate significance levels from one-way ANOVA comparisons (***$p < 0.001$; NS: not significant).

#### SI-2: Methods on demultiplexing and SNP calling

Demultiplexing of raw Illumina reads was carried out for each species using the process_radtags utility included in Stacks v2.66. In this step, we cleaned the data, discarding reads with any uncalled base and reads with phred quality scores lower than 10. We rescued barcodes and RAD-Tag cut sites, and then trimmed the barcode and cut site sequences. Cleaned reads were then processed through the Stacks pipeline for de novo loci assembly and SNP calling. First, ustacks was used to align reads from each individual and build putative loci, with the minimum depth of coverage required to create a stack (-m) set to 3 (default), and the maximum distance allowed between stacks within individuals (-M) also set to 3, as it helps group similar reads into the same locus, allowing for allelic variation and sequencing errors, but avoiding collapsing (pooling) of distinct loci. The resulting loci were then passed to cstacks to create a catalog of consensus loci across all individuals. In this case, three individuals per population with the highest number of constructed loci were selected for catalog construction. Then, we used sstacks to match each individual's loci to the catalog. Tsv2bam and gstacks were used to align reads to the catalog and call SNPs across all individuals. Finally, the Stacks populations module generated population-level summary statistics and export variant data to three VCF files, one for each species.

<u>**SI-3: Phenotypic variation across soil types**</u>

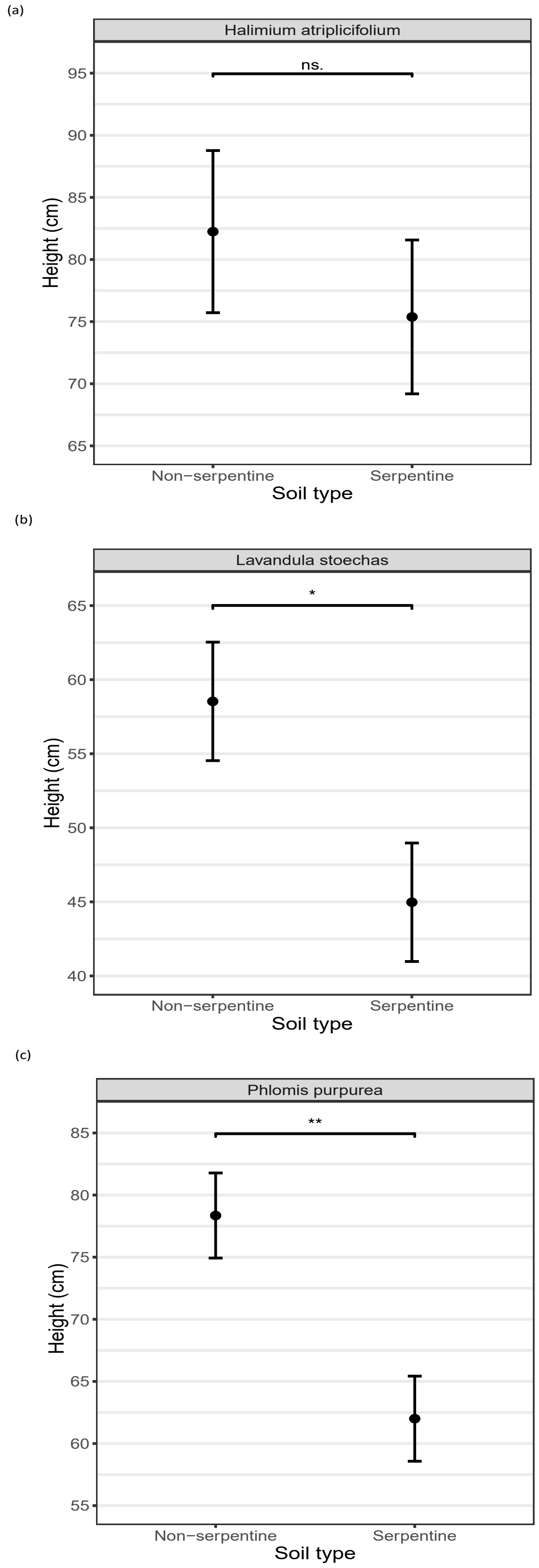


**SI-3 Figure S1**: Effects of soil type on plant height for (a) *H. atriplicifolium*, (b) *L. stoechas,* and (c) *P. purpurea*. Estimated marginal means and standard errors from linear mixed-effects models with populations as random effect. Significant differences in soil type are indicated by asterisks.

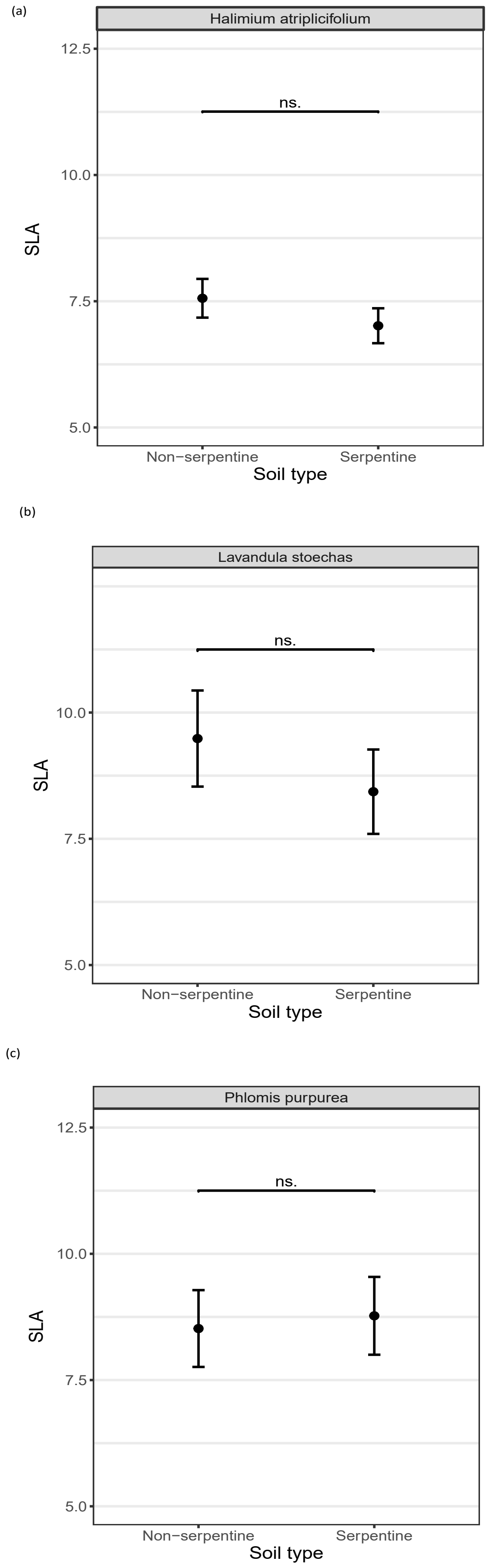


**SI-3 Figure S2**: Effects of soil type on plant Specific Leaf Area (SLA) for (a) *H. atriplicifolium*, (b) *L. stoechas,* and (c) *P. purpurea*. Estimated marginal means and standard errors from linear mixed-effects models with populations as random effect. Significant differences in soil type are indicated by asterisks.

**SI-3 Table S2**: Fixed effect estimates of soil type (serpentine vs. non-serpentine) on plant height for three species from linear mixed-effects models with population as random effect. Bold denotes statistical significance.

| Fixed effect | Estimate | SE | F | P |
|---|---|---|---|---|
| | | *Halimium atriplicifolium* | | |
| **$Soil_{serpentine}$** | -6.864 | 8.995 | $F_{1,17}$= 0.582 | 0.456 |
| | | *Lavandula stoechas* | | |
| **$Soil_{serpentine}$** | **-13.560** | **5.657** | **$F_{1,18}$= 5.746** | **0.027** |
| | | *Phlomis purpurea* | | |
| **$Soil_{serpentine}$** | **-16.353** | **4.843** | **F1,18= 11.401** | **0.003** |

**SI-3 Table S3**: Fixed effect estimates of soil type (serpentine vs. non-serpentine) on plant SLA for three species from linear mixed-effects models with population as random effect.

| Fixed effect | Estimate | SE | F | P |
|---|---|---|---|---|
| | | *Halimium atriplicifolium* | | |
| **$Soil_{serpentine}$** | -0.544 | 0.514 | $F_{1,16.93}$ = 1.115 | 0.306 |
| | | *Lavandula stoechas* | | |
| **$Soil_{serpentine}$** | -1.053 | 1.265 | $F_{1,15.24}$ = 0.692 | 0.418 |
| | | *Phlomis purpurea* | | |
| **$Soil_{serpentine}$** | 0.252 | 1.082 | $F_{1,18.22}$ = 0.054 | 0.818 |

### SI-4: Missingness patterns of individuals at SNP loci

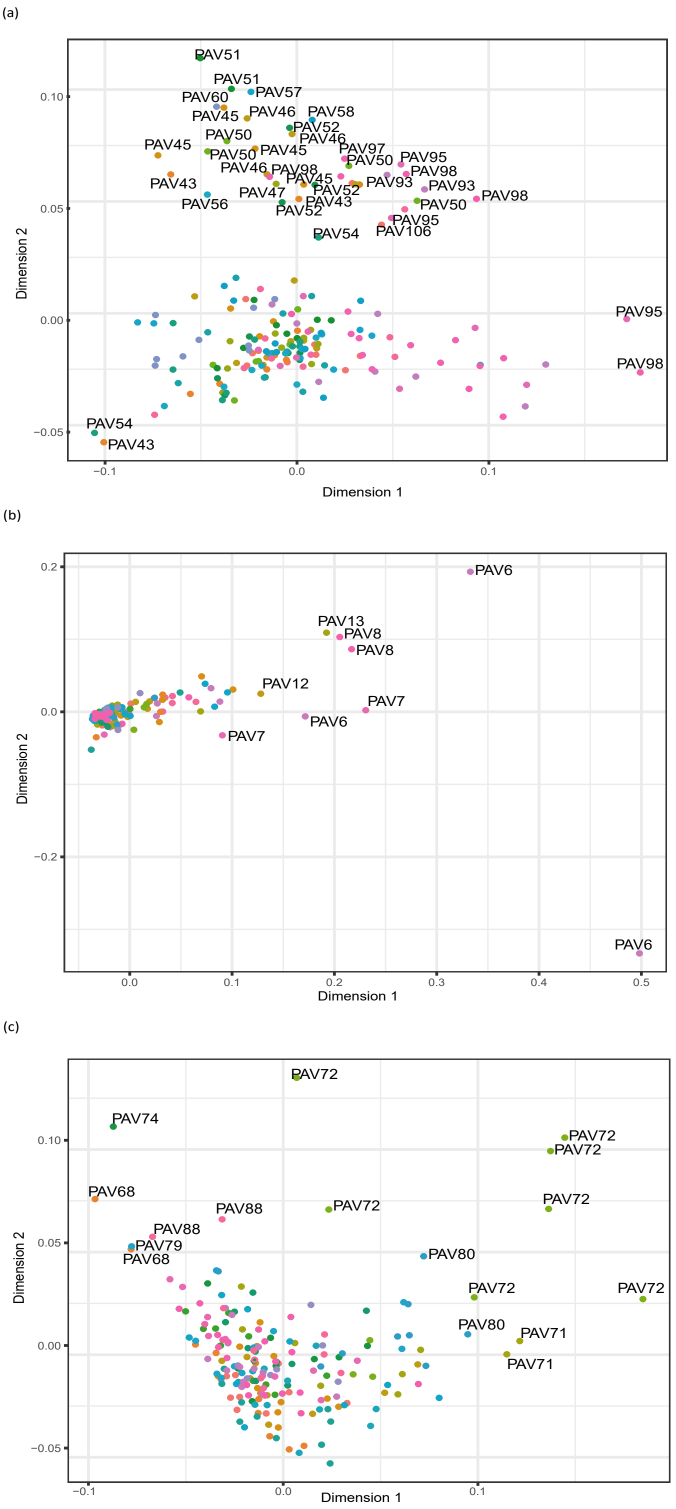


**SI-4 Figure S1**: Non-metric Multidimensional Scaling (NMDS) ordinations based on Jaccard distances derived from binary matrix at SNP loci. Panels show results for (a) *H. atriplicifolium*, (b) *L. stoechas*, and (c) *P. purpurea*. Populations with labels represent those showing greater deviation from the central cloud of points, indicating comparatively higher divergence in missing data patterns.

### SI-5: Nei's distances among populations

(a)

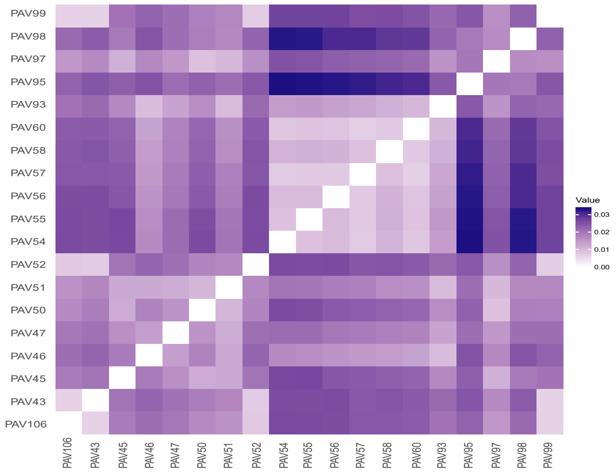


(a)

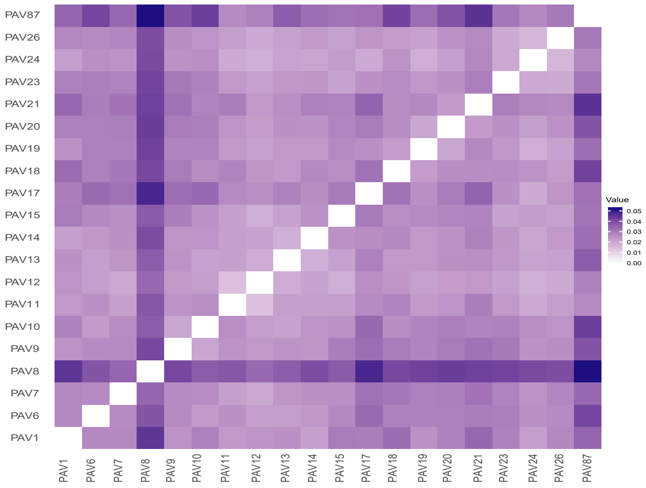


(c)

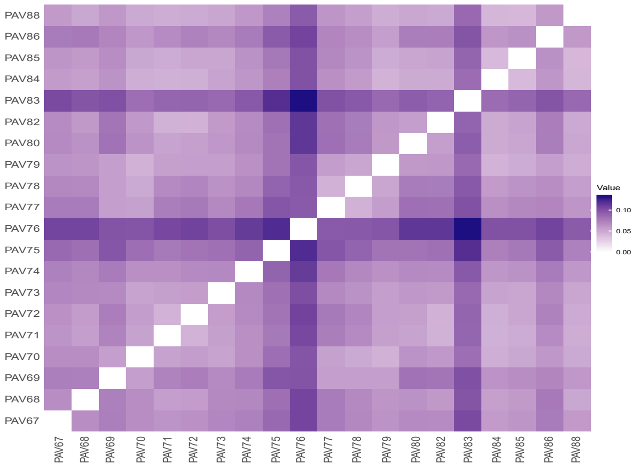


**SI-5 Figure S1**: Heatmaps of Nei's genetic distances among populations, illustrating genomic similarity per species. Darker blue shades indicate greater genomic distances, while lighter shades indicate lower genomic distances (greater similarity). Panels correspond to (a) *H. atriplicifolium*, (b) *L. stoechas*, and (c) *P. purpurea*.

<u>**SI-6: Isolation by distance (IBD) tests**</u>

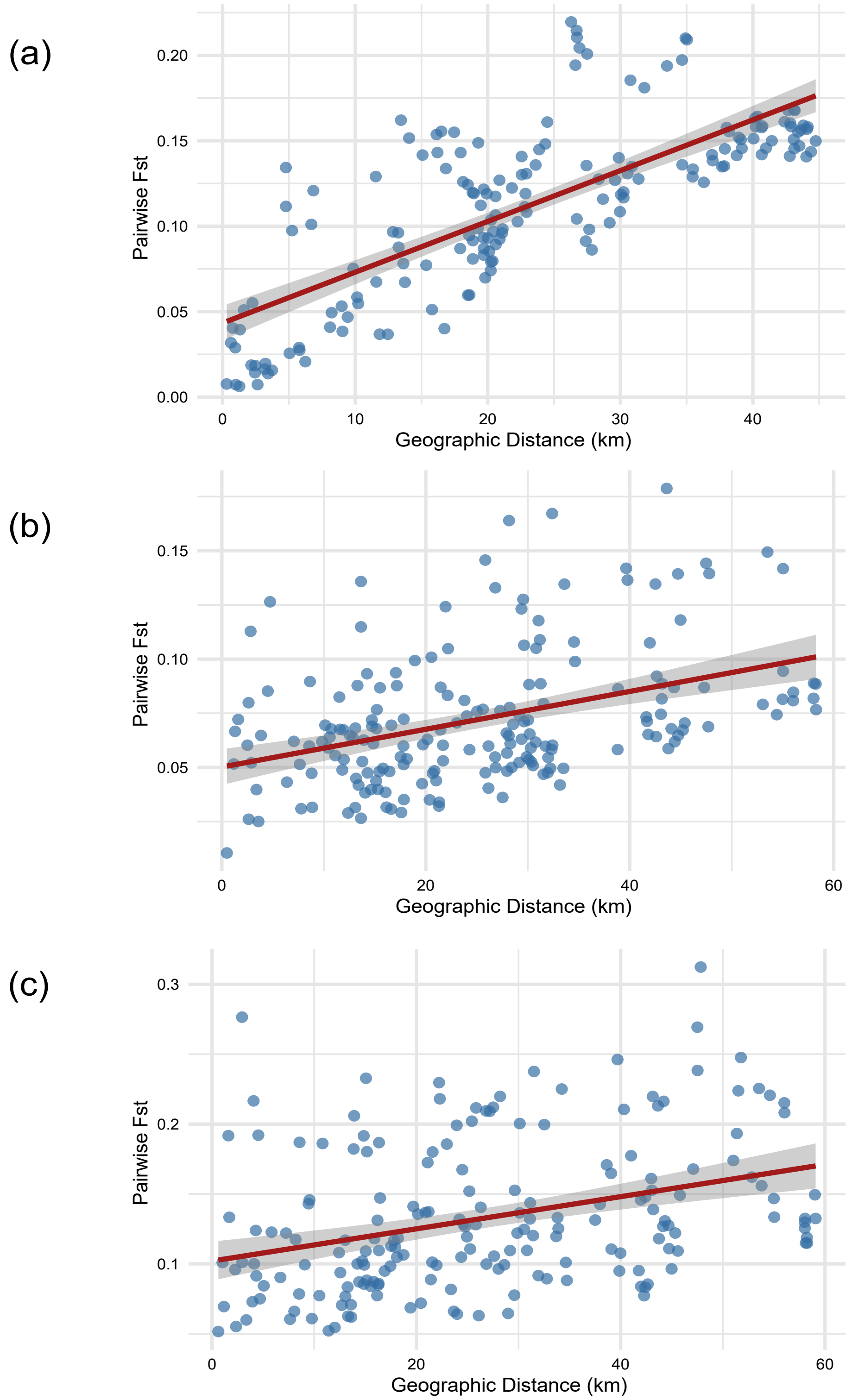


**SI-6 Figure S1**: Relationships between genomic differentiation and geographic distance (IBD) for (a) *H. atriplicifolium*, (b) *L. stoechas*, and (c) *P. purpurea*. Pairwise $F_{st}$ values are plotted against geographic distances between sampling sites. The red lines represent the linear regression fit. The significance of the correlation is assessed using Mantel test with Pearson's correlation method.

### SI-7: Cross-validation errors from ADMIXTURE runs

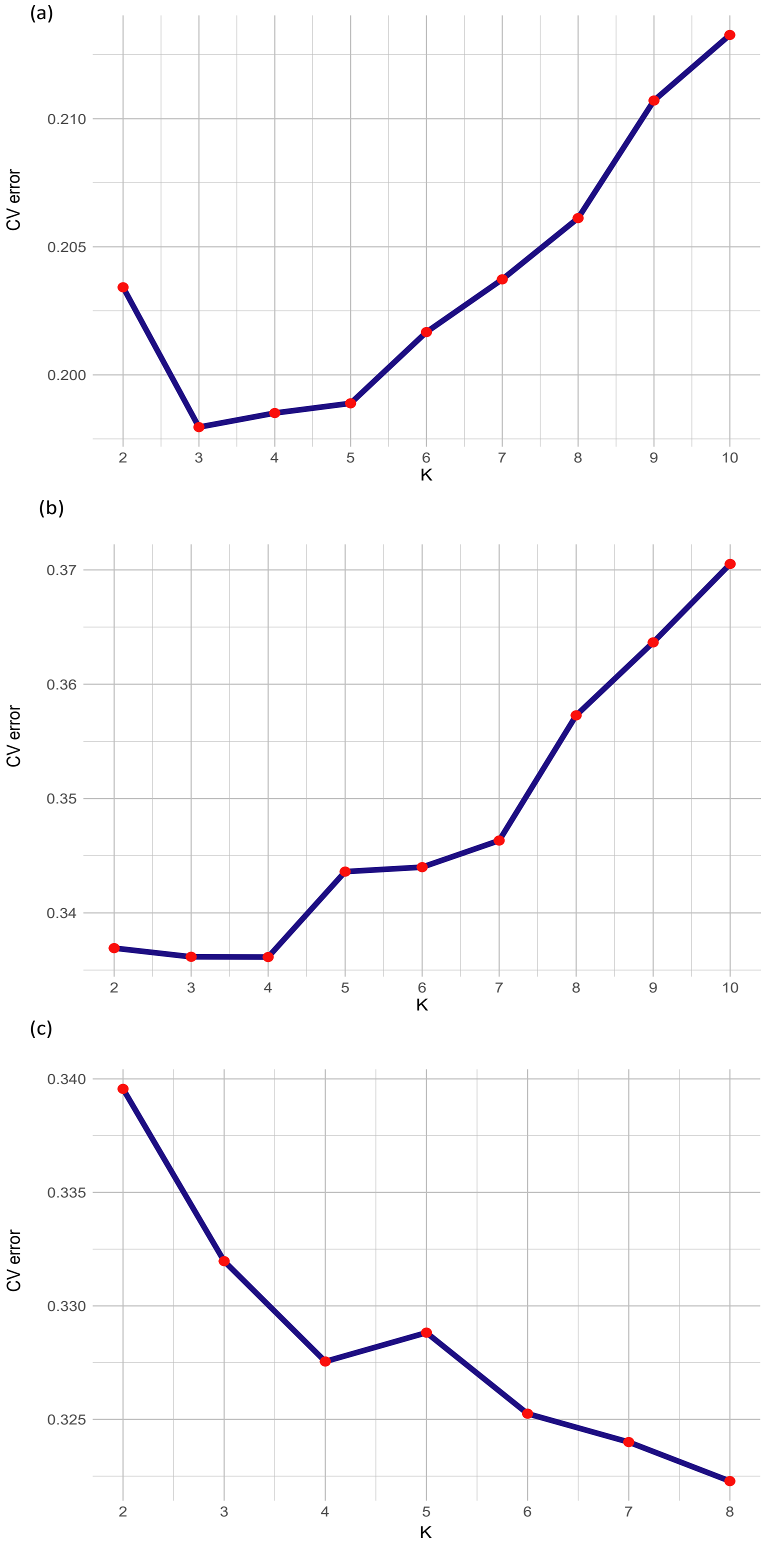


**SI-7 Figure S1**: Cross-validation (CV) error profiles from ADMIXTURE runs for (a) *H. atriplicifolium*, (b) *L. stoechas*, and (c) *P. purpurea*. The optimal K was determined either as the value yielding the lowest CV error or, in cases where CV errors showed a disminishing rate of decrease, as the inflection point, beyond which further increases in K provided negligible improvement in model fit.

### SI-8: Selection of migration edges (m) in TreeMix

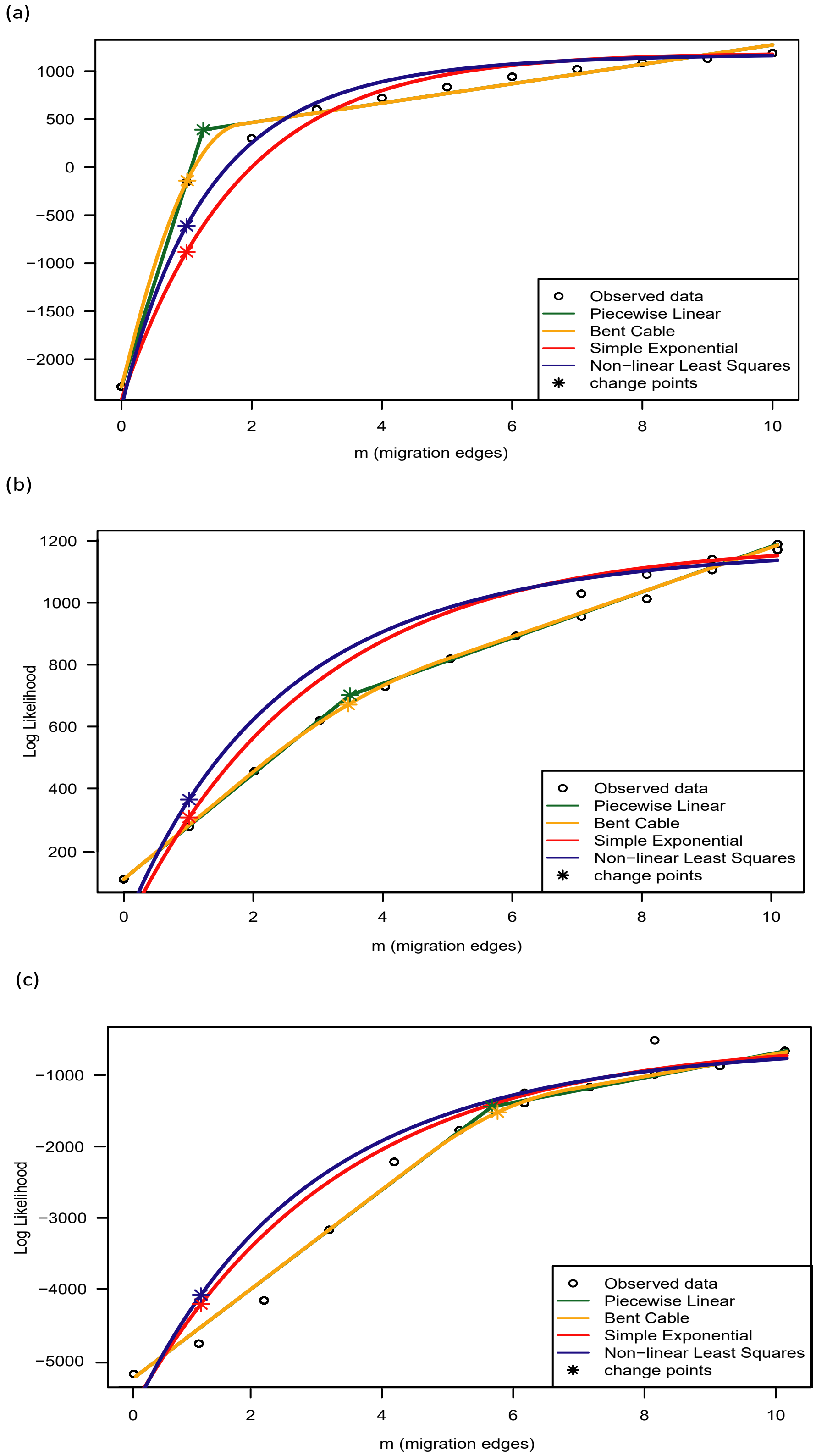


**SI-8 Figure S1**: Determination of the optimal number of migration edges (m) in TreeMix analyses using optM for (a) *H. atriplicifolium*, (b) *L. stoechas*, and (c) *P. purpurea*. Change points in log-likelihood curves indicate the optimal m for each species.
