## Supplementary material for "Independent colonisations of serpentine habitats highlight species-specific evolutionary histories of lineage diversification": SI-1 Table S1

| Identifier | Species | Soil | Acidity (pH - KCl) | Arsenic (As) - (mg/Kg s.m.s.) | Cadmium (Cd) - (mg/Kg s.m.s.) | Copper (Cu) - (mg/Kg s.m.s.) | Chromium (Cr) - (mg/Kg s.m.s.) | Iron (Fe) - (mg/Kg s.m.s.) | Mercury (Hg) - (mg/Kg s.m.s.) | Nickel (Ni) - (mg/Kg s.m.s.) | Lead (Pb) - (mg/Kg s.m.s.) | Zinc (Zn) - (mg/Kg s.m.s.) | Ca/Mg ratio |
|---|---|---|---|---|---|---|---|---|---|---|---|---|---|
| PAV43 | Halimium atriplicifolium | Serpentine | 7.2 | 5 | 0.4 | 14 | 1300 | 78000 | 0.1 | 2600 | 23 | 38 | 1.40 |
| PAV95 | Halimium atriplicifolium | Non-serpentine | 6.8 | 14 | 0.4 | 46 | 110 | 59000 | 0.1 | 120 | 48 | 65 | 36.22 |
| PAV45 | Halimium atriplicifolium | Serpentine | 7.1 | 5 | 0.4 | 23 | 530 | 61000 | 0.1 | 1300 | 10 | 71 | 1.22 |
| PAV97 | Halimium atriplicifolium | Non-serpentine | 7.8 | 5 | 0.4 | 22 | 480 | 39000 | 0.1 | 1300 | 10 | 48 | 3.83 |
| PAV46 | Halimium atriplicifolium | Serpentine | 7.1 | 5 | 0.4 | 22 | 940 | 74000 | 0.1 | 2400 | 10 | 58 | 0.80 |
| PAV93 | Halimium atriplicifolium | Non-serpentine | 7.5 | 5 | 0.4 | 15 | 480 | 37000 | 0.1 | 1300 | 12 | 210 | 2.71 |
| PAV50 | Halimium atriplicifolium | Serpentine | 6.8 | 5 | 0.4 | 14 | 780 | 71000 | 0.1 | 2000 | 10 | 37 | 0.62 |
| PAV98 | Halimium atriplicifolium | Non-serpentine | 8.0 | 5 | 0.4 | 7.6 | 11 | 7500 | 0.1 | 11 | 24 | 34 | 7.20 |
| PAV51 | Halimium atriplicifolium | Serpentine | 6.2 | 5 | 0.4 | 36 | 1100 | 130000 | 0.1 | 3600 | 10 | 42 | 0.47 |
| PAV47 | Halimium atriplicifolium | Non-serpentine | 6.6 | 7.3 | 0.4 | 37 | 75 | 38000 | 0.10 | 72 | 33 | 100 | 7.33 |
| PAV52 | Halimium atriplicifolium | Serpentine | 7.2 | 5 | 0.4 | 11 | 550 | 62000 | 0.1 | 2400 | 10 | 28 | 0.94 |
| PAV57 | Halimium atriplicifolium | Serpentine | 7.0 | 5 | 0.4 | 14 | 320 | 57000 | 0.1 | 2000 | 10 | 41 | 1.59 |
| PAV54 | Halimium atriplicifolium | Non-serpentine | 6.7 | 7.8 | 0.4 | 38 | 56 | 49000 | 0.1 | 75 | 25 | 65 | 5.75 |
| PAV58 | Halimium atriplicifolium | Serpentine | 6.7 | 9.8 | 0.4 | 34 | 860 | 57000 | 0.1 | 1300 | 13 | 62 | 0.65 |
| PAV56 | Halimium atriplicifolium | Non-serpentine | 6.4 | 5 | 0.4 | 31 | 76 | 46000 | 0.1 | 62 | 29 | 73 | 0.46 |
| PAV60 | Halimium atriplicifolium | Serpentine | 7.4 | 5 | 0.4 | 13 | 1300 | 90000 | 0.1 | 3700 | 10 | 38 | 1.00 |
| PAV55 | Halimium atriplicifolium | Non-serpentine | 4.3 | 5 | 0.4 | 12 | 37 | 28000 | 0.1 | 23 | 13 | 64 | 1.20 |
| PAV99 | Halimium atriplicifolium | Serpentine | 6.5 | 5 | 0.4 | 28 | 1500 | 89000 | 0.1 | 2800 | 13 | 45 | 0.73 |
| PAV106 | Halimium atriplicifolium | Non-serpentine | 7.6 | 26 | 0.40 | 20 | 100 | 31000 | 0.11 | 160 | 130 | 160 | 8.74 |
| PAV1 | Lavandula stoechas | Serpentine | 6.9 | 5.1 | 0.4 | 35 | 1500 | 100000 | 0.1 | 3100 | 17 | 76 | 0.52 |
| PAV6 | Lavandula stoechas | Non-Serpentine | 7.5 | 94 | 0.79 | 44 | 160 | 54000 | 0.28 | 170 | 620 | 900 | 4.58 |
| PAV7 | Lavandula stoechas | Serpentine | 7.2 | 5 | 0.4 | 23 | 1100 | 72000 | 0.1 | 2600 | 20 | 40 | 0.52 |
| PAV8 | Lavandula stoechas | Non-Serpentine | 7.5 | 38 | 0.4 | 37 | 23 | 50000 | 0.28 | 20 | 360 | 200 | 4.58 |
| PAV9 | Lavandula stoechas | Serpentine | 7.2 | 7.6 | 0.4 | 33 | 1300 | 86000 | 0.1 | 2600 | 27 | 37 | 0.52 |
| PAV10 | Lavandula stoechas | Non-Serpentine | 7.3 | 33 | 0.4 | 29 | 95 | 34000 | 0.13 | 110 | 18 | 38 | 4.58 |
| PAV11 | Lavandula stoechas | Serpentine | 7.3 | 5 | 0.4 | 12 | 530 | 40000 | 0.1 | 1200 | 10 | 33 | 0.52 |
| PAV12 | Lavandula stoechas | Non-Serpentine | 6.9 | 9.1 | 0.4 | 15 | 330 | 39000 | 0.1 | 750 | 10 | 47 | 4.58 |
| PAV13 | Lavandula stoechas | Non-Serpentine | 6.7 | 5 | 0.4 | 40 | 97 | 49000 | 0.1 | 64 | 10 | 52 | 4.58 |
| PAV14 | Lavandula stoechas | Serpentine | 7.5 | 5 | 0.4 | 29 | 770 | 75000 | 0.1 | 3000 | 10 | 43 | 0.52 |
| PAV15 | Lavandula stoechas | Non-Serpentine | 7.3 | 9.8 | 0.4 | 32 | 170 | 46000 | 0.1 | 230 | 10 | 67 | 3.20 |
| PAV87 | Lavandula stoechas | Serpentine | 6.5 | 5 | 0.4 | 26 | 1200 | 92000 | 0.1 | 3100 | 10 | 49 | 0.29 |
| PAV17 | Lavandula stoechas | Serpentine | 7.4 | 5.3 | 0.4 | 40 | 900 | 77000 | 0.1 | 2900 | 10 | 52 | 0.39 |
| PAV18 | Lavandula stoechas | Non-Serpentine | 7.6 | 11 | 0.4 | 22 | 38 | 34000 | 0.1 | 35 | 19 | 58 | 2.67 |
| PAV19 | Lavandula stoechas | Serpentine | 6.7 | 5 | 0.4 | 20 | 510 | 67000 | 0.1 | 2500 | 10 | 37 | 0.61 |
| PAV20 | Lavandula stoechas | Serpentine | 7.1 | 16 | 0.4 | 32 | 1400 | 87000 | 0.1 | 2900 | 10 | 44 | 0.78 |
| PAV21 | Lavandula stoechas | Non-Serpentine | 7.1 | 5 | 0.4 | 31 | 72 | 38000 | 0.1 | 74 | 10 | 94 | 7.21 |
| PAV23 | Lavandula stoechas | Non-Serpentine | 7.2 | 5.1 | 0.4 | 30 | 68 | 35000 | 0.1 | 96 | 27 | 110 | 5.53 |
| PAV24 | Lavandula stoechas | Serpentine | 7.7 | 5 | 0.4 | 26 | 400 | 54000 | 0.1 | 2000 | 15 | 42 | 0.52 |
| PAV26 | Lavandula stoechas | Non-Serpentine | 5.7 | 5 | 0.4 | 44 | 49 | 34000 | 0.1 | 44 | 27 | 110 | 4.28 |
| PAV67 | Phlomis purpurea | Serpentine | 6.8 | 5 | 0.4 | 40 | 760 | 57000 | 0.1 | 1500 | 10 | 65 | 0.44 |
| PAV74 | Phlomis purpurea | Non-Serpentine | 6.9 | 9.1 | 0.4 | 15 | 330 | 39000 | 0.1 | 750 | 10 | 47 | 3.87 |
| PAV68 | Phlomis purpurea | Serpentine | 7.2 | 5 | 0.4 | 14 | 1300 | 78000 | 0.1 | 2600 | 23 | 38 | 1.40 |
| PAV69 | Phlomis purpurea | Non-Serpentine | 7.2 | 14 | 0.4 | 12 | 64 | 28000 | 0.17 | 81 | 65 | 150 | 4.97 |
| PAV70 | Phlomis purpurea | Non-Serpentine | 6.4 | 5 | 0.4 | 31 | 76 | 46000 | 0.1 | 62 | 29 | 73 | 0.46 |
| PAV71 | Phlomis purpurea | Serpentine | 7.0 | 5 | 0.4 | 14 | 320 | 57000 | 0.1 | 2000 | 10 | 41 | 1.59 |
| PAV72 | Phlomis purpurea | Serpentine | 6.2 | 7.1 | 0.4 | 32 | 1500 | 69000 | 0.14 | 2400 | 10 | 34 | 0.44 |

| | | | | | | | | | | | | | |
|---|---|---|---|---|---|---|---|---|---|---|---|---|---|
| PAV73 | Phlomis purpurea | Non-Serpentine | 6.0 | 5 | 0.4 | 35 | 60 | 37000 | 0.1 | 55 | 10 | 68 | 2.44 |
| PAV75 | Phlomis purpurea | Serpentine | 7.2 | 5 | 0.4 | 23 | 1100 | 72000 | 0.1 | 2600 | 20 | 40 | 0.77 |
| PAV76 | Phlomis purpurea | Non-Serpentine | 7.5 | 38 | 0.4 | 37 | 23 | 50000 | 0.28 | 20 | 360 | 200 | 3.87 |
| PAV77 | Phlomis purpurea | Non-Serpentine | 7.3 | 33 | 0.4 | 29 | 95 | 34000 | 0.13 | 110 | 18 | 38 | 3.87 |
| PAV78 | Phlomis purpurea | Serpentine | 7.2 | 7.6 | 0.4 | 33 | 1300 | 86000 | 0.1 | 2600 | 27 | 37 | 0.77 |
| PAV79 | Phlomis purpurea | Non-Serpentine | 5.9 | 5 | 0.4 | 22 | 56 | 35000 | 0.1 | 44 | 28 | 80 | 6.38 |
| PAV80 | Phlomis purpurea | Serpentine | 7.4 | 5.3 | 0.4 | 40 | 900 | 77000 | 0.1 | 2900 | 10 | 52 | 0.39 |
| PAV82 | Phlomis purpurea | Serpentine | 7.1 | 16 | 0.4 | 32 | 1400 | 87000 | 0.1 | 2900 | 10 | 44 | 0.78 |
| PAV83 | Phlomis purpurea | Non-Serpentine | 7.1 | 5 | 0.4 | 31 | 72 | 38000 | 0.1 | 74 | 10 | 94 | 7.21 |
| PAV84 | Phlomis purpurea | Serpentine | 7.1 | 5 | 0.4 | 22 | 940 | 74000 | 0.1 | 2400 | 10 | 58 | 0.80 |
| PAV85 | Phlomis purpurea | Non-Serpentine | 7.0 | 5 | 0.4 | 18 | 160 | 34000 | 0.1 | 320 | 20 | 52 | 1.75 |
| PAV86 | Phlomis purpurea | Non-Serpentine | 7.9 | 18 | 0.4 | 11 | 13 | 14000 | 0.1 | 14 | 23 | 44 | 3.87 |
| PAV88 | Phlomis purpurea | Serpentine | 6.2 | 5 | 0.4 | 25 | 1400 | 86000 | 0.1 | 2400 | 13 | 56 | 0.32 |
