## Supplementary material for "Independent colonisations of serpentine habitats highlight species-specific evolutionary histories of lineage diversification": SI-3 Table S1

| Sample | Population | Soil | Species | Height (cm) | Weight_g | Weight_mg | Area_mm2 | SLA |
|---|---|---|---|---|---|---|---|---|
| PAV1A | PAV1 | Serpentine | Lavandula stoechas | 33 | | | | |
| PAV1B | PAV1 | Serpentine | Lavandula stoechas | 16 | | | | |
| PAV1C | PAV1 | Serpentine | Lavandula stoechas | 23 | 0.001580 | 1.58 | 19.743 | 12.49557 |
| PAV1D | PAV1 | Serpentine | Lavandula stoechas | 26 | 0.001890 | 1.89 | 21.912 | 11.59365 |
| PAV1E | PAV1 | Serpentine | Lavandula stoechas | 22 | | | | |
| PAV1F | PAV1 | Serpentine | Lavandula stoechas | 39 | | | | |
| PAV1G | PAV1 | Serpentine | Lavandula stoechas | 32 | 0.002890 | 2.89 | 43.041 | 14.89308 |
| PAV1H | PAV1 | Serpentine | Lavandula stoechas | 36 | | | | |
| PAV1I | PAV1 | Serpentine | Lavandula stoechas | 31 | | | | |
| PAV1J | PAV1 | Serpentine | Lavandula stoechas | 33 | | | | |
| PAV1K | PAV1 | Serpentine | Lavandula stoechas | 38 | | | | |
| PAV1L | PAV1 | Serpentine | Lavandula stoechas | 35 | | | | |
| PAV1M | PAV1 | Serpentine | Lavandula stoechas | 26 | | | | |
| PAV1N | PAV1 | Serpentine | Lavandula stoechas | 34 | 0.002770 | 2.77 | 34.171 | 12.33610 |
| PAV1Ñ | PAV1 | Serpentine | Lavandula stoechas | 30 | | | | |
| PAV6A | PAV6 | Nonserpentine | Lavandula stoechas | 37 | | | | |
| PAV6B | PAV6 | Nonserpentine | Lavandula stoechas | 61 | | | | |
| PAV6C | PAV6 | Nonserpentine | Lavandula stoechas | 50 | | | | |
| PAV6D | PAV6 | Nonserpentine | Lavandula stoechas | 69 | | | | |
| PAV6E | PAV6 | Nonserpentine | Lavandula stoechas | 51 | | | | |
| PAV6F | PAV6 | Nonserpentine | Lavandula stoechas | 51 | | | | |
| PAV6G | PAV6 | Nonserpentine | Lavandula stoechas | 44 | | | | |
| PAV6H | PAV6 | Nonserpentine | Lavandula stoechas | 38 | | | | |
| PAV6I | PAV6 | Nonserpentine | Lavandula stoechas | 49 | | | | |
| PAV6J | PAV6 | Nonserpentine | Lavandula stoechas | 29 | | | | |
| PAV6K | PAV6 | Nonserpentine | Lavandula stoechas | 42 | 0.00408 | 4.08 | 53.419 | 13.09289 |
| PAV6L | PAV6 | Nonserpentine | Lavandula stoechas | 45 | 0.00236 | 2.36 | 46.481 | 19.69534 |
| PAV6M | PAV6 | Nonserpentine | Lavandula stoechas | 26 | | | | |
| PAV6N | PAV6 | Nonserpentine | Lavandula stoechas | 60 | | | | |
| PAV6Ñ | PAV6 | Nonserpentine | Lavandula stoechas | 33 | | | | |
| PAV7A | PAV7 | Serpentine | Lavandula stoechas | 25 | | | | |
| PAV7B | PAV7 | Serpentine | Lavandula stoechas | 34 | 0.00544 | 5.44 | 44.477 | 8.17592 |
| PAV7C | PAV7 | Serpentine | Lavandula stoechas | 32 | | | | |
| PAV7D | PAV7 | Serpentine | Lavandula stoechas | 25 | | | | |
| PAV7E | PAV7 | Serpentine | Lavandula stoechas | 15 | | | | |
| PAV7F | PAV7 | Serpentine | Lavandula stoechas | 31 | | | | |
| PAV7G | PAV7 | Serpentine | Lavandula stoechas | 28 | 0.00542 | 5.42 | 47.630 | 8.78782 |
| PAV7H | PAV7 | Serpentine | Lavandula stoechas | 48 | | | | |
| PAV7I | PAV7 | Serpentine | Lavandula stoechas | 23 | 0.00461 | 4.61 | 30.257 | 6.56334 |
| PAV7J | PAV7 | Serpentine | Lavandula stoechas | 27 | 0.00432 | 4.32 | 40.448 | 9.36296 |
| PAV7K | PAV7 | Serpentine | Lavandula stoechas | 25 | | | | |
| PAV7L | PAV7 | Serpentine | Lavandula stoechas | 18 | | | | |
| PAV7M | PAV7 | Serpentine | Lavandula stoechas | 29 | | | | |
| PAV7N | PAV7 | Serpentine | Lavandula stoechas | 14 | | | | |
| PAV7Ñ | PAV7 | Serpentine | Lavandula stoechas | 17 | 0.00399 | 3.99 | 33.281 | 8.34110 |
| PAV8A | PAV8 | Nonserpentine | Lavandula stoechas | 64 | 0.00367 | 3.67 | 28.174 | 7.67684 |
| PAV8B | PAV8 | Nonserpentine | Lavandula stoechas | 63 | 0.00302 | 3.02 | 24.943 | 8.25927 |
| PAV8C | PAV8 | Nonserpentine | Lavandula stoechas | 73 | 0.00294 | 2.94 | 24.066 | 8.18571 |
| PAV8D | PAV8 | Nonserpentine | Lavandula stoechas | 51 | 0.00372 | 3.72 | 29.977 | 8.05833 |
| PAV8E | PAV8 | Nonserpentine | Lavandula stoechas | 64 | 0.00731 | 7.31 | 54.244 | 7.42052 |
| PAV8F | PAV8 | Nonserpentine | Lavandula stoechas | 22 | | | | |
| PAV8G | PAV8 | Nonserpentine | Lavandula stoechas | 82 | 0.00444 | 4.44 | 32.268 | 7.26757 |
| PAV8H | PAV8 | Nonserpentine | Lavandula stoechas | 26 | 0.00177 | 1.77 | 17.351 | 9.80282 |
| PAV8I | PAV8 | Nonserpentine | Lavandula stoechas | 59 | 0.0069 | 6.9 | 61.498 | 8.91275 |
| PAV8J | PAV8 | Nonserpentine | Lavandula stoechas | 56 | 0.00436 | 4.36 | 50.302 | 11.53716 |
| PAV8K | PAV8 | Nonserpentine | Lavandula stoechas | 28 | 0.00327 | 3.27 | 26.142 | 7.99450 |
| PAV8L | PAV8 | Nonserpentine | Lavandula stoechas | 56 | | | | |
| PAV8M | PAV8 | Nonserpentine | Lavandula stoechas | 45 | | | | |
| PAV8N | PAV8 | Nonserpentine | Lavandula stoechas | 37 | 0.005242 | 5.242 | 52.930 | 10.09729 |
| PAV8Ñ | PAV8 | Nonserpentine | Lavandula stoechas | 19 | 0.00408 | 4.08 | 51.537 | 12.63162 |
| PAV9A | PAV9 | Serpentine | Lavandula stoechas | 48 | 0.00486 | 4.86 | 35.284 | 7.26008 |
| PAV9B | PAV9 | Serpentine | Lavandula stoechas | 46 | | | | |
| PAV9C | PAV9 | Serpentine | Lavandula stoechas | 58 | 0.00391 | 3.91 | 30.415 | 7.77877 |
| PAV9D | PAV9 | Serpentine | Lavandula stoechas | 55 | | | | |
| PAV9E | PAV9 | Serpentine | Lavandula stoechas | 63 | 0.00252 | 2.52 | 12.317 | 4.88770 |
| PAV9F | PAV9 | Serpentine | Lavandula stoechas | 68 | 0.002 | 2 | 14.270 | 7.13500 |
| PAV9G | PAV9 | Serpentine | Lavandula stoechas | 61 | | | | |
| PAV9H | PAV9 | Serpentine | Lavandula stoechas | 57 | 0.00248 | 2.48 | 13.617 | 5.49073 |
| PAV9I | PAV9 | Serpentine | Lavandula stoechas | 80 | 0.00343 | 3.43 | 20.346 | 5.93178 |
| PAV9J | PAV9 | Serpentine | Lavandula stoechas | 92 | 0.00423 | 4.23 | 31.600 | 7.47045 |
| PAV9K | PAV9 | Serpentine | Lavandula stoechas | 31 | | | | |
| PAV9L | PAV9 | Serpentine | Lavandula stoechas | 44 | 0.00191 | 1.91 | 16.403 | 8.58796 |
| PAV9M | PAV9 | Serpentine | Lavandula stoechas | 35 | 0.00174 | 1.74 | 12.841 | 7.37989 |
| PAV9N | PAV9 | Serpentine | Lavandula stoechas | 72 | 0.00342 | 3.42 | 23.118 | 6.75965 |
| PAV9Ñ | PAV9 | Serpentine | Lavandula stoechas | 74 | | | | |
| PAV10A | PAV10 | Nonserpentine | Lavandula stoechas | 77 | | | | |
| PAV10B | PAV10 | Nonserpentine | Lavandula stoechas | 53 | | | | |
| PAV10C | PAV10 | Nonserpentine | Lavandula stoechas | 64 | | | | |
| PAV10D | PAV10 | Nonserpentine | Lavandula stoechas | 79 | | | | |
| PAV10E | PAV10 | Nonserpentine | Lavandula stoechas | 60 | | | | |
| PAV10F | PAV10 | Nonserpentine | Lavandula stoechas | 75 | | | | |
| PAV10G | PAV10 | Nonserpentine | Lavandula stoechas | 88 | | | | |
| PAV10H | PAV10 | Nonserpentine | Lavandula stoechas | 83 | | | | |
| PAV10I | PAV10 | Nonserpentine | Lavandula stoechas | 94 | | | | |
| PAV10J | PAV10 | Nonserpentine | Lavandula stoechas | 75 | | | | |
| PAV10K | PAV10 | Nonserpentine | Lavandula stoechas | 79 | | | | |
| PAV10L | PAV10 | Nonserpentine | Lavandula stoechas | 49 | | | | |
| PAV10M | PAV10 | Nonserpentine | Lavandula stoechas | 64 | | | | |
| PAV10N | PAV10 | Nonserpentine | Lavandula stoechas | 57 | | | | |

| | | | | | | | | |
|---|---|---|---|---|---|---|---|---|
| PAV10Ñ | PAV10 | Nonserpentine | Lavandula stoechas | 74 | | | | |
| PAV11A | PAV11 | Serpentine | Lavandula stoechas | 26 | 0.00277 | 2.77 | 16.317 | 5.89061 |
| PAV11B | PAV11 | Serpentine | Lavandula stoechas | 20 | 0.00356 | 3.56 | 29.230 | 8.21067 |
| PAV11C | PAV11 | Serpentine | Lavandula stoechas | 19 | | | | |
| PAV11D | PAV11 | Serpentine | Lavandula stoechas | 41 | | | | |
| PAV11E | PAV11 | Serpentine | Lavandula stoechas | 35 | 0.00597 | 5.97 | 44.010 | 7.37186 |
| PAV11F | PAV11 | Serpentine | Lavandula stoechas | 34 | | | | |
| PAV11G | PAV11 | Serpentine | Lavandula stoechas | 30 | | | | |
| PAV11H | PAV11 | Serpentine | Lavandula stoechas | 21 | | | | |
| PAV11I | PAV11 | Serpentine | Lavandula stoechas | 38 | 0.00487 | 4.87 | 43.177 | 8.86591 |
| PAV11J | PAV11 | Serpentine | Lavandula stoechas | 54 | 0.00317 | 3.17 | 22.824 | 7.20000 |
| PAV11K | PAV11 | Serpentine | Lavandula stoechas | 28 | 0.00242 | 2.42 | 24.720 | 10.21488 |
| PAV11L | PAV11 | Serpentine | Lavandula stoechas | 32 | | | | |
| PAV11M | PAV11 | Serpentine | Lavandula stoechas | 29 | 0.00558 | 5.58 | 36.699 | 6.57688 |
| PAV11N | PAV11 | Serpentine | Lavandula stoechas | 22 | | | | |
| PAV11Ñ | PAV11 | Serpentine | Lavandula stoechas | 33 | | | | |
| PAV12A | PAV12 | Nonserpentine | Lavandula stoechas | 87 | | | | |
| PAV12B | PAV12 | Nonserpentine | Lavandula stoechas | 86 | | | | |
| PAV12C | PAV12 | Nonserpentine | Lavandula stoechas | 62 | | | | |
| PAV12D | PAV12 | Nonserpentine | Lavandula stoechas | 48 | | | | |
| PAV12E | PAV12 | Nonserpentine | Lavandula stoechas | 130 | | | | |
| PAV12F | PAV12 | Nonserpentine | Lavandula stoechas | 75 | | | | |
| PAV12G | PAV12 | Nonserpentine | Lavandula stoechas | 35 | | | | |
| PAV12H | PAV12 | Nonserpentine | Lavandula stoechas | 68 | | | | |
| PAV12I | PAV12 | Nonserpentine | Lavandula stoechas | 96 | | | | |
| PAV12J | PAV12 | Nonserpentine | Lavandula stoechas | 60 | | | | |
| PAV12K | PAV12 | Nonserpentine | Lavandula stoechas | 40 | | | | |
| PAV12L | PAV12 | Nonserpentine | Lavandula stoechas | 53 | | | | |
| PAV12M | PAV12 | Nonserpentine | Lavandula stoechas | 81 | | | | |
| PAV12N | PAV12 | Nonserpentine | Lavandula stoechas | 37 | | | | |
| PAV12Ñ | PAV12 | Nonserpentine | Lavandula stoechas | 91 | | | | |
| PAV13A | PAV13 | Nonserpentine | Lavandula stoechas | 58 | | | | |
| PAV13B | PAV13 | Nonserpentine | Lavandula stoechas | 51 | | | | |
| PAV13C | PAV13 | Nonserpentine | Lavandula stoechas | 49 | 0.00145 | 1.45 | 7.419 | 5.11655 |
| PAV13D | PAV13 | Nonserpentine | Lavandula stoechas | 20 | 0.0028 | 2.8 | 13.121 | 4.68607 |
| PAV13E | PAV13 | Nonserpentine | Lavandula stoechas | 47 | 0.00178 | 1.78 | 12.302 | 6.91124 |
| PAV13F | PAV13 | Nonserpentine | Lavandula stoechas | 54 | 0.00246 | 2.46 | 18.831 | 7.65488 |
| PAV13G | PAV13 | Nonserpentine | Lavandula stoechas | 61 | | | | |
| PAV13H | PAV13 | Nonserpentine | Lavandula stoechas | 71 | | | | |
| PAV13I | PAV13 | Nonserpentine | Lavandula stoechas | 22 | 0.00191 | 1.91 | 12.381 | 6.48220 |
| PAV13J | PAV13 | Nonserpentine | Lavandula stoechas | 35 | 0.00271 | 2.71 | 17.373 | 6.41070 |
| PAV13K | PAV13 | Nonserpentine | Lavandula stoechas | 47 | | | | |
| PAV13L | PAV13 | Nonserpentine | Lavandula stoechas | 37 | | | | |
| PAV13M | PAV13 | Nonserpentine | Lavandula stoechas | 46 | 0.00289 | 2.89 | 16.590 | 5.74048 |
| PAV13N | PAV13 | Nonserpentine | Lavandula stoechas | 55 | 0.00178 | 1.78 | 16.224 | 9.11461 |
| PAV13Ñ | PAV13 | Nonserpentine | Lavandula stoechas | 42 | 0.00269 | 2.69 | 17.244 | 6.41041 |
| PAV14A | PAV14 | Serpentine | Lavandula stoechas | 62 | | | | |
| PAV14B | PAV14 | Serpentine | Lavandula stoechas | 50 | | | | |
| PAV14C | PAV14 | Serpentine | Lavandula stoechas | 62 | 0.00262 | 2.62 | 22.529 | 8.59885 |
| PAV14D | PAV14 | Serpentine | Lavandula stoechas | 54 | 0.0021 | 2.1 | 22.407 | 10.67000 |
| PAV14E | PAV14 | Serpentine | Lavandula stoechas | 19 | 0.00363 | 3.63 | 44.413 | 12.23499 |
| PAV14F | PAV14 | Serpentine | Lavandula stoechas | 38 | | | | |
| PAV14G | PAV14 | Serpentine | Lavandula stoechas | 48 | | | | |
| PAV14H | PAV14 | Serpentine | Lavandula stoechas | 66 | | | | |
| PAV14I | PAV14 | Serpentine | Lavandula stoechas | 46 | 0.0033 | 3.3 | 36.484 | 11.05576 |
| PAV14J | PAV14 | Serpentine | Lavandula stoechas | 60 | 0.00263 | 2.63 | 24.418 | 9.28441 |
| PAV14K | PAV14 | Serpentine | Lavandula stoechas | 33 | 0.00322 | 3.22 | 40.822 | 12.67764 |
| PAV14L | PAV14 | Serpentine | Lavandula stoechas | 65 | | | | |
| PAV14M | PAV14 | Serpentine | Lavandula stoechas | 39 | 0.00373 | 3.73 | 43.666 | 11.70670 |
| PAV14N | PAV14 | Serpentine | Lavandula stoechas | 26 | 0.00406 | 4.06 | 37.238 | 9.17192 |
| PAV14Ñ | PAV14 | Serpentine | Lavandula stoechas | 53 | | | | |
| PAV15A | PAV15 | Nonserpentine | Lavandula stoechas | 69 | 0.00125 | 1.25 | 10.019 | 8.01520 |
| PAV15B | PAV15 | Nonserpentine | Lavandula stoechas | 84 | | | | |
| PAV15C | PAV15 | Nonserpentine | Lavandula stoechas | 42 | | | | |
| PAV15D | PAV15 | Nonserpentine | Lavandula stoechas | 47 | 0.00243 | 2.43 | 11.979 | 4.92963 |
| PAV15E | PAV15 | Nonserpentine | Lavandula stoechas | 50 | 0.00368 | 3.68 | 20.756 | 5.64022 |
| PAV15F | PAV15 | Nonserpentine | Lavandula stoechas | 60 | | | | |
| PAV15G | PAV15 | Nonserpentine | Lavandula stoechas | 102 | 0.00251 | 2.51 | 18.594 | 7.40797 |
| PAV15H | PAV15 | Nonserpentine | Lavandula stoechas | 92 | | | | |
| PAV15I | PAV15 | Nonserpentine | Lavandula stoechas | 78 | 0.00256 | 2.56 | 13.078 | 5.10859 |
| PAV15J | PAV15 | Nonserpentine | Lavandula stoechas | 81 | 0.00203 | 2.03 | 3.124 | 1.53892 |
| PAV15K | PAV15 | Nonserpentine | Lavandula stoechas | 55 | | | | |
| PAV15L | PAV15 | Nonserpentine | Lavandula stoechas | 37 | 0.00229 | 2.29 | 18.594 | 8.11965 |
| PAV15M | PAV15 | Nonserpentine | Lavandula stoechas | 52 | | | | |
| PAV15N | PAV15 | Nonserpentine | Lavandula stoechas | 40 | 0.00319 | 3.19 | 21.402 | 6.70909 |
| PAV15Ñ | PAV15 | Nonserpentine | Lavandula stoechas | 55 | 0.0022 | 2.2 | 12.870 | 5.85000 |
| PAV17A | PAV17 | Serpentine | Lavandula stoechas | 17 | 0.00196 | 1.96 | 12.690 | 6.47449 |
| PAV17B | PAV17 | Serpentine | Lavandula stoechas | 99 | | | | |
| PAV17C | PAV17 | Serpentine | Lavandula stoechas | 61 | 0.00391 | 3.91 | 22.881 | 5.85192 |
| PAV17D | PAV17 | Serpentine | Lavandula stoechas | 22 | 0.0019 | 1.9 | 8.668 | 4.56211 |
| PAV17E | PAV17 | Serpentine | Lavandula stoechas | 32 | 0.00362 | 3.62 | 20.562 | 5.68011 |
| PAV17F | PAV17 | Serpentine | Lavandula stoechas | 52 | 0.00252 | 2.52 | 16.992 | 6.74286 |
| PAV17G | PAV17 | Serpentine | Lavandula stoechas | 49 | | | | |
| PAV17H | PAV17 | Serpentine | Lavandula stoechas | 51 | | | | |
| PAV17I | PAV17 | Serpentine | Lavandula stoechas | 31 | | | | |
| PAV17J | PAV17 | Serpentine | Lavandula stoechas | 72 | 0.00242 | 2.42 | 13.897 | 5.74256 |
| PAV17K | PAV17 | Serpentine | Lavandula stoechas | 22 | | | | |
| PAV17L | PAV17 | Serpentine | Lavandula stoechas | 96 | 0.00226 | 2.26 | 11.570 | 5.11947 |
| PAV17M | PAV17 | Serpentine | Lavandula stoechas | 83 | 0.00339 | 3.39 | 14.486 | 4.27316 |
| PAV17N | PAV17 | Serpentine | Lavandula stoechas | 62 | 0.00553 | 5.53 | 29.388 | 5.31429 |

| | | | | | | | | |
|---|---|---|---|---|---|---|---|---|
| PAV17Ñ | PAV17 | Serpentine | *Lavandula stoechas* | 119 | 0.00559 | 5.59 | 26.386 | 4.72021 |
| PAV18A | PAV18 | Nonserpentine | *Lavandula stoechas* | 42 | | | | |
| PAV18B | PAV18 | Nonserpentine | *Lavandula stoechas* | 59 | 0.00259 | 2.59 | 25.589 | 9.87992 |
| PAV18C | PAV18 | Nonserpentine | *Lavandula stoechas* | 59 | | | | |
| PAV18D | PAV18 | Nonserpentine | *Lavandula stoechas* | 66 | 0.0027 | 2.7 | 27.061 | 10.02259 |
| PAV18E | PAV18 | Nonserpentine | *Lavandula stoechas* | 37 | | | | |
| PAV18F | PAV18 | Nonserpentine | *Lavandula stoechas* | 26 | | | | |
| PAV18G | PAV18 | Nonserpentine | *Lavandula stoechas* | 40 | | | | |
| PAV18H | PAV18 | Nonserpentine | *Lavandula stoechas* | 36 | | | | |
| PAV18I | PAV18 | Nonserpentine | *Lavandula stoechas* | 60 | | | | |
| PAV18J | PAV18 | Nonserpentine | *Lavandula stoechas* | 75 | | | | |
| PAV18K | PAV18 | Nonserpentine | *Lavandula stoechas* | 63 | | | | |
| PAV18L | PAV18 | Nonserpentine | *Lavandula stoechas* | 52 | | | | |
| PAV18M | PAV18 | Nonserpentine | *Lavandula stoechas* | 53 | | | | |
| PAV18N | PAV18 | Nonserpentine | *Lavandula stoechas* | 46 | 0.00339 | 3.39 | 24.167 | 7.12891 |
| PAV18Ñ | PAV18 | Nonserpentine | *Lavandula stoechas* | 40 | 0.00235 | 2.35 | 16.813 | 7.15447 |
| PAV19A | PAV19 | Serpentine | *Lavandula stoechas* | 28 | | | | |
| PAV19B | PAV19 | Serpentine | *Lavandula stoechas* | 31 | 0.00373 | 3.73 | 20.906 | 5.60483 |
| PAV19C | PAV19 | Serpentine | *Lavandula stoechas* | 73 | 0.00574 | 5.74 | 54.173 | 9.43780 |
| PAV19D | PAV19 | Serpentine | *Lavandula stoechas* | 60 | | | | |
| PAV19E | PAV19 | Serpentine | *Lavandula stoechas* | 41 | 0.00372 | 3.72 | 32.318 | 8.68763 |
| PAV19F | PAV19 | Serpentine | *Lavandula stoechas* | 32 | 0.00736 | 7.36 | 66.504 | 9.03587 |
| PAV19G | PAV19 | Serpentine | *Lavandula stoechas* | 34 | | | | |
| PAV19H | PAV19 | Serpentine | *Lavandula stoechas* | 59 | 0.00169 | 1.69 | 18.500 | 10.94675 |
| PAV19I | PAV19 | Serpentine | *Lavandula stoechas* | 36 | 0.00483 | 4.83 | 49.943 | 10.34017 |
| PAV19J | PAV19 | Serpentine | *Lavandula stoechas* | 47 | 0.00659 | 6.59 | 49.943 | 7.57860 |
| PAV19K | PAV19 | Serpentine | *Lavandula stoechas* | 57 | 0.00733 | 7.33 | 62.554 | 8.53397 |
| PAV19L | PAV19 | Serpentine | *Lavandula stoechas* | 50 | | | | |
| PAV19M | PAV19 | Serpentine | *Lavandula stoechas* | 49 | 0.00432 | 4.32 | 31.622 | 7.31991 |
| PAV19N | PAV19 | Serpentine | *Lavandula stoechas* | 29 | 0.00312 | 3.12 | 28.275 | 9.06250 |
| PAV19Ñ | PAV19 | Serpentine | *Lavandula stoechas* | 57 | 0.00598 | 5.98 | 49.641 | 8.30117 |
| PAV20A | PAV20 | Serpentine | *Lavandula stoechas* | 37 | 0.00577 | 5.77 | 58.597 | 10.15546 |
| PAV20B | PAV20 | Serpentine | *Lavandula stoechas* | 46 | | | | |
| PAV20C | PAV20 | Serpentine | *Lavandula stoechas* | 39 | | | | |
| PAV20D | PAV20 | Serpentine | *Lavandula stoechas* | 31 | | | | |
| PAV20E | PAV20 | Serpentine | *Lavandula stoechas* | 21 | | | | |
| PAV20F | PAV20 | Serpentine | *Lavandula stoechas* | 24 | | | | |
| PAV20G | PAV20 | Serpentine | *Lavandula stoechas* | 18 | 0.00607 | 6.07 | 47.264 | 7.78649 |
| PAV20H | PAV20 | Serpentine | *Lavandula stoechas* | 33 | | | | |
| PAV20I | PAV20 | Serpentine | *Lavandula stoechas* | 45 | | | | |
| PAV20J | PAV20 | Serpentine | *Lavandula stoechas* | 47 | | | | |
| PAV20K | PAV20 | Serpentine | *Lavandula stoechas* | 20 | | | | |
| PAV20L | PAV20 | Serpentine | *Lavandula stoechas* | 36 | | | | |
| PAV20M | PAV20 | Serpentine | *Lavandula stoechas* | 46 | | | | |
| PAV20N | PAV20 | Serpentine | *Lavandula stoechas* | 32 | | | | |
| PAV20Ñ | PAV20 | Serpentine | *Lavandula stoechas* | 40 | | | | |
| PAV21A | PAV21 | Nonserpentine | *Lavandula stoechas* | 57 | | | | |
| PAV21B | PAV21 | Nonserpentine | *Lavandula stoechas* | 62 | | | | |
| PAV21C | PAV21 | Nonserpentine | *Lavandula stoechas* | 42 | | | | |
| PAV21D | PAV21 | Nonserpentine | *Lavandula stoechas* | 23 | | | | |
| PAV21E | PAV21 | Nonserpentine | *Lavandula stoechas* | 33 | 0.00428 | 4.28 | 38.495 | 8.99416 |
| PAV21F | PAV21 | Nonserpentine | *Lavandula stoechas* | 103 | | | | |
| PAV21G | PAV21 | Nonserpentine | *Lavandula stoechas* | 57 | | | | |
| PAV21H | PAV21 | Nonserpentine | *Lavandula stoechas* | 26 | | | | |
| PAV21I | PAV21 | Nonserpentine | *Lavandula stoechas* | 71 | | | | |
| PAV21J | PAV21 | Nonserpentine | *Lavandula stoechas* | 24 | | | | |
| PAV21K | PAV21 | Nonserpentine | *Lavandula stoechas* | 45 | | | | |
| PAV21L | PAV21 | Nonserpentine | *Lavandula stoechas* | 29 | | | | |
| PAV21M | PAV21 | Nonserpentine | *Lavandula stoechas* | 47 | | | | |
| PAV21N | PAV21 | Nonserpentine | *Lavandula stoechas* | 22 | | | | |
| PAV21Ñ | PAV21 | Nonserpentine | *Lavandula stoechas* | 81 | | | | |
| PAV23A | PAV23 | Nonserpentine | *Lavandula stoechas* | 112 | 0.00478 | 4.78 | 53.165 | 11.12238 |
| PAV23B | PAV23 | Nonserpentine | *Lavandula stoechas* | 86 | 0.0035 | 3.5 | 46.871 | 13.39171 |
| PAV23C | PAV23 | Nonserpentine | *Lavandula stoechas* | 88 | | | | |
| PAV23D | PAV23 | Nonserpentine | *Lavandula stoechas* | 104 | | | | |
| PAV23E | PAV23 | Nonserpentine | *Lavandula stoechas* | 94 | 0.00342 | 3.42 | 38.512 | 11.26082 |
| PAV23F | PAV23 | Nonserpentine | *Lavandula stoechas* | 115 | | | | |
| PAV23G | PAV23 | Nonserpentine | *Lavandula stoechas* | 45 | | | | |
| PAV23H | PAV23 | Nonserpentine | *Lavandula stoechas* | 52 | | | | |
| PAV23I | PAV23 | Nonserpentine | *Lavandula stoechas* | 18 | | | | |
| PAV23J | PAV23 | Nonserpentine | *Lavandula stoechas* | 59 | | | | |
| PAV23K | PAV23 | Nonserpentine | *Lavandula stoechas* | 51 | 0.00419 | 4.19 | 31.794 | 7.58807 |
| PAV23L | PAV23 | Nonserpentine | *Lavandula stoechas* | 65 | 0.00267 | 2.67 | 15.333 | 5.74270 |
| PAV23M | PAV23 | Nonserpentine | *Lavandula stoechas* | 58 | 0.00278 | 2.78 | 17.509 | 6.29820 |
| PAV23N | PAV23 | Nonserpentine | *Lavandula stoechas* | 128 | 0.00261 | 2.61 | 17.732 | 6.79387 |
| PAV23Ñ | PAV23 | Nonserpentine | *Lavandula stoechas* | 42 | 0.00258 | 2.58 | 17.438 | 6.75891 |
| PAV24A | PAV24 | Serpentine | *Lavandula stoechas* | 89 | 0.00512 | 5.12 | 37.058 | 7.23789 |
| PAV24B | PAV24 | Serpentine | *Lavandula stoechas* | 75 | | | | |
| PAV24C | PAV24 | Serpentine | *Lavandula stoechas* | 88 | | | | |
| PAV24D | PAV24 | Serpentine | *Lavandula stoechas* | 56 | 0.00304 | 3.04 | 20.296 | 6.67632 |
| PAV24E | PAV24 | Serpentine | *Lavandula stoechas* | 94 | 0.00161 | 1.61 | 15.513 | 9.63540 |
| PAV24F | PAV24 | Serpentine | *Lavandula stoechas* | 78 | 0.00224 | 2.24 | 13.258 | 5.91875 |
| PAV24G | PAV24 | Serpentine | *Lavandula stoechas* | 63 | 0.00213 | 2.13 | 18.357 | 8.61831 |
| PAV24H | PAV24 | Serpentine | *Lavandula stoechas* | 79 | 0.0021 | 2.1 | 11.893 | 5.66333 |
| PAV24I | PAV24 | Serpentine | *Lavandula stoechas* | 59 | | | | |
| PAV24J | PAV24 | Serpentine | *Lavandula stoechas* | 42 | 0.00128 | 1.28 | 11.498 | 8.98281 |
| PAV24K | PAV24 | Serpentine | *Lavandula stoechas* | 57 | 0.00236 | 2.36 | 13.818 | 5.85508 |
| PAV24L | PAV24 | Serpentine | *Lavandula stoechas* | 35 | 0.00238 | 2.38 | 23.492 | 9.87059 |
| PAV24M | PAV24 | Serpentine | *Lavandula stoechas* | 51 | 0.0027 | 2.7 | 19.255 | 7.13148 |
| PAV24N | PAV24 | Serpentine | *Lavandula stoechas* | 38 | 0.00231 | 2.31 | 14.536 | 6.29264 |

| | | | | | | | | |
|---|---|---|---|---|---|---|---|---|
| PAV24Ñ | PAV24 | Serpentine | Lavandula stoechas | 37 | | | | |
| PAV26A | PAV26 | Nonserpentine | Lavandula stoechas | 61 | | | | |
| PAV26B | PAV26 | Nonserpentine | Lavandula stoechas | 68 | | | | |
| PAV26C | PAV26 | Nonserpentine | Lavandula stoechas | 98 | | | | |
| PAV26D | PAV26 | Nonserpentine | Lavandula stoechas | 88 | | | | |
| PAV26E | PAV26 | Nonserpentine | Lavandula stoechas | 72 | 0.00762 | 7.62 | 100.119 | 13.13898 |
| PAV26F | PAV26 | Nonserpentine | Lavandula stoechas | 67 | 0.00672 | 6.72 | 85.834 | 12.77292 |
| PAV26G | PAV26 | Nonserpentine | Lavandula stoechas | 51 | 0.00762 | 7.62 | 100.119 | 13.13898 |
| PAV26H | PAV26 | Nonserpentine | Lavandula stoechas | 58 | 0.00626 | 6.26 | 78.532 | 12.54505 |
| PAV26I | PAV26 | Nonserpentine | Lavandula stoechas | 59 | | | | |
| PAV26J | PAV26 | Nonserpentine | Lavandula stoechas | 76 | 0.00825 | 8.25 | 110.119 | 13.34776 |
| PAV26K | PAV26 | Nonserpentine | Lavandula stoechas | 65 | 0.00453 | 4.53 | 51.072 | 11.27417 |
| PAV26L | PAV26 | Nonserpentine | Lavandula stoechas | 67 | 0.00727 | 7.27 | 94.564 | 13.00743 |
| PAV26M | PAV26 | Nonserpentine | Lavandula stoechas | 49 | 0.00534 | 5.34 | 63.929 | 11.97172 |
| PAV26N | PAV26 | Nonserpentine | Lavandula stoechas | 68 | | | | |
| PAV26Ñ | PAV26 | Nonserpentine | Lavandula stoechas | 51 | | | | |
| PAV87A | PAV87 | Serpentine | Lavandula stoechas | 54 | 0.00392 | 3.92 | 25.072 | 6.39592 |
| PAV87B | PAV87 | Serpentine | Lavandula stoechas | 107 | 0.00356 | 3.56 | 25.503 | 7.16376 |
| PAV87C | PAV87 | Serpentine | Lavandula stoechas | 68 | 0.00205 | 2.05 | 20.303 | 9.90390 |
| PAV87D | PAV87 | Serpentine | Lavandula stoechas | 105 | 0.00319 | 3.19 | 30.386 | 9.52539 |
| PAV87E | PAV87 | Serpentine | Lavandula stoechas | 53 | 0.00486 | 4.86 | 27.966 | 5.75432 |
| PAV87F | PAV87 | Serpentine | Lavandula stoechas | 44 | | | | |
| PAV87G | PAV87 | Serpentine | Lavandula stoechas | 30 | 0.00172 | 1.72 | 17.624 | 10.24651 |
| PAV87H | PAV87 | Serpentine | Lavandula stoechas | 58 | 0.00204 | 2.04 | 16.238 | 7.95980 |
| PAV87I | PAV87 | Serpentine | Lavandula stoechas | 41 | 0.00449 | 4.49 | 32.189 | 7.16904 |
| PAV87J | PAV87 | Serpentine | Lavandula stoechas | 39 | 0.00308 | 3.08 | 18.989 | 6.16526 |
| PAV87K | PAV87 | Serpentine | Lavandula stoechas | 33 | 0.00593 | 5.93 | 42.581 | 7.18061 |
| PAV87L | PAV87 | Serpentine | Lavandula stoechas | 61 | 0.00434 | 4.34 | 30.832 | 7.10415 |
| PAV87M | PAV87 | Serpentine | Lavandula stoechas | 42 | 0.00274 | 2.74 | 26.422 | 9.64307 |
| PAV87N | PAV87 | Serpentine | Lavandula stoechas | 56 | 0.00335 | 3.35 | 23.542 | 7.02746 |
| PAV87Ñ | PAV87 | Serpentine | Lavandula stoechas | 36 | 0.00273 | 2.73 | 21.093 | 7.72637 |
| PAV43A | PAV43 | Serpentine | Halimium atriplicifolium | 159 | 0.397195 | 397.195 | 54.120 | 7.33920 |
| PAV43B | PAV43 | Serpentine | Halimium atriplicifolium | 91 | 0.437971 | 437.971 | 63.330 | 6.91570 |
| PAV43C | PAV43 | Serpentine | Halimium atriplicifolium | 89 | 0.458427 | 458.427 | 70.410 | 6.51080 |
| PAV43D | PAV43 | Serpentine | Halimium atriplicifolium | 104 | 0.326169 | 326.169 | 44.870 | 7.26920 |
| PAV43E | PAV43 | Serpentine | Halimium atriplicifolium | 89 | 0.379194 | 379.194 | 59.020 | 6.42480 |
| PAV43F | PAV43 | Serpentine | Halimium atriplicifolium | 82 | 0.335599 | 335.599 | 53.250 | 6.30230 |
| PAV43G | PAV43 | Serpentine | Halimium atriplicifolium | 60 | 0.286402 | 286.402 | 45.180 | 6.33910 |
| PAV43H | PAV43 | Serpentine | Halimium atriplicifolium | 147 | 0.25515 | 255.15 | 44.700 | 5.70810 |
| PAV43I | PAV43 | Serpentine | Halimium atriplicifolium | 69 | 0.35149 | 351.49 | 48.690 | 7.21890 |
| PAV43J | PAV43 | Serpentine | Halimium atriplicifolium | 107 | 0.446772 | 446.772 | 63.580 | 7.02690 |
| PAV43K | PAV43 | Serpentine | Halimium atriplicifolium | 74 | 0.328995 | 328.995 | 44.760 | 7.35020 |
| PAV43L | PAV43 | Serpentine | Halimium atriplicifolium | 80 | 0.329847 | 329.847 | 37.290 | 8.84550 |
| PAV43M | PAV43 | Serpentine | Halimium atriplicifolium | 65 | 0.275727 | 275.727 | 43.720 | 6.30670 |
| PAV43N | PAV43 | Serpentine | Halimium atriplicifolium | 100 | 0.249319 | 249.319 | 42.780 | 5.82790 |
| PAV43Ñ | PAV43 | Serpentine | Halimium atriplicifolium | 75 | 0.324037 | 324.037 | 54.310 | 5.96640 |
| PAV45A | PAV45 | Serpentine | Halimium atriplicifolium | 92 | 0.08278 | 82.78 | 530.868 | 6.41300 |
| PAV45B | PAV45 | Serpentine | Halimium atriplicifolium | 70 | | | | |
| PAV45C | PAV45 | Serpentine | Halimium atriplicifolium | 61 | | | | |
| PAV45D | PAV45 | Serpentine | Halimium atriplicifolium | 110 | 0.13781 | 137.81 | 720.698 | 5.22965 |
| PAV45E | PAV45 | Serpentine | Halimium atriplicifolium | 97 | 0.0737 | 73.7 | 422.508 | 5.73281 |
| PAV45F | PAV45 | Serpentine | Halimium atriplicifolium | 86 | 0.07791 | 77.91 | 683.238 | 8.76958 |
| PAV45G | PAV45 | Serpentine | Halimium atriplicifolium | 84 | 0.08124 | 81.24 | 628.684 | 7.73860 |
| PAV45H | PAV45 | Serpentine | Halimium atriplicifolium | 137 | | | | |
| PAV45I | PAV45 | Serpentine | Halimium atriplicifolium | 66 | 0.04249 | 42.49 | 365.656 | 8.60570 |
| PAV45J | PAV45 | Serpentine | Halimium atriplicifolium | 76 | | | | |
| PAV45K | PAV45 | Serpentine | Halimium atriplicifolium | 99 | 0.12391 | 123.91 | 731.837 | 5.90620 |
| PAV45L | PAV45 | Serpentine | Halimium atriplicifolium | 141 | 0.11868 | 118.68 | 1004.323 | 8.46245 |
| PAV45M | PAV45 | Serpentine | Halimium atriplicifolium | 74 | 0.0677 | 67.7 | 638.509 | 9.43145 |
| PAV45N | PAV45 | Serpentine | Halimium atriplicifolium | 65 | 0.08526 | 85.26 | 665.376 | 7.80408 |
| PAV45Ñ | PAV45 | Serpentine | Halimium atriplicifolium | 62 | 0.0713 | 71.3 | 611.893 | 8.58195 |
| PAV46A | PAV46 | Serpentine | Halimium atriplicifolium | 49 | 0.05523 | 55.23 | 424.447 | 7.68508 |
| PAV46B | PAV46 | Serpentine | Halimium atriplicifolium | 131 | 0.03613 | 36.13 | 310.399 | 8.59117 |
| PAV46C | PAV46 | Serpentine | Halimium atriplicifolium | 12 | 0.04262 | 42.62 | 361.757 | 8.48796 |
| PAV46D | PAV46 | Serpentine | Halimium atriplicifolium | 63 | 0.0551 | 55.1 | 502.894 | 9.12693 |
| PAV46E | PAV46 | Serpentine | Halimium atriplicifolium | 74 | 0.03566 | 35.66 | 261.469 | 7.33228 |
| PAV46F | PAV46 | Serpentine | Halimium atriplicifolium | 57 | 0.06443 | 64.43 | 384.983 | 5.97521 |
| PAV46G | PAV46 | Serpentine | Halimium atriplicifolium | 98 | 0.06948 | 69.48 | 339.931 | 4.89250 |
| PAV46H | PAV46 | Serpentine | Halimium atriplicifolium | 33 | 0.05964 | 59.64 | 344.542 | 5.77703 |
| PAV46I | PAV46 | Serpentine | Halimium atriplicifolium | 52 | 0.10007 | 100.07 | 576.271 | 5.75868 |
| PAV46J | PAV46 | Serpentine | Halimium atriplicifolium | 62 | 0.07686 | 76.86 | 470.468 | 6.12110 |
| PAV46K | PAV46 | Serpentine | Halimium atriplicifolium | 42 | 0.04366 | 43.66 | 399.792 | 9.15694 |
| PAV46L | PAV46 | Serpentine | Halimium atriplicifolium | 77 | | | | |
| PAV46M | PAV46 | Serpentine | Halimium atriplicifolium | 165 | 0.04361 | 43.61 | 281.988 | 6.46613 |
| PAV46N | PAV46 | Serpentine | Halimium atriplicifolium | 137 | 0.0307 | 30.7 | 319.219 | 10.39801 |
| PAV46Ñ | PAV46 | Serpentine | Halimium atriplicifolium | 39 | 0.03504 | 35.04 | 299.109 | 8.53622 |
| PAV47A | PAV47 | Nonserpentine | Halimium atriplicifolium | 202 | | | | |
| PAV47B | PAV47 | Nonserpentine | Halimium atriplicifolium | 102 | 0.08215 | 82.15 | 671.179 | 8.17016 |
| PAV47C | PAV47 | Nonserpentine | Halimium atriplicifolium | 47 | 0.07499 | 74.99 | 513.215 | 6.84378 |
| PAV47D | PAV47 | Nonserpentine | Halimium atriplicifolium | 32 | 0.07824 | 78.24 | 966.087 | 12.34774 |
| PAV47E | PAV47 | Nonserpentine | Halimium atriplicifolium | 97 | | | | |
| PAV47F | PAV47 | Nonserpentine | Halimium atriplicifolium | 77 | 0.08 | 80 | 718.687 | 8.98359 |
| PAV47G | PAV47 | Nonserpentine | Halimium atriplicifolium | 12 | 0.0578 | 57.8 | 709.983 | 12.28344 |
| PAV47H | PAV47 | Nonserpentine | Halimium atriplicifolium | 21 | 0.05883 | 58.83 | 711.656 | 12.09682 |
| PAV47I | PAV47 | Nonserpentine | Halimium atriplicifolium | 84 | 0.12293 | 122.93 | 956.923 | 7.78429 |
| PAV47J | PAV47 | Nonserpentine | Halimium atriplicifolium | 98 | 0.0943 | 94.3 | 770.691 | 8.17276 |
| PAV47K | PAV47 | Nonserpentine | Halimium atriplicifolium | 27 | 0.06443 | 64.43 | 536.383 | 8.32505 |
| PAV47L | PAV47 | Nonserpentine | Halimium atriplicifolium | 107 | 0.08494 | 84.94 | 712.712 | 8.39077 |
| PAV47M | PAV47 | Nonserpentine | Halimium atriplicifolium | 113 | 0.09819 | 98.19 | 927.399 | 9.44494 |
| PAV47N | PAV47 | Nonserpentine | Halimium atriplicifolium | 89 | 0.17383 | 173.83 | 1178.533 | 6.77980 |

| | | | | | | | | |
|---|---|---|---|---|---|---|---|---|
| PAV47Ñ | PAV47 | Nonserpentine | Halimium atriplicifolium | 68 | 0.07499 | 74.99 | 819.671 | 10.93040 |
| PAV50A | PAV50 | Serpentine | Halimium atriplicifolium | 109 | 0.10234 | 102.34 | 584.415 | 5.71052 |
| PAV50B | PAV50 | Serpentine | Halimium atriplicifolium | 92 | 0.0995 | 99.5 | 558.489 | 5.61295 |
| PAV50C | PAV50 | Serpentine | Halimium atriplicifolium | 87 | 0.09201 | 92.01 | 518.737 | 5.63783 |
| PAV50D | PAV50 | Serpentine | Halimium atriplicifolium | 76 | 0.06299 | 62.99 | 463.954 | 7.36552 |
| PAV50E | PAV50 | Serpentine | Halimium atriplicifolium | 66 | | | | |
| PAV50F | PAV50 | Serpentine | Halimium atriplicifolium | 117 | 0.1216 | 121.6 | 639.852 | 5.26194 |
| PAV50G | PAV50 | Serpentine | Halimium atriplicifolium | 35 | | | | |
| PAV50H | PAV50 | Serpentine | Halimium atriplicifolium | 82 | | | | |
| PAV50I | PAV50 | Serpentine | Halimium atriplicifolium | 91 | | | | |
| PAV50J | PAV50 | Serpentine | Halimium atriplicifolium | 75 | | | | |
| PAV50K | PAV50 | Serpentine | Halimium atriplicifolium | 61 | 0.07887 | 78.87 | 433.661 | 5.49843 |
| PAV50L | PAV50 | Serpentine | Halimium atriplicifolium | 110 | | | | |
| PAV50M | PAV50 | Serpentine | Halimium atriplicifolium | 23 | | | | |
| PAV50N | PAV50 | Serpentine | Halimium atriplicifolium | 107 | 0.06565 | 65.65 | 384.724 | 5.86023 |
| PAV50Ñ | PAV50 | Serpentine | Halimium atriplicifolium | 109 | 0.12627 | 126.27 | 696.366 | 5.51490 |
| PAV51A | PAV51 | Serpentine | Halimium atriplicifolium | 67 | | | | |
| PAV51B | PAV51 | Serpentine | Halimium atriplicifolium | 51 | | | | |
| PAV51C | PAV51 | Serpentine | Halimium atriplicifolium | 57 | | | | |
| PAV51D | PAV51 | Serpentine | Halimium atriplicifolium | 67 | | | | |
| PAV51E | PAV51 | Serpentine | Halimium atriplicifolium | 126 | | | | |
| PAV51F | PAV51 | Serpentine | Halimium atriplicifolium | 148 | | | | |
| PAV51G | PAV51 | Serpentine | Halimium atriplicifolium | 180 | | | | |
| PAV51H | PAV51 | Serpentine | Halimium atriplicifolium | 141 | 0.05078 | 50.78 | 270.756 | 5.33194 |
| PAV51I | PAV51 | Serpentine | Halimium atriplicifolium | 144 | 0.07951 | 79.51 | 421.804 | 5.30504 |
| PAV51J | PAV51 | Serpentine | Halimium atriplicifolium | 76 | 0.10347 | 103.47 | 598.506 | 5.78434 |
| PAV51K | PAV51 | Serpentine | Halimium atriplicifolium | 114 | 0.12952 | 129.52 | 720.971 | 5.56648 |
| PAV51L | PAV51 | Serpentine | Halimium atriplicifolium | 78 | 0.14612 | 146.12 | 756.679 | 5.17848 |
| PAV51M | PAV51 | Serpentine | Halimium atriplicifolium | 104 | 0.07063 | 70.63 | 444.836 | 6.29812 |
| PAV51N | PAV51 | Serpentine | Halimium atriplicifolium | 128 | 0.05393 | 53.93 | 343.263 | 6.36497 |
| PAV51Ñ | PAV51 | Serpentine | Halimium atriplicifolium | 109 | | | | |
| PAV52A | PAV52 | Serpentine | Halimium atriplicifolium | 71 | | | | |
| PAV52B | PAV52 | Serpentine | Halimium atriplicifolium | 88 | 0.06478 | 64.78 | 424.548 | 6.55369 |
| PAV52C | PAV52 | Serpentine | Halimium atriplicifolium | 64 | 0.10106 | 101.06 | 522.874 | 5.17390 |
| PAV52D | PAV52 | Serpentine | Halimium atriplicifolium | 61 | | | | |
| PAV52E | PAV52 | Serpentine | Halimium atriplicifolium | 62 | | | | |
| PAV52F | PAV52 | Serpentine | Halimium atriplicifolium | 87 | | | | |
| PAV52G | PAV52 | Serpentine | Halimium atriplicifolium | 89 | 0.0844 | 84.4 | 512.001 | 6.06636 |
| PAV52H | PAV52 | Serpentine | Halimium atriplicifolium | 99 | 0.12393 | 123.93 | 780.444 | 6.29746 |
| PAV52I | PAV52 | Serpentine | Halimium atriplicifolium | 108 | 0.1575 | 157.5 | 830.451 | 5.27270 |
| PAV52J | PAV52 | Serpentine | Halimium atriplicifolium | 63 | 0.12466 | 124.66 | 783.123 | 6.28207 |
| PAV52K | PAV52 | Serpentine | Halimium atriplicifolium | 34 | 0.0925 | 92.5 | 606.500 | 6.55676 |
| PAV52L | PAV52 | Serpentine | Halimium atriplicifolium | 101 | 0.13407 | 134.07 | 787.769 | 5.87580 |
| PAV52M | PAV52 | Serpentine | Halimium atriplicifolium | 43 | 0.05393 | 53.93 | 360.837 | 6.69084 |
| PAV52N | PAV52 | Serpentine | Halimium atriplicifolium | 23 | 0.05715 | 57.15 | 352.822 | 6.17361 |
| PAV52Ñ | PAV52 | Serpentine | Halimium atriplicifolium | 86 | 0.09685 | 96.85 | 622.084 | 6.42317 |
| PAV54A | PAV54 | Nonserpentine | Halimium atriplicifolium | 136 | 0.05355 | 53.55 | 342.481 | 6.39554 |
| PAV54B | PAV54 | Nonserpentine | Halimium atriplicifolium | 110 | 0.02402 | 24.02 | 190.182 | 7.91765 |
| PAV54C | PAV54 | Nonserpentine | Halimium atriplicifolium | 104 | 0.0473 | 47.3 | 249.799 | 5.28116 |
| PAV54D | PAV54 | Nonserpentine | Halimium atriplicifolium | 97 | 0.0354 | 35.4 | 249.871 | 7.05850 |
| PAV54E | PAV54 | Nonserpentine | Halimium atriplicifolium | 138 | 0.06508 | 65.08 | 398.750 | 6.12707 |
| PAV54F | PAV54 | Nonserpentine | Halimium atriplicifolium | 173 | 0.06403 | 64.03 | 393.888 | 6.15162 |
| PAV54G | PAV54 | Nonserpentine | Halimium atriplicifolium | 74 | 0.03264 | 32.64 | 285.550 | 8.74847 |
| PAV54H | PAV54 | Nonserpentine | Halimium atriplicifolium | 117 | | | | |
| PAV54I | PAV54 | Nonserpentine | Halimium atriplicifolium | 180 | 0.04337 | 43.37 | 296.122 | 6.82781 |
| PAV54J | PAV54 | Nonserpentine | Halimium atriplicifolium | 101 | | | | |
| PAV54K | PAV54 | Nonserpentine | Halimium atriplicifolium | 103 | | | | |
| PAV54L | PAV54 | Nonserpentine | Halimium atriplicifolium | 105 | 0.05936 | 59.36 | 405.652 | 6.83376 |
| PAV54M | PAV54 | Nonserpentine | Halimium atriplicifolium | 117 | 0.01892 | 18.92 | 145.526 | 7.69165 |
| PAV54N | PAV54 | Nonserpentine | Halimium atriplicifolium | 124 | 0.0558 | 55.8 | 276.558 | 4.95624 |
| PAV54Ñ | PAV54 | Nonserpentine | Halimium atriplicifolium | 106 | 0.02816 | 28.16 | 243.680 | 8.65341 |
| PAV55A | PAV55 | Nonserpentine | Halimium atriplicifolium | 66 | | | | |
| PAV55B | PAV55 | Nonserpentine | Halimium atriplicifolium | 81 | | | | |
| PAV55C | PAV55 | Nonserpentine | Halimium atriplicifolium | 170 | | | | |
| PAV55D | PAV55 | Nonserpentine | Halimium atriplicifolium | 91 | | | | |
| PAV55E | PAV55 | Nonserpentine | Halimium atriplicifolium | 90 | | | | |
| PAV55F | PAV55 | Nonserpentine | Halimium atriplicifolium | 72 | 0.03809 | 38.09 | 317.071 | 8.32426 |
| PAV55G | PAV55 | Nonserpentine | Halimium atriplicifolium | 61 | | | | |
| PAV55H | PAV55 | Nonserpentine | Halimium atriplicifolium | 31 | | | | |
| PAV55I | PAV55 | Nonserpentine | Halimium atriplicifolium | 35 | | | | |
| PAV55J | PAV55 | Nonserpentine | Halimium atriplicifolium | 102 | | | | |
| PAV55K | PAV55 | Nonserpentine | Halimium atriplicifolium | 45 | | | | |
| PAV55L | PAV55 | Nonserpentine | Halimium atriplicifolium | 74 | 0.04929 | 49.29 | 443.371 | 8.99515 |
| PAV55M | PAV55 | Nonserpentine | Halimium atriplicifolium | 57 | | | | |
| PAV55N | PAV55 | Nonserpentine | Halimium atriplicifolium | 96 | | | | |
| PAV55Ñ | PAV55 | Nonserpentine | Halimium atriplicifolium | 105 | | | | |
| PAV56A | PAV56 | Nonserpentine | Halimium atriplicifolium | 76 | 0.43918 | 439.18 | 53.330 | 8.23510 |
| PAV56B | PAV56 | Nonserpentine | Halimium atriplicifolium | 68 | 0.50698 | 506.98 | 83.250 | 6.08980 |
| PAV56C | PAV56 | Nonserpentine | Halimium atriplicifolium | 103 | 0.238644 | 238.644 | 49.210 | 4.84950 |
| PAV56D | PAV56 | Nonserpentine | Halimium atriplicifolium | 94 | 0.402533 | 402.533 | 80.460 | 5.00290 |
| PAV56E | PAV56 | Nonserpentine | Halimium atriplicifolium | 175 | | | | |
| PAV56F | PAV56 | Nonserpentine | Halimium atriplicifolium | 68 | 0.188688 | 188.688 | 27.430 | 6.87890 |
| PAV56G | PAV56 | Nonserpentine | Halimium atriplicifolium | 25 | 0.235932 | 235.932 | 43.610 | 5.41000 |
| PAV56H | PAV56 | Nonserpentine | Halimium atriplicifolium | 82 | 0.294974 | 294.974 | 49.760 | 5.92790 |
| PAV56I | PAV56 | Nonserpentine | Halimium atriplicifolium | 129 | 0.416456 | 416.456 | 68.470 | 6.08230 |
| PAV56J | PAV56 | Nonserpentine | Halimium atriplicifolium | 53 | 0.226208 | 226.208 | 30.760 | 7.35400 |
| PAV56K | PAV56 | Nonserpentine | Halimium atriplicifolium | 37 | 0.26878 | 268.78 | 37.400 | 7.18660 |
| PAV56L | PAV56 | Nonserpentine | Halimium atriplicifolium | 46 | 0.232905 | 232.905 | 29.470 | 7.90310 |
| PAV56M | PAV56 | Nonserpentine | Halimium atriplicifolium | 77 | 0.243409 | 243.409 | 31.600 | 7.70280 |
| PAV56N | PAV56 | Nonserpentine | Halimium atriplicifolium | 57 | 0.184224 | 184.224 | 20.100 | 9.16540 |

| | | | | | | | | |
|---|---|---|---|---|---|---|---|---|
| PAV56Ñ | PAV56 | Nonserpentine | Halimium atriplicifolium | 71 | 0.283576 | 283.576 | 58.340 | 4.86070 |
| PAV57A | PAV57 | Serpentine | Halimium atriplicifolium | 52 | | | | |
| PAV57B | PAV57 | Serpentine | Halimium atriplicifolium | 67 | | | | |
| PAV57C | PAV57 | Serpentine | Halimium atriplicifolium | 70 | | | | |
| PAV57D | PAV57 | Serpentine | Halimium atriplicifolium | 64 | 0.542625 | 542.625 | 50.270 | 10.79420 |
| PAV57E | PAV57 | Serpentine | Halimium atriplicifolium | 32 | 0.269882 | 269.882 | 28.150 | 9.58730 |
| PAV57F | PAV57 | Serpentine | Halimium atriplicifolium | 27 | 0.501163 | 501.163 | 77.780 | 6.44330 |
| PAV57G | PAV57 | Serpentine | Halimium atriplicifolium | 53 | 0.370265 | 370.265 | 54.950 | 6.73820 |
| PAV57H | PAV57 | Serpentine | Halimium atriplicifolium | 58 | 0.379516 | 379.516 | 43.710 | 8.68260 |
| PAV57I | PAV57 | Serpentine | Halimium atriplicifolium | 45 | 0.47376 | 473.76 | 38.350 | 12.35360 |
| PAV57J | PAV57 | Serpentine | Halimium atriplicifolium | 27 | 0.373499 | 373.499 | 37.970 | 9.83670 |
| PAV57K | PAV57 | Serpentine | Halimium atriplicifolium | 40 | 0.55544 | 555.44 | 67.120 | 8.27530 |
| PAV57L | PAV57 | Serpentine | Halimium atriplicifolium | 65 | 0.420535 | 420.535 | 50.350 | 8.35220 |
| PAV57M | PAV57 | Serpentine | Halimium atriplicifolium | 77 | 0.349201 | 349.201 | 40.800 | 8.55880 |
| PAV57N | PAV57 | Serpentine | Halimium atriplicifolium | 29 | 0.504898 | 504.898 | 67.220 | 7.51110 |
| PAV57Ñ | PAV57 | Serpentine | Halimium atriplicifolium | 37 | | | | |
| PAV58A | PAV58 | Serpentine | Halimium atriplicifolium | 28 | 0.04411 | 44.11 | 364.340 | 8.25981 |
| PAV58B | PAV58 | Serpentine | Halimium atriplicifolium | 59 | 0.06953 | 69.53 | 526.620 | 7.57400 |
| PAV58C | PAV58 | Serpentine | Halimium atriplicifolium | 33 | 0.04076 | 40.76 | 322.120 | 7.90285 |
| PAV58D | PAV58 | Serpentine | Halimium atriplicifolium | 31 | 0.06071 | 60.71 | 371.800 | 6.12420 |
| PAV58E | PAV58 | Serpentine | Halimium atriplicifolium | 25 | 0.04755 | 47.55 | 297.590 | 6.25846 |
| PAV58F | PAV58 | Serpentine | Halimium atriplicifolium | 44 | 0.06922 | 69.22 | 409.620 | 5.91765 |
| PAV58G | PAV58 | Serpentine | Halimium atriplicifolium | 34 | 0.04622 | 46.22 | 327.050 | 7.07594 |
| PAV58H | PAV58 | Serpentine | Halimium atriplicifolium | 39 | 0.05746 | 57.46 | 336.890 | 5.86304 |
| PAV58I | PAV58 | Serpentine | Halimium atriplicifolium | 27 | 0.04413 | 44.13 | 329.250 | 7.46091 |
| PAV58J | PAV58 | Serpentine | Halimium atriplicifolium | 61 | 0.03254 | 32.54 | 199.070 | 6.11770 |
| PAV58K | PAV58 | Serpentine | Halimium atriplicifolium | 40 | 0.04966 | 49.66 | 359.320 | 7.23560 |
| PAV58L | PAV58 | Serpentine | Halimium atriplicifolium | 34 | | | | |
| PAV58M | PAV58 | Serpentine | Halimium atriplicifolium | 42 | | | | |
| PAV58N | PAV58 | Serpentine | Halimium atriplicifolium | 71 | | | | |
| PAV58Ñ | PAV58 | Serpentine | Halimium atriplicifolium | 51 | | | | |
| PAV60A | PAV60 | Serpentine | Halimium atriplicifolium | 31 | | | | |
| PAV60B | PAV60 | Serpentine | Halimium atriplicifolium | 87 | 0.05839 | 58.39 | 602.550 | 10.31940 |
| PAV60C | PAV60 | Serpentine | Halimium atriplicifolium | 118 | 0.09756 | 97.56 | 587.690 | 6.02388 |
| PAV60D | PAV60 | Serpentine | Halimium atriplicifolium | 39 | 0.05023 | 50.23 | 617.624 | 12.29592 |
| PAV60E | PAV60 | Serpentine | Halimium atriplicifolium | 38 | | | | |
| PAV60F | PAV60 | Serpentine | Halimium atriplicifolium | 35 | 0.09832 | 98.32 | 757.239 | 7.70178 |
| PAV60G | PAV60 | Serpentine | Halimium atriplicifolium | 87 | 0.04677 | 46.77 | 345.605 | 7.38946 |
| PAV60H | PAV60 | Serpentine | Halimium atriplicifolium | 96 | | | | |
| PAV60I | PAV60 | Serpentine | Halimium atriplicifolium | 79 | 0.06932 | 69.32 | 387.453 | 5.58934 |
| PAV60J | PAV60 | Serpentine | Halimium atriplicifolium | 72 | | | | |
| PAV60K | PAV60 | Serpentine | Halimium atriplicifolium | 68 | | | | |
| PAV60L | PAV60 | Serpentine | Halimium atriplicifolium | 97 | | | | |
| PAV60M | PAV60 | Serpentine | Halimium atriplicifolium | 73 | 0.05017 | 50.17 | 399.354 | 7.96002 |
| PAV60N | PAV60 | Serpentine | Halimium atriplicifolium | 26 | 0.05888 | 58.88 | 456.679 | 7.75610 |
| PAV60Ñ | PAV60 | Serpentine | Halimium atriplicifolium | 33 | | | | |
| PAV93A | PAV93 | Nonserpentine | Halimium atriplicifolium | 73 | 0.06491 | 64.91 | 420.691 | 6.48114 |
| PAV93B | PAV93 | Nonserpentine | Halimium atriplicifolium | 81 | 0.05775 | 57.75 | 340.391 | 5.89422 |
| PAV93C | PAV93 | Nonserpentine | Halimium atriplicifolium | 35 | | | | |
| PAV93D | PAV93 | Nonserpentine | Halimium atriplicifolium | 28 | | | | |
| PAV93E | PAV93 | Nonserpentine | Halimium atriplicifolium | 52 | 0.10167 | 101.67 | 640.060 | 6.29547 |
| PAV93F | PAV93 | Nonserpentine | Halimium atriplicifolium | 88 | 0.07586 | 75.86 | 472.522 | 6.22887 |
| PAV93G | PAV93 | Nonserpentine | Halimium atriplicifolium | 96 | 0.05731 | 57.31 | 381.586 | 6.65828 |
| PAV93H | PAV93 | Nonserpentine | Halimium atriplicifolium | 21 | 0.03474 | 34.74 | 232.570 | 6.69459 |
| PAV93I | PAV93 | Nonserpentine | Halimium atriplicifolium | 36 | 0.02972 | 29.72 | 248.090 | 8.34758 |
| PAV93J | PAV93 | Nonserpentine | Halimium atriplicifolium | 83 | 0.04014 | 40.14 | 288.337 | 7.18328 |
| PAV93K | PAV93 | Nonserpentine | Halimium atriplicifolium | 26 | | | | |
| PAV93L | PAV93 | Nonserpentine | Halimium atriplicifolium | 53 | 0.05574 | 55.74 | 475.259 | 8.52635 |
| PAV93M | PAV93 | Nonserpentine | Halimium atriplicifolium | 33 | 0.03819 | 38.19 | 282.440 | 7.39565 |
| PAV93N | PAV93 | Nonserpentine | Halimium atriplicifolium | 68 | 0.12402 | 124.02 | 730.192 | 5.88770 |
| PAV93Ñ | PAV93 | Nonserpentine | Halimium atriplicifolium | 62 | | | | |
| PAV95A | PAV95 | Nonserpentine | Halimium atriplicifolium | 43 | 0.04052 | 40.52 | 347.673 | 8.58028 |
| PAV95B | PAV95 | Nonserpentine | Halimium atriplicifolium | 84 | 0.0666 | 66.6 | 768.946 | 11.54574 |
| PAV95C | PAV95 | Nonserpentine | Halimium atriplicifolium | 51 | 0.07313 | 73.13 | 832.749 | 11.38724 |
| PAV95D | PAV95 | Nonserpentine | Halimium atriplicifolium | 65 | 0.05471 | 54.71 | 494.427 | 9.03723 |
| PAV95E | PAV95 | Nonserpentine | Halimium atriplicifolium | 51 | 0.03708 | 37.08 | 277.973 | 7.49657 |
| PAV95F | PAV95 | Nonserpentine | Halimium atriplicifolium | 78 | 0.05145 | 51.45 | 439.866 | 8.54939 |
| PAV95G | PAV95 | Nonserpentine | Halimium atriplicifolium | 149 | 0.15314 | 153.14 | 1249.490 | 8.15914 |
| PAV95H | PAV95 | Nonserpentine | Halimium atriplicifolium | 109 | 0.08746 | 87.46 | 939.306 | 10.73984 |
| PAV95I | PAV95 | Nonserpentine | Halimium atriplicifolium | 146 | 0.07076 | 70.76 | 578.950 | 8.18188 |
| PAV95J | PAV95 | Nonserpentine | Halimium atriplicifolium | 117 | | | | |
| PAV95K | PAV95 | Nonserpentine | Halimium atriplicifolium | 200 | | | | |
| PAV95L | PAV95 | Nonserpentine | Halimium atriplicifolium | 71 | | | | |
| PAV95M | PAV95 | Nonserpentine | Halimium atriplicifolium | 79 | 0.09829 | 98.29 | 995.037 | 10.12348 |
| PAV95N | PAV95 | Nonserpentine | Halimium atriplicifolium | 98 | 0.06719 | 67.19 | 767.955 | 11.42960 |
| PAV95Ñ | PAV95 | Nonserpentine | Halimium atriplicifolium | 74 | 0.11731 | 117.31 | 925.564 | 7.88990 |
| PAV97A | PAV97 | Nonserpentine | Halimium atriplicifolium | 121 | 0.0985 | 98.5 | 553.691 | 5.62123 |
| PAV97B | PAV97 | Nonserpentine | Halimium atriplicifolium | 54 | 0.08201 | 82.01 | 549.253 | 6.69739 |
| PAV97C | PAV97 | Nonserpentine | Halimium atriplicifolium | 45 | 0.04563 | 45.63 | 443.737 | 9.72468 |
| PAV97D | PAV97 | Nonserpentine | Halimium atriplicifolium | 116 | 0.07446 | 74.46 | 561.060 | 7.53505 |
| PAV97E | PAV97 | Nonserpentine | Halimium atriplicifolium | 49 | 0.04874 | 48.74 | 430.027 | 8.82288 |
| PAV97F | PAV97 | Nonserpentine | Halimium atriplicifolium | 48 | | | | |
| PAV97G | PAV97 | Nonserpentine | Halimium atriplicifolium | 77 | 0.09758 | 97.58 | 551.063 | 5.64729 |
| PAV97H | PAV97 | Nonserpentine | Halimium atriplicifolium | 45 | 0.06058 | 60.58 | 400.410 | 6.60961 |
| PAV97I | PAV97 | Nonserpentine | Halimium atriplicifolium | 66 | 0.05939 | 59.39 | 424.081 | 7.14061 |
| PAV97J | PAV97 | Nonserpentine | Halimium atriplicifolium | 64 | | | | |
| PAV97K | PAV97 | Nonserpentine | Halimium atriplicifolium | 67 | | | | |
| PAV97L | PAV97 | Nonserpentine | Halimium atriplicifolium | 26 | | | | |
| PAV97M | PAV97 | Nonserpentine | Halimium atriplicifolium | 85 | 0.02433 | 24.33 | 163.279 | 6.71102 |
| PAV97N | PAV97 | Nonserpentine | Halimium atriplicifolium | 86 | 0.08073 | 80.73 | 470.238 | 5.82482 |

| | | | | | | | | |
|---|---|---|---|---|---|---|---|---|
| PAV97Ñ | PAV97 | Nonserpentine | Halimium atriplicifolium | 51 | | | | |
| PAV98A | PAV98 | Nonserpentine | Halimium atriplicifolium | 115 | | | | |
| PAV98B | PAV98 | Nonserpentine | Halimium atriplicifolium | 104 | | | | |
| PAV98C | PAV98 | Nonserpentine | Halimium atriplicifolium | 72 | 0.10683 | 106.83 | 677.573 | 6.34253 |
| PAV98D | PAV98 | Nonserpentine | Halimium atriplicifolium | 127 | | | | |
| PAV98E | PAV98 | Nonserpentine | Halimium atriplicifolium | 52 | | | | |
| PAV98F | PAV98 | Nonserpentine | Halimium atriplicifolium | 132 | | | | |
| PAV98G | PAV98 | Nonserpentine | Halimium atriplicifolium | 84 | | | | |
| PAV98H | PAV98 | Nonserpentine | Halimium atriplicifolium | 107 | | | | |
| PAV98I | PAV98 | Nonserpentine | Halimium atriplicifolium | 119 | | | | |
| PAV98J | PAV98 | Nonserpentine | Halimium atriplicifolium | 91 | | | | |
| PAV98K | PAV98 | Nonserpentine | Halimium atriplicifolium | 84 | | | | |
| PAV98L | PAV98 | Nonserpentine | Halimium atriplicifolium | 76 | | | | |
| PAV98M | PAV98 | Nonserpentine | Halimium atriplicifolium | 73 | | | | |
| PAV98N | PAV98 | Nonserpentine | Halimium atriplicifolium | 116 | | | | |
| PAV98Ñ | PAV98 | Nonserpentine | Halimium atriplicifolium | 169 | | | | |
| PAV99A | PAV99 | Serpentine | Halimium atriplicifolium | 51 | | | | |
| PAV99B | PAV99 | Serpentine | Halimium atriplicifolium | 66 | 0.04733 | 47.33 | 335.557 | 7.08973 |
| PAV99C | PAV99 | Serpentine | Halimium atriplicifolium | 118 | 0.07396 | 73.96 | 622.034 | 8.41041 |
| PAV99D | PAV99 | Serpentine | Halimium atriplicifolium | 97 | | | | |
| PAV99E | PAV99 | Serpentine | Halimium atriplicifolium | 116 | | | | |
| PAV99F | PAV99 | Serpentine | Halimium atriplicifolium | 48 | 0.04719 | 47.19 | 344.958 | 7.30998 |
| PAV99G | PAV99 | Serpentine | Halimium atriplicifolium | 75 | 0.07315 | 73.15 | 469.527 | 6.41869 |
| PAV99H | PAV99 | Serpentine | Halimium atriplicifolium | 102 | 0.05187 | 51.87 | 368.860 | 7.11124 |
| PAV99I | PAV99 | Serpentine | Halimium atriplicifolium | 82 | 0.08268 | 82.68 | 582.728 | 7.04799 |
| PAV99J | PAV99 | Serpentine | Halimium atriplicifolium | 83 | 0.06955 | 69.55 | 443.587 | 6.37796 |
| PAV99K | PAV99 | Serpentine | Halimium atriplicifolium | 113 | 0.04783 | 47.83 | 325.079 | 6.79655 |
| PAV99L | PAV99 | Serpentine | Halimium atriplicifolium | 91 | 0.05172 | 51.72 | 321.467 | 6.21553 |
| PAV99M | PAV99 | Serpentine | Halimium atriplicifolium | 62 | 0.11564 | 115.64 | 737.109 | 6.37417 |
| PAV99N | PAV99 | Serpentine | Halimium atriplicifolium | 77 | | | | |
| PAV99Ñ | PAV99 | Serpentine | Halimium atriplicifolium | 74 | 0.08991 | 89.91 | 554.259 | 6.16460 |
| PAV106A | PAV106 | Nonserpentine | Halimium atriplicifolium | 38 | | | | |
| PAV106B | PAV106 | Nonserpentine | Halimium atriplicifolium | 55 | 0.06874 | 68.74 | 483.898 | 7.03954 |
| PAV106C | PAV106 | Nonserpentine | Halimium atriplicifolium | 63 | 0.09913 | 99.13 | 647.687 | 6.53371 |
| PAV106D | PAV106 | Nonserpentine | Halimium atriplicifolium | 85 | 0.05142 | 51.42 | 528.110 | 10.27052 |
| PAV106E | PAV106 | Nonserpentine | Halimium atriplicifolium | 79 | 0.08018 | 80.18 | 489.299 | 6.10251 |
| PAV106F | PAV106 | Nonserpentine | Halimium atriplicifolium | 68 | | | | |
| PAV106G | PAV106 | Nonserpentine | Halimium atriplicifolium | 32 | 0.12768 | 127.68 | 769.920 | 6.03008 |
| PAV106H | PAV106 | Nonserpentine | Halimium atriplicifolium | 59 | 0.1044 | 104.4 | 644.089 | 6.16943 |
| PAV106I | PAV106 | Nonserpentine | Halimium atriplicifolium | 72 | | | | |
| PAV106J | PAV106 | Nonserpentine | Halimium atriplicifolium | 56 | 0.05776 | 57.76 | 538.272 | 9.31911 |
| PAV106K | PAV106 | Nonserpentine | Halimium atriplicifolium | 88 | 0.1073 | 107.3 | 620.361 | 5.78156 |
| PAV106L | PAV106 | Nonserpentine | Halimium atriplicifolium | 83 | 0.04698 | 46.98 | 298.765 | 6.35941 |
| PAV106M | PAV106 | Nonserpentine | Halimium atriplicifolium | 117 | 0.0844 | 84.4 | 432.196 | 5.12081 |
| PAV106N | PAV106 | Nonserpentine | Halimium atriplicifolium | 24 | 0.08213 | 82.13 | 468.034 | 5.69870 |
| PAV106Ñ | PAV106 | Nonserpentine | Halimium atriplicifolium | 115 | 0.08154 | 81.54 | 403.505 | 4.94855 |
| PAV67A | PAV67 | Serpentine | Phlomis purpurea | 61 | 0.08193 | 81.93 | 869.039 | 10.60709 |
| PAV67B | PAV67 | Serpentine | Phlomis purpurea | 60 | | | | |
| PAV67C | PAV67 | Serpentine | Phlomis purpurea | 81 | | | | |
| PAV67D | PAV67 | Serpentine | Phlomis purpurea | 47 | | | | |
| PAV67E | PAV67 | Serpentine | Phlomis purpurea | 68 | 0.12092 | 120.92 | 1352.528 | 11.18531 |
| PAV67F | PAV67 | Serpentine | Phlomis purpurea | 45 | 0.11068 | 110.68 | 1342.165 | 12.12654 |
| PAV67G | PAV67 | Serpentine | Phlomis purpurea | 125 | 0.09519 | 95.19 | 1260.780 | 13.24488 |
| PAV67H | PAV67 | Serpentine | Phlomis purpurea | 42 | 0.06003 | 60.03 | 1024.691 | 17.06965 |
| PAV67I | PAV67 | Serpentine | Phlomis purpurea | 74 | 0.07625 | 76.25 | 1306.435 | 17.13357 |
| PAV67J | PAV67 | Serpentine | Phlomis purpurea | 55 | 0.07321 | 73.21 | 1273.370 | 17.39339 |
| PAV67K | PAV67 | Serpentine | Phlomis purpurea | 46 | | | | |
| PAV67L | PAV67 | Serpentine | Phlomis purpurea | 64 | 0.12105 | 121.05 | 1768.960 | 14.61347 |
| PAV67M | PAV67 | Serpentine | Phlomis purpurea | 72 | 0.12633 | 126.33 | 1738.193 | 13.75915 |
| PAV67N | PAV67 | Serpentine | Phlomis purpurea | 65 | 0.08901 | 89.01 | 1038.042 | 11.66208 |
| PAV67Ñ | PAV67 | Serpentine | Phlomis purpurea | 82 | | | | |
| PAV68A | PAV68 | Serpentine | Phlomis purpurea | 120 | 0.068590 | 68.59 | 439.094 | 6.40172 |
| PAV68B | PAV68 | Serpentine | Phlomis purpurea | 68 | 0.086940 | 86.94 | 680.922 | 7.83209 |
| PAV68C | PAV68 | Serpentine | Phlomis purpurea | 69 | 0.052500 | 52.5 | 352.599 | 6.71617 |
| PAV68D | PAV68 | Serpentine | Phlomis purpurea | 48 | 0.063360 | 63.36 | 396.723 | 6.26141 |
| PAV68E | PAV68 | Serpentine | Phlomis purpurea | 72 | 0.057850 | 57.85 | 451.344 | 7.80197 |
| PAV68F | PAV68 | Serpentine | Phlomis purpurea | 87 | 0.091520 | 91.52 | 553.272 | 6.04537 |
| PAV68G | PAV68 | Serpentine | Phlomis purpurea | 47 | 0.039660 | 39.66 | 299.689 | 7.55645 |
| PAV68H | PAV68 | Serpentine | Phlomis purpurea | 73 | 0.112750 | 112.75 | 596.924 | 5.29423 |
| PAV68I | PAV68 | Serpentine | Phlomis purpurea | 47 | 0.051960 | 51.96 | 298.322 | 5.74138 |
| PAV68J | PAV68 | Serpentine | Phlomis purpurea | 40 | 0.065110 | 65.11 | 356.191 | 5.47060 |
| PAV68K | PAV68 | Serpentine | Phlomis purpurea | 59 | 0.134650 | 134.65 | 745.437 | 5.53611 |
| PAV68L | PAV68 | Serpentine | Phlomis purpurea | 82 | 0.083890 | 83.89 | 465.839 | 5.55297 |
| PAV68M | PAV68 | Serpentine | Phlomis purpurea | 44 | 0.070400 | 70.4 | 426.223 | 6.05430 |
| PAV68N | PAV68 | Serpentine | Phlomis purpurea | 45 | 0.143850 | 143.85 | 899.403 | 6.25237 |
| PAV68Ñ | PAV68 | Serpentine | Phlomis purpurea | 51 | 0.081220 | 81.22 | 534.297 | 6.57839 |
| PAV69A | PAV69 | Nonserpentine | Phlomis purpurea | 111 | 0.10849 | 108.49 | 991.744 | 9.14134 |
| PAV69B | PAV69 | Nonserpentine | Phlomis purpurea | 82 | 0.13126 | 131.26 | 1283.662 | 9.77954 |
| PAV69C | PAV69 | Nonserpentine | Phlomis purpurea | 78 | 0.10203 | 102.03 | 914.085 | 8.95898 |
| PAV69D | PAV69 | Nonserpentine | Phlomis purpurea | 74 | 0.11286 | 112.86 | 1235.889 | 10.95064 |
| PAV69E | PAV69 | Nonserpentine | Phlomis purpurea | 99 | 0.11688 | 116.88 | 1420.735 | 12.15550 |
| PAV69F | PAV69 | Nonserpentine | Phlomis purpurea | 84 | 0.09926 | 99.26 | 1031.375 | 10.39064 |
| PAV69G | PAV69 | Nonserpentine | Phlomis purpurea | 50 | 0.07971 | 79.71 | 898.773 | 11.27554 |
| PAV69H | PAV69 | Nonserpentine | Phlomis purpurea | 115 | 0.06433 | 64.33 | 1126.778 | 17.51559 |
| PAV69I | PAV69 | Nonserpentine | Phlomis purpurea | 80 | 0.06432 | 64.32 | 951.970 | 14.80053 |
| PAV69J | PAV69 | Nonserpentine | Phlomis purpurea | 96 | 0.048 | 48 | 835.195 | 17.39990 |
| PAV69K | PAV69 | Nonserpentine | Phlomis purpurea | 116 | 0.10138 | 101.38 | 1879.248 | 18.53667 |
| PAV69L | PAV69 | Nonserpentine | Phlomis purpurea | 149 | 0.1671 | 167.1 | 2018.997 | 12.08257 |
| PAV69M | PAV69 | Nonserpentine | Phlomis purpurea | 87 | 0.12786 | 127.86 | 1609.402 | 12.58722 |
| PAV69N | PAV69 | Nonserpentine | Phlomis purpurea | 118 | 0.07464 | 74.64 | 1582.335 | 21.19956 |

| | | | | | | | | |
|---|---|---|---|---|---|---|---|---|
| PAV69Ñ | PAV69 | Nonserpentine | Phlomis purpurea | 81 | 0.13972 | 139.72 | 2516.611 | 18.01182 |
| PAV70A | PAV70 | Nonserpentine | Phlomis purpurea | 109 | 0.12983 | 129.83 | 919.150 | 7.07964 |
| PAV70B | PAV70 | Nonserpentine | Phlomis purpurea | 113 | 0.11507 | 115.07 | 733.904 | 6.37789 |
| PAV70C | PAV70 | Nonserpentine | Phlomis purpurea | 83 | 0.10005 | 100.05 | 535.592 | 5.35324 |
| PAV70D | PAV70 | Nonserpentine | Phlomis purpurea | 106 | 0.08678 | 86.78 | 643.180 | 7.41162 |
| PAV70E | PAV70 | Nonserpentine | Phlomis purpurea | 90 | 0.9445 | 944.5 | 717.791 | 0.75997 |
| PAV70F | PAV70 | Nonserpentine | Phlomis purpurea | 69 | 0.08646 | 86.46 | 688.828 | 7.96701 |
| PAV70G | PAV70 | Nonserpentine | Phlomis purpurea | 105 | 0.09653 | 96.53 | 686.896 | 7.11588 |
| PAV70H | PAV70 | Nonserpentine | Phlomis purpurea | 63 | 0.1319 | 131.9 | 804.579 | 6.09992 |
| PAV70I | PAV70 | Nonserpentine | Phlomis purpurea | 60 | 0.09287 | 92.87 | 671.341 | 7.22883 |
| PAV70J | PAV70 | Nonserpentine | Phlomis purpurea | 87 | 0.10928 | 109.28 | 834.601 | 7.63727 |
| PAV70K | PAV70 | Nonserpentine | Phlomis purpurea | 112 | 0.10633 | 106.33 | 750.288 | 7.05622 |
| PAV70L | PAV70 | Nonserpentine | Phlomis purpurea | 67 | 0.08753 | 87.53 | 689.551 | 7.87788 |
| PAV70M | PAV70 | Nonserpentine | Phlomis purpurea | 91 | 0.0894 | 89.4 | 573.642 | 6.41658 |
| PAV70N | PAV70 | Nonserpentine | Phlomis purpurea | 109 | 0.0976 | 97.6 | 770.601 | 7.89550 |
| PAV70Ñ | PAV70 | Nonserpentine | Phlomis purpurea | 78 | 0.1025 | 102.5 | 711.266 | 6.93918 |
| PAV71A | PAV71 | Serpentine | Phlomis purpurea | 78 | 0.0586 | 58.6 | 634.143 | 10.82155 |
| PAV71B | PAV71 | Serpentine | Phlomis purpurea | 59 | 0.06171 | 61.71 | 589.361 | 9.55049 |
| PAV71C | PAV71 | Serpentine | Phlomis purpurea | 74 | 0.11084 | 110.84 | 1001.088 | 9.03183 |
| PAV71D | PAV71 | Serpentine | Phlomis purpurea | 55 | 0.07132 | 71.32 | 750.839 | 10.52775 |
| PAV71E | PAV71 | Serpentine | Phlomis purpurea | 57 | 0.2811 | 281.1 | 1444.547 | 5.13891 |
| PAV71F | PAV71 | Serpentine | Phlomis purpurea | 75 | 0.08375 | 83.75 | 726.985 | 8.68042 |
| PAV71G | PAV71 | Serpentine | Phlomis purpurea | 44 | 0.09167 | 91.67 | 634.251 | 6.91885 |
| PAV71H | PAV71 | Serpentine | Phlomis purpurea | 72 | 0.07334 | 73.34 | 599.428 | 8.17328 |
| PAV71I | PAV71 | Serpentine | Phlomis purpurea | 62 | 0.08726 | 87.26 | 847.265 | 9.70966 |
| PAV71J | PAV71 | Serpentine | Phlomis purpurea | 88 | 0.12171 | 121.71 | 1144.550 | 9.40391 |
| PAV71K | PAV71 | Serpentine | Phlomis purpurea | 62 | 0.11755 | 117.55 | 1103.159 | 9.38459 |
| PAV71L | PAV71 | Serpentine | Phlomis purpurea | 53 | 0.1415 | 141.5 | 1198.255 | 8.46823 |
| PAV71M | PAV71 | Serpentine | Phlomis purpurea | 56 | 0.07163 | 71.63 | 735.099 | 10.26245 |
| PAV71N | PAV71 | Serpentine | Phlomis purpurea | 76 | 0.10416 | 104.16 | 919.558 | 8.82832 |
| PAV71Ñ | PAV71 | Serpentine | Phlomis purpurea | 41 | 0.09192 | 91.92 | 1418.424 | 15.43107 |
| PAV72A | PAV72 | Serpentine | Phlomis purpurea | 27 | 0.16051 | 160.51 | 811.820 | 5.05775 |
| PAV72B | PAV72 | Serpentine | Phlomis purpurea | 33 | 0.255 | 255 | 1309.835 | 5.13661 |
| PAV72C | PAV72 | Serpentine | Phlomis purpurea | 45 | 0.13566 | 135.66 | 1024.635 | 7.55296 |
| PAV72D | PAV72 | Serpentine | Phlomis purpurea | 51 | 0.06051 | 60.51 | 624.005 | 10.31243 |
| PAV72E | PAV72 | Serpentine | Phlomis purpurea | 62 | 0.25708 | 257.08 | 1539.664 | 5.98905 |
| PAV72F | PAV72 | Serpentine | Phlomis purpurea | 70 | 0.2421 | 242.1 | 1487.877 | 6.14571 |
| PAV72G | PAV72 | Serpentine | Phlomis purpurea | 57 | 0.18935 | 189.35 | 1028.076 | 5.42950 |
| PAV72H | PAV72 | Serpentine | Phlomis purpurea | 60 | 0.08487 | 84.87 | 753.286 | 8.87576 |
| PAV72I | PAV72 | Serpentine | Phlomis purpurea | 81 | 0.04101 | 41.01 | 365.821 | 8.92029 |
| PAV72J | PAV72 | Serpentine | Phlomis purpurea | 54 | 0.07466 | 74.66 | 549.050 | 7.35400 |
| PAV72K | PAV72 | Serpentine | Phlomis purpurea | 97 | 0.07489 | 74.89 | 610.196 | 8.14790 |
| PAV72L | PAV72 | Serpentine | Phlomis purpurea | 50 | 0.1178 | 117.8 | 810.332 | 6.87888 |
| PAV72M | PAV72 | Serpentine | Phlomis purpurea | 37 | 0.16479 | 164.79 | 1077.373 | 6.53785 |
| PAV72N | PAV72 | Serpentine | Phlomis purpurea | 28 | 0.13225 | 132.25 | 783.043 | 5.92093 |
| PAV72Ñ | PAV72 | Serpentine | Phlomis purpurea | 54 | 0.08811 | 88.11 | 849.548 | 9.64190 |
| PAV73A | PAV73 | Nonserpentine | Phlomis purpurea | 56 | 0.11707 | 117.07 | 1053.226 | 8.99655 |
| PAV73B | PAV73 | Nonserpentine | Phlomis purpurea | 76 | 0.136 | 136 | 1157.322 | 8.50972 |
| PAV73C | PAV73 | Nonserpentine | Phlomis purpurea | 78 | 0.167 | 167 | 1237.878 | 7.41244 |
| PAV73D | PAV73 | Nonserpentine | Phlomis purpurea | 83 | 0.08632 | 86.32 | 886.796 | 10.27335 |
| PAV73E | PAV73 | Nonserpentine | Phlomis purpurea | 48 | 0.12028 | 120.28 | 1049.934 | 8.72908 |
| PAV73F | PAV73 | Nonserpentine | Phlomis purpurea | 43 | 0.07599 | 75.99 | 574.092 | 7.55484 |
| PAV73G | PAV73 | Nonserpentine | Phlomis purpurea | 41 | 0.13824 | 138.24 | 1055.966 | 7.63864 |
| PAV73H | PAV73 | Nonserpentine | Phlomis purpurea | 66 | 0.13169 | 131.69 | 953.630 | 7.24148 |
| PAV73I | PAV73 | Nonserpentine | Phlomis purpurea | 60 | 0.25796 | 257.96 | 2543.757 | 9.86105 |
| PAV73J | PAV73 | Nonserpentine | Phlomis purpurea | 91 | 0.08556 | 85.56 | 1154.174 | 13.48964 |
| PAV73K | PAV73 | Nonserpentine | Phlomis purpurea | 97 | 0.09282 | 92.82 | 1445.284 | 15.57083 |
| PAV73L | PAV73 | Nonserpentine | Phlomis purpurea | 77 | 0.16028 | 160.28 | 1930.327 | 12.04347 |
| PAV73M | PAV73 | Nonserpentine | Phlomis purpurea | 84 | 0.18061 | 180.61 | 1836.612 | 10.16894 |
| PAV73N | PAV73 | Nonserpentine | Phlomis purpurea | 62 | 0.21919 | 219.19 | 2042.909 | 9.32027 |
| PAV73Ñ | PAV73 | Nonserpentine | Phlomis purpurea | 47 | | | | |
| PAV74A | PAV74 | Nonserpentine | Phlomis purpurea | 27 | | | | |
| PAV74B | PAV74 | Nonserpentine | Phlomis purpurea | 80 | | | | |
| PAV74C | PAV74 | Nonserpentine | Phlomis purpurea | 78 | 0.1493 | 149.3 | 1457.920 | 9.76504 |
| PAV74D | PAV74 | Nonserpentine | Phlomis purpurea | 84 | 0.14239 | 142.39 | 1204.137 | 8.45661 |
| PAV74E | PAV74 | Nonserpentine | Phlomis purpurea | 161 | 0.12618 | 126.18 | 507.196 | 4.01962 |
| PAV74F | PAV74 | Nonserpentine | Phlomis purpurea | 73 | 0.1255 | 125.5 | 1020.784 | |
| PAV74G | PAV74 | Nonserpentine | Phlomis purpurea | 46 | | | | |
| PAV74H | PAV74 | Nonserpentine | Phlomis purpurea | 19 | | | | |
| PAV74I | PAV74 | Nonserpentine | Phlomis purpurea | 41 | 0.13869 | 138.69 | 744.922 | 5.37113 |
| PAV74J | PAV74 | Nonserpentine | Phlomis purpurea | 43 | 0.19643 | 196.43 | 1886.046 | 9.60162 |
| PAV74K | PAV74 | Nonserpentine | Phlomis purpurea | 82 | | | | |
| PAV74L | PAV74 | Nonserpentine | Phlomis purpurea | 143 | 0.11274 | 112.74 | 578.397 | 5.13036 |
| PAV74M | PAV74 | Nonserpentine | Phlomis purpurea | 109 | 0.16465 | 164.65 | 1746.869 | 10.60959 |
| PAV74N | PAV74 | Nonserpentine | Phlomis purpurea | 126 | 0.11419 | 114.19 | 569.599 | 4.98817 |
| PAV74Ñ | PAV74 | Nonserpentine | Phlomis purpurea | 106 | 0.14925 | 149.25 | 809.832 | 5.42601 |
| PAV75A | PAV75 | Serpentine | Phlomis purpurea | 62 | | | | |
| PAV75B | PAV75 | Serpentine | Phlomis purpurea | 61 | | | | |
| PAV75C | PAV75 | Serpentine | Phlomis purpurea | 29 | | | | |
| PAV75D | PAV75 | Serpentine | Phlomis purpurea | 28 | 0.09825 | 98.25 | 590.901 | 6.01426 |
| PAV75E | PAV75 | Serpentine | Phlomis purpurea | 75 | 0.06734 | 67.34 | 490.254 | 7.28028 |
| PAV75F | PAV75 | Serpentine | Phlomis purpurea | 33 | | | | |
| PAV75G | PAV75 | Serpentine | Phlomis purpurea | 39 | | | | |
| PAV75H | PAV75 | Serpentine | Phlomis purpurea | 52 | | | | |
| PAV75I | PAV75 | Serpentine | Phlomis purpurea | 93 | | | | |
| PAV75J | PAV75 | Serpentine | Phlomis purpurea | 142 | | | | |
| PAV75K | PAV75 | Serpentine | Phlomis purpurea | 38 | | | | |
| PAV75L | PAV75 | Serpentine | Phlomis purpurea | 57 | | | | |
| PAV75M | PAV75 | Serpentine | Phlomis purpurea | 43 | | | | |
| PAV75N | PAV75 | Serpentine | Phlomis purpurea | 59 | | | | |

| | | | | | | | | |
|---|---|---|---|---|---|---|---|---|
| PAV75Ñ | PAV75 | Serpentine | Phlomis purpurea | 148 | | | | |
| PAV76A | PAV76 | Nonserpentine | Phlomis purpurea | 102 | 0.07648 | 76.48 | 673.370 | 8.80452 |
| PAV76B | PAV76 | Nonserpentine | Phlomis purpurea | 46 | | | | |
| PAV76C | PAV76 | Nonserpentine | Phlomis purpurea | 47 | | | | |
| PAV76D | PAV76 | Nonserpentine | Phlomis purpurea | 67 | | | | |
| PAV76E | PAV76 | Nonserpentine | Phlomis purpurea | 33 | | | | |
| PAV76F | PAV76 | Nonserpentine | Phlomis purpurea | 99 | | | | |
| PAV76G | PAV76 | Nonserpentine | Phlomis purpurea | 117 | 0.09037 | 90.37 | 841.001 | 9.30620 |
| PAV76H | PAV76 | Nonserpentine | Phlomis purpurea | 32 | | | | |
| PAV76I | PAV76 | Nonserpentine | Phlomis purpurea | 71 | 0.06221 | 62.21 | 470.799 | 7.56790 |
| PAV76J | PAV76 | Nonserpentine | Phlomis purpurea | 64 | 0.09239 | 92.39 | 965.010 | 10.44496 |
| PAV76K | PAV76 | Nonserpentine | Phlomis purpurea | 69 | 0.07388 | 73.88 | 952.344 | 12.89042 |
| PAV76L | PAV76 | Nonserpentine | Phlomis purpurea | 71 | 0.07387 | 73.87 | 451.487 | 6.11191 |
| PAV76M | PAV76 | Nonserpentine | Phlomis purpurea | 66 | 0.10443 | 104.43 | 930.753 | 8.91270 |
| PAV76N | PAV76 | Nonserpentine | Phlomis purpurea | 42 | | | | |
| PAV76Ñ | PAV76 | Nonserpentine | Phlomis purpurea | 112 | 0.06448 | 64.48 | 470.490 | 7.29668 |
| PAV77A | PAV77 | Nonserpentine | Phlomis purpurea | 44 | 0.12908 | 129.08 | 875.172 | 6.78007 |
| PAV77B | PAV77 | Nonserpentine | Phlomis purpurea | 60 | 0.09439 | 94.39 | 592.416 | 6.27626 |
| PAV77C | PAV77 | Nonserpentine | Phlomis purpurea | 48 | 0.0623 | 62.3 | 414.148 | 6.64764 |
| PAV77D | PAV77 | Nonserpentine | Phlomis purpurea | 104 | | | | |
| PAV77E | PAV77 | Nonserpentine | Phlomis purpurea | 45 | 0.05621 | 56.21 | 423.312 | 7.53090 |
| PAV77F | PAV77 | Nonserpentine | Phlomis purpurea | 98 | 0.09856 | 98.56 | 559.473 | 5.67647 |
| PAV77G | PAV77 | Nonserpentine | Phlomis purpurea | 64 | | | | |
| PAV77H | PAV77 | Nonserpentine | Phlomis purpurea | 49 | 0.13292 | 132.92 | 666.439 | 5.01384 |
| PAV77I | PAV77 | Nonserpentine | Phlomis purpurea | 81 | 0.11863 | 118.63 | 653.677 | 5.51022 |
| PAV77J | PAV77 | Nonserpentine | Phlomis purpurea | 72 | 0.06627 | 66.27 | 445.626 | 6.72440 |
| PAV77K | PAV77 | Nonserpentine | Phlomis purpurea | 91 | 0.12216 | 122.16 | 721.897 | 5.90944 |
| PAV77L | PAV77 | Nonserpentine | Phlomis purpurea | 76 | | | | |
| PAV77M | PAV77 | Nonserpentine | Phlomis purpurea | 57 | 0.10672 | 106.72 | 765.039 | 7.16866 |
| PAV77N | PAV77 | Nonserpentine | Phlomis purpurea | 146 | 0.10816 | 108.16 | 650.869 | 6.01765 |
| PAV77Ñ | PAV77 | Nonserpentine | Phlomis purpurea | 120 | 0.1362 | 136.2 | 920.813 | 6.76074 |
| PAV78A | PAV78 | Serpentine | Phlomis purpurea | 35 | | | | |
| PAV78B | PAV78 | Serpentine | Phlomis purpurea | 48 | 0.11953 | 119.53 | 917.725 | 7.67778 |
| PAV78C | PAV78 | Serpentine | Phlomis purpurea | 83 | 0.10518 | 105.18 | 1077.887 | 10.24802 |
| PAV78D | PAV78 | Serpentine | Phlomis purpurea | 64 | | | | |
| PAV78E | PAV78 | Serpentine | Phlomis purpurea | 36 | 0.11895 | 118.95 | 835.902 | 7.02734 |
| PAV78F | PAV78 | Serpentine | Phlomis purpurea | 33 | 0.09537 | 95.37 | 752.866 | 7.89416 |
| PAV78G | PAV78 | Serpentine | Phlomis purpurea | 61 | | | | |
| PAV78H | PAV78 | Serpentine | Phlomis purpurea | 68 | 0.10348 | 103.48 | 931.622 | 9.00292 |
| PAV78I | PAV78 | Serpentine | Phlomis purpurea | 46 | 0.17932 | 179.32 | 1372.580 | 7.65436 |
| PAV78J | PAV78 | Serpentine | Phlomis purpurea | 32 | 0.09195 | 91.95 | 841.913 | 9.15620 |
| PAV78K | PAV78 | Serpentine | Phlomis purpurea | 103 | | | | |
| PAV78L | PAV78 | Serpentine | Phlomis purpurea | 109 | 0.22688 | 226.88 | 1426.027 | 6.28538 |
| PAV78M | PAV78 | Serpentine | Phlomis purpurea | 48 | 0.06573 | 65.73 | 631.500 | 9.60749 |
| PAV78N | PAV78 | Serpentine | Phlomis purpurea | 57 | | | | |
| PAV78Ñ | PAV78 | Serpentine | Phlomis purpurea | 28 | 0.14393 | 143.93 | 913.373 | 6.34595 |
| PAV79A | PAV79 | Nonserpentine | Phlomis purpurea | 61 | 0.07601 | 76.01 | 305.559 | 4.01998 |
| PAV79B | PAV79 | Nonserpentine | Phlomis purpurea | 82 | 0.16511 | 165.11 | 983.216 | 5.95491 |
| PAV79C | PAV79 | Nonserpentine | Phlomis purpurea | 97 | 0.04866 | 48.66 | 274.210 | 5.63522 |
| PAV79D | PAV79 | Nonserpentine | Phlomis purpurea | 74 | 0.12346 | 123.46 | 700.919 | 5.67730 |
| PAV79E | PAV79 | Nonserpentine | Phlomis purpurea | 80 | 0.14015 | 140.15 | 729.094 | 5.20224 |
| PAV79F | PAV79 | Nonserpentine | Phlomis purpurea | 50 | 0.13578 | 135.78 | 948.061 | 6.98233 |
| PAV79G | PAV79 | Nonserpentine | Phlomis purpurea | 72 | | | | |
| PAV79H | PAV79 | Nonserpentine | Phlomis purpurea | 46 | | | | |
| PAV79I | PAV79 | Nonserpentine | Phlomis purpurea | 59 | 0.0413 | 41.3 | 555.631 | 13.45354 |
| PAV79J | PAV79 | Nonserpentine | Phlomis purpurea | 42 | | | | |
| PAV79K | PAV79 | Nonserpentine | Phlomis purpurea | 152 | 0.22108 | 221.08 | 1049.030 | 4.74502 |
| PAV79L | PAV79 | Nonserpentine | Phlomis purpurea | 39 | | | | |
| PAV79M | PAV79 | Nonserpentine | Phlomis purpurea | 141 | | | | |
| PAV79N | PAV79 | Nonserpentine | Phlomis purpurea | 26 | 0.15108 | 151.08 | 1808.927 | 11.97331 |
| PAV79Ñ | PAV79 | Nonserpentine | Phlomis purpurea | 105 | 0.15323 | 153.23 | 1825.409 | 11.91287 |
| PAV80A | PAV80 | Serpentine | Phlomis purpurea | 83 | 0.09505 | 95.05 | 883.611 | 9.29628 |
| PAV80B | PAV80 | Serpentine | Phlomis purpurea | 76 | 0.0807 | 80.7 | 693.824 | 8.59757 |
| PAV80C | PAV80 | Serpentine | Phlomis purpurea | 65 | | | | |
| PAV80D | PAV80 | Serpentine | Phlomis purpurea | 96 | 0.1104 | 110.4 | 881.148 | 7.98141 |
| PAV80E | PAV80 | Serpentine | Phlomis purpurea | 147 | 0.26457 | 264.57 | 1691.662 | 6.39401 |
| PAV80F | PAV80 | Serpentine | Phlomis purpurea | 39 | 0.06 | 60 | 447.257 | 7.45428 |
| PAV80G | PAV80 | Serpentine | Phlomis purpurea | 29 | 0.15007 | 150.07 | 1412.417 | 9.41172 |
| PAV80H | PAV80 | Serpentine | Phlomis purpurea | 55 | 0.08253 | 82.53 | 634.911 | 7.69309 |
| PAV80I | PAV80 | Serpentine | Phlomis purpurea | 57 | 0.11061 | 110.61 | 1006.909 | 9.10324 |
| PAV80J | PAV80 | Serpentine | Phlomis purpurea | 71 | 0.06802 | 68.02 | 548.801 | 8.06823 |
| PAV80K | PAV80 | Serpentine | Phlomis purpurea | 66 | | | | |
| PAV80L | PAV80 | Serpentine | Phlomis purpurea | 37 | 0.10766 | 107.66 | 991.059 | 9.20545 |
| PAV80M | PAV80 | Serpentine | Phlomis purpurea | 31 | 0.20998 | 209.98 | 1608.841 | 7.66188 |
| PAV80N | PAV80 | Serpentine | Phlomis purpurea | 110 | 0.14492 | 144.92 | 1531.478 | 10.56775 |
| PAV80Ñ | PAV80 | Serpentine | Phlomis purpurea | 41 | 0.09638 | 96.38 | 936.003 | 9.71159 |
| PAV82A | PAV82 | Serpentine | Phlomis purpurea | 55 | | | | |
| PAV82B | PAV82 | Serpentine | Phlomis purpurea | 47 | 0.0881 | 88.1 | 604.173 | 6.85781 |
| PAV82C | PAV82 | Serpentine | Phlomis purpurea | 52 | 0.05978 | 59.78 | 590.024 | 9.86992 |
| PAV82D | PAV82 | Serpentine | Phlomis purpurea | 48 | 0.1094 | 109.4 | 860.263 | 7.86346 |
| PAV82E | PAV82 | Serpentine | Phlomis purpurea | 74 | 0.08759 | 87.59 | 808.108 | 9.22603 |
| PAV82F | PAV82 | Serpentine | Phlomis purpurea | 62 | 0.07404 | 74.04 | 608.618 | 8.22012 |
| PAV82G | PAV82 | Serpentine | Phlomis purpurea | 61 | 0.05627 | 56.27 | 480.142 | 8.53282 |
| PAV82H | PAV82 | Serpentine | Phlomis purpurea | 59 | 0.09425 | 94.25 | 885.327 | 9.39339 |
| PAV82I | PAV82 | Serpentine | Phlomis purpurea | 79 | 0.08893 | 88.93 | 806.413 | 9.06795 |
| PAV82J | PAV82 | Serpentine | Phlomis purpurea | 43 | 0.12105 | 121.05 | 1049.950 | 8.67369 |
| PAV82K | PAV82 | Serpentine | Phlomis purpurea | 27 | 0.08357 | 83.57 | 788.121 | 9.43067 |
| PAV82L | PAV82 | Serpentine | Phlomis purpurea | 78 | 0.12285 | 122.85 | 884.724 | 7.20166 |
| PAV82M | PAV82 | Serpentine | Phlomis purpurea | 43 | 0.16977 | 169.77 | 1017.172 | 5.99147 |
| PAV82N | PAV82 | Serpentine | Phlomis purpurea | 42 | 0.09729 | 97.29 | 991.188 | 10.18797 |

| | | | | | | | | |
|---|---|---|---|---|---|---|---|---|
| PAV82Ñ | PAV82 | Serpentine | Phlomis purpurea | 69 | 0.1536 | 153.6 | 1121.043 | 7.29846 |
| PAV83A | PAV83 | Nonserpentine | Phlomis purpurea | 129 | 0.07823 | 78.23 | 887.762 | 11.34810 |
| PAV83B | PAV83 | Nonserpentine | Phlomis purpurea | 43 | 0.06468 | 64.68 | 1089.407 | 16.84303 |
| PAV83C | PAV83 | Nonserpentine | Phlomis purpurea | 19 | | | | |
| PAV83D | PAV83 | Nonserpentine | Phlomis purpurea | 49 | | | | |
| PAV83E | PAV83 | Nonserpentine | Phlomis purpurea | 47 | 0.12909 | 129.09 | 1198.090 | 9.28104 |
| PAV83F | PAV83 | Nonserpentine | Phlomis purpurea | 72 | 0.18954 | 189.54 | 2399.907 | 12.66174 |
| PAV83G | PAV83 | Nonserpentine | Phlomis purpurea | 86 | 0.11711 | 117.11 | 1081.830 | 9.23773 |
| PAV83H | PAV83 | Nonserpentine | Phlomis purpurea | 85 | 0.11322 | 113.22 | 1007.814 | 8.90138 |
| PAV83I | PAV83 | Nonserpentine | Phlomis purpurea | 87 | 0.08459 | 84.59 | 618.005 | 7.30589 |
| PAV83J | PAV83 | Nonserpentine | Phlomis purpurea | 38 | 0.10317 | 103.17 | 717.394 | 6.95351 |
| PAV83K | PAV83 | Nonserpentine | Phlomis purpurea | 114 | | | | |
| PAV83L | PAV83 | Nonserpentine | Phlomis purpurea | 58 | 0.09613 | 96.13 | 730.257 | 7.59656 |
| PAV83M | PAV83 | Nonserpentine | Phlomis purpurea | 89 | | | | |
| PAV83N | PAV83 | Nonserpentine | Phlomis purpurea | 67 | 0.06403 | 64.03 | 422.113 | 6.59243 |
| PAV83Ñ | PAV83 | Nonserpentine | Phlomis purpurea | 18 | 0.10156 | 101.56 | 859.071 | 8.45875 |
| PAV84A | PAV84 | Serpentine | Phlomis purpurea | 24 | | | | |
| PAV84B | PAV84 | Serpentine | Phlomis purpurea | 20 | | | | |
| PAV84C | PAV84 | Serpentine | Phlomis purpurea | 56 | | | | |
| PAV84D | PAV84 | Serpentine | Phlomis purpurea | 97 | 0.17149 | 171.49 | 1766.073 | 10.29840 |
| PAV84E | PAV84 | Serpentine | Phlomis purpurea | 117 | 0.1067 | 106.7 | 1040.965 | 9.75600 |
| PAV84F | PAV84 | Serpentine | Phlomis purpurea | 82 | 0.16531 | 165.31 | 1711.778 | 10.35496 |
| PAV84G | PAV84 | Serpentine | Phlomis purpurea | 101 | 0.16048 | 160.48 | 1784.516 | 11.11987 |
| PAV84H | PAV84 | Serpentine | Phlomis purpurea | 40 | 0.13854 | 138.54 | 1167.430 | 8.42666 |
| PAV84I | PAV84 | Serpentine | Phlomis purpurea | 48 | 0.08488 | 84.88 | 1341.949 | 15.80996 |
| PAV84J | PAV84 | Serpentine | Phlomis purpurea | 56 | 0.08997 | 89.97 | 932.110 | 10.36023 |
| PAV84K | PAV84 | Serpentine | Phlomis purpurea | 108 | 0.08581 | 85.81 | 891.238 | 10.38618 |
| PAV84L | PAV84 | Serpentine | Phlomis purpurea | 81 | 0.09962 | 99.62 | 1172.702 | 11.77175 |
| PAV84M | PAV84 | Serpentine | Phlomis purpurea | 82 | 0.09009 | 90.09 | 1049.957 | 11.65453 |
| PAV84N | PAV84 | Serpentine | Phlomis purpurea | 127 | 0.0697 | 69.7 | 1237.834 | 17.75945 |
| PAV84Ñ | PAV84 | Serpentine | Phlomis purpurea | 53 | 0.13868 | 138.68 | 1574.774 | 11.35545 |
| PAV85A | PAV85 | Nonserpentine | Phlomis purpurea | 121 | | | | |
| PAV85B | PAV85 | Nonserpentine | Phlomis purpurea | 111 | 0.1579 | 157.9 | 1431.456 | 9.06559 |
| PAV85C | PAV85 | Nonserpentine | Phlomis purpurea | 108 | 0.13059 | 130.59 | 1737.216 | 13.30283 |
| PAV85D | PAV85 | Nonserpentine | Phlomis purpurea | 53 | 0.1764 | 176.4 | 1705.487 | 9.66829 |
| PAV85E | PAV85 | Nonserpentine | Phlomis purpurea | 59 | | | | |
| PAV85F | PAV85 | Nonserpentine | Phlomis purpurea | 75 | 0.13558 | 135.58 | 1140.498 | 8.41199 |
| PAV85G | PAV85 | Nonserpentine | Phlomis purpurea | 188 | 0.15334 | 153.34 | 1333.920 | 8.69910 |
| PAV85H | PAV85 | Nonserpentine | Phlomis purpurea | 67 | | | | |
| PAV85I | PAV85 | Nonserpentine | Phlomis purpurea | 178 | | | | |
| PAV85J | PAV85 | Nonserpentine | Phlomis purpurea | 56 | 0.22192 | 221.92 | 1690.987 | 7.61980 |
| PAV85K | PAV85 | Nonserpentine | Phlomis purpurea | 60 | 0.1906 | 190.6 | 2592.552 | 13.60206 |
| PAV85L | PAV85 | Nonserpentine | Phlomis purpurea | 49 | 0.19209 | 192.09 | 2454.517 | 12.77795 |
| PAV85M | PAV85 | Nonserpentine | Phlomis purpurea | 233 | 0.11876 | 118.76 | 1098.449 | 9.24932 |
| PAV85N | PAV85 | Nonserpentine | Phlomis purpurea | 75 | | | | |
| PAV85Ñ | PAV85 | Nonserpentine | Phlomis purpurea | 128 | | | | |
| PAV86A | PAV86 | Nonserpentine | Phlomis purpurea | 72 | 0.03952 | 39.52 | 316.511 | 8.00888 |
| PAV86B | PAV86 | Nonserpentine | Phlomis purpurea | 50 | | | | |
| PAV86C | PAV86 | Nonserpentine | Phlomis purpurea | 91 | 0.09432 | 94.32 | 780.738 | 8.27754 |
| PAV86D | PAV86 | Nonserpentine | Phlomis purpurea | 37 | 0.19775 | 197.75 | 1224.878 | 6.19407 |
| PAV86E | PAV86 | Nonserpentine | Phlomis purpurea | 42 | 0.207 | 207 | 1316.863 | 6.36166 |
| PAV86F | PAV86 | Nonserpentine | Phlomis purpurea | 70 | 0.15266 | 152.66 | 734.372 | 4.81051 |
| PAV86G | PAV86 | Nonserpentine | Phlomis purpurea | 35 | | | | |
| PAV86H | PAV86 | Nonserpentine | Phlomis purpurea | 56 | 0.12445 | 124.45 | 555.085 | 4.46031 |
| PAV86I | PAV86 | Nonserpentine | Phlomis purpurea | 71 | 0.07268 | 72.68 | 518.816 | 7.13836 |
| PAV86J | PAV86 | Nonserpentine | Phlomis purpurea | 67 | 0.10522 | 105.22 | 424.770 | 4.03697 |
| PAV86K | PAV86 | Nonserpentine | Phlomis purpurea | 67 | 0.10106 | 101.06 | 538.466 | 5.32818 |
| PAV86L | PAV86 | Nonserpentine | Phlomis purpurea | 81 | 0.12828 | 128.28 | 775.373 | 6.04438 |
| PAV86M | PAV86 | Nonserpentine | Phlomis purpurea | 64 | 0.10175 | 101.75 | 436.491 | 4.28984 |
| PAV86N | PAV86 | Nonserpentine | Phlomis purpurea | 59 | 0.9889 | 988.9 | 445.131 | 0.45013 |
| PAV86Ñ | PAV86 | Nonserpentine | Phlomis purpurea | 21 | 0.11633 | 116.33 | 641.877 | 5.51773 |
| PAV88A | PAV88 | Serpentine | Phlomis purpurea | 36 | 0.11283 | 112.83 | 878.419 | 7.78533 |
| PAV88B | PAV88 | Serpentine | Phlomis purpurea | 28 | 0.07896 | 78.96 | 469.549 | 5.94667 |
| PAV88C | PAV88 | Serpentine | Phlomis purpurea | 94 | | | | |
| PAV88D | PAV88 | Serpentine | Phlomis purpurea | 109 | 0.12377 | 123.77 | 1239.385 | 10.01361 |
| PAV88E | PAV88 | Serpentine | Phlomis purpurea | 31 | 0.15902 | 159.02 | 947.853 | 5.96059 |
| PAV88F | PAV88 | Serpentine | Phlomis purpurea | 77 | 0.10067 | 100.67 | 569.010 | 5.65223 |
| PAV88G | PAV88 | Serpentine | Phlomis purpurea | 49 | | | | |
| PAV88H | PAV88 | Serpentine | Phlomis purpurea | 41 | 0.0484 | 48.4 | 351.968 | 7.27207 |
| PAV88I | PAV88 | Serpentine | Phlomis purpurea | 84 | | | | |
| PAV88J | PAV88 | Serpentine | Phlomis purpurea | 13 | | | | |
| PAV88K | PAV88 | Serpentine | Phlomis purpurea | 57 | | | | |
| PAV88L | PAV88 | Serpentine | Phlomis purpurea | 85 | | | | |
| PAV88M | PAV88 | Serpentine | Phlomis purpurea | 51 | | | | |
| PAV88N | PAV88 | Serpentine | Phlomis purpurea | 50 | | | | |
| PAV88Ñ | PAV88 | Serpentine | Phlomis purpurea | 54 | | | | |
